# A microRNA Signature of Metastatic Colorectal Cancer

**DOI:** 10.1101/2020.06.01.127647

**Authors:** Eirik Høye, Bastian Fromm, Paul Heinrich Michael Böttger, Diana Domanska, Annette Torgunrud, Christin Lund-Andersen, Torveig Weum Abrahamsen, Åsmund Avdem Fretland, Vegar Johansen Dagenborg, Susanne Lorenz, Bjørn Edwin, Eivind Hovig, Kjersti Flatmark

## Abstract

Although microRNAs (miRNA) are involved in all hallmarks of cancer, miRNA dysregulation in metastasis remains poorly understood and contradictory results have been published. The aim of this work was to identify miRNAs associated with metastatic progression of colorectal cancer (CRC). Novel and previously published next generation sequencing (NGS) datasets generated from 268 samples with primary (pCRC) and metastatic CRC (mCRC; liver, lung and peritoneal metastases) and tumor adjacent tissues were analyzed. Differential expression analysis was performed using a meticulous bioinformatics pipeline, including only bona fide miRNAs, utilizing miRNA-tailored quality control and processing, and applying a physiologically meaningful cut-off value (100 reads per million). The results were adjusted for host tissue background expression and samples from the different metastatic sites were independently analyzed. A metastatic signature containing five miRNAs up-regulated at multiple metastatic sites was identified (Mir-210_3p, Mir-191_5p, Mir-8-P1b_3p *(mir-141-3p)*, Mir-1307_5p, and Mir-155_5p) along with a number of miRNAs that were differentially expressed at individual metastatic sites. Several of these have previously been implicated in metastasis through involvement in epithelial-to-mesenchymal transition and hypoxia, while other identified miRNAs represent novel findings. The identified differentially expressed miRNAs confirm known associations and contribute novel insights into miRNA involvement in the metastatic process. The use of open science practices facilitates reproducibility, and new datasets may easily be added to the publicly available pipeline to continuously improve the knowledge in the field. The identified set of miRNAs provides a reliable starting-point for further research into the role of miRNAs in metastatic progression.

## Introduction

Colorectal cancer (CRC) is a heterogeneous disease and a leading cause of cancer-related deaths worldwide^1^, and metastatic progression to the liver, lungs and peritoneal surface remains the primary cause of CRC-related mortality. Metastasis is a complex process where cancer cells undergo adaptation to enable survival and establishment of tumors in organs with very different microenvironments^2,3^. The genomic and transcriptomic changes in metastatic CRC (mCRC) remain incompletely understood, particularly in the context of organ-specific metastasis^4–6^.

MicroRNAs (miRNAs) are evolutionary ancient post-transcriptional gene regulators that are involved in numerous biological processes and are molecular players in human disease, including cancer^7,8^. MiRNAs can be extracted from tissues and body fluids, and because of their remarkable chemical stability and the availability of sensitive detection methods, miRNAs have been suggested as cancer biomarkers^9–11^. However, to date, no miRNAs have been clinically implemented as biomarkers of CRC^5,12–18^. In mCRC, in particular, there is little consensus regarding which miRNAs are up- and down-regulated, limiting our understanding of their role in metastatic progression^5^.

The lack of consensus likely reflects some well-known caveats related to analysis of miRNAs in human disease. Non-miRNA sequences have been incorrectly annotated as miRNA genes^19^, and a bioinformatics workflow specifically tailored for analysis of miRNAs in bulk tissue samples has not been available. Furthermore, differential expression analysis has been performed without ensuring the presence of physiologically relevant tissue expression levels^20^. Also, accounting for differences in cellular composition is important, since many miRNAs are exclusively expressed in particular cell types or at specific developmental time points^21–24^, likely confounding analysis of bulk tissue samples. Finally, to elucidate the role of miRNAs in mCRC, failure to consider differences related to metastatic location, and differences in normal background expression, may have contributed to inconsistent results^12,25^.

To overcome these challenges, the publicly available, manually curated miRNA gene database MirGeneDB (mirgenedb.org), was used as miRNA reference^26^, and a novel bioinformatics pipeline was developed. The bioinformatics work-flow included use of the miRTrace software as a universal quality control pipeline specifically for miRNA next-generation sequencing (NGS) data^27^ with subsequent processing using miRge3.0^28^. A strict cut-off of 100 reads per million (RPM) was applied as the minimum expression level for physiological relevance^12^. Taking cell-type specific miRNA expression into account, metastatic samples from different sites were independently analyzed, which allowed correction for different background expression levels at the individual sites. Existing publicly available miRNA-sequencing (miRNA-seq) patient derived datasets, combined with novel miRNA-seq datasets from pCRC and mCRC with normal adjacent tissues, were analyzed, after quality control totaling 268 datasets. Using this unbiased analytical approach, a novel mCRC miRNA signature was identified, which partially overlaps with previous reports, but which also includes several miRNAs previously not identified in this context.

## Results

### NGS data collection and processing

New NGS datasets were successfully generated from 85 samples: pCRC (n=3), tumor adjacent colorectum (nCR; n=3), liver metastases (mLi; n=19), tumor adjacent liver (nLi; n=9), lung metastases (mLu; n=25), tumor adjacent lung (nLu; n=7), and peritoneal metastases (PM; n=20). From five studies, previously published datasets containing NGS analyses of miRNA in pCRC and mCRC were also included^12,14,15,29^ while data from four published studies were not accessible despite repeated requests^30–33^. In total, 350 NGS datasets were subjected to quality assessment using the miRTrace pipeline. The majority of the new NGS datasets and datasets from three previously published studies^12,15,34^ fulfilled the quality control criteria. Two studies were excluded, one because of low quality reads in the majority of samples^29^, the other because of contamination of reads from other organisms^14^ (Supplementary file 1). After quality control, a total of 268 NGS datasets remained for further analysis, including pCRC (n=120), nCR (n=25), mLi (n=35), nLi (n=20), mLu (n=28), nLu (n=10) and PM (n=30). These datasets were then successfully processed and mapped to MirGeneDB^26^ using miRge3.0^28^.

### Global miRNA expression

When analyzing global miRNA expression using the dimensionality reducing algorithm uniform manifold approximation and projection for dimension reduction (UMAP), the datasets clustered according to the tissue of origin (Fig. 1). The normal tissues (nLi, nLu and nCR) formed distinct clusters reflecting the unique transcriptional profiles of these organs. nLi and nCR clustered separately from the corresponding tumor tissues, while nLu clustered close to the mLu tissue datasets. In general, the pCRC and mCRC dataset clusters were less homogeneous than the normal tissue counterparts, but with exception of the PM datasets, which exhibited considerable overlap with the other malignant datasets, the pCRC, mLi, and mLu datasets all formed distinct clusters.

**Fig. 1.**
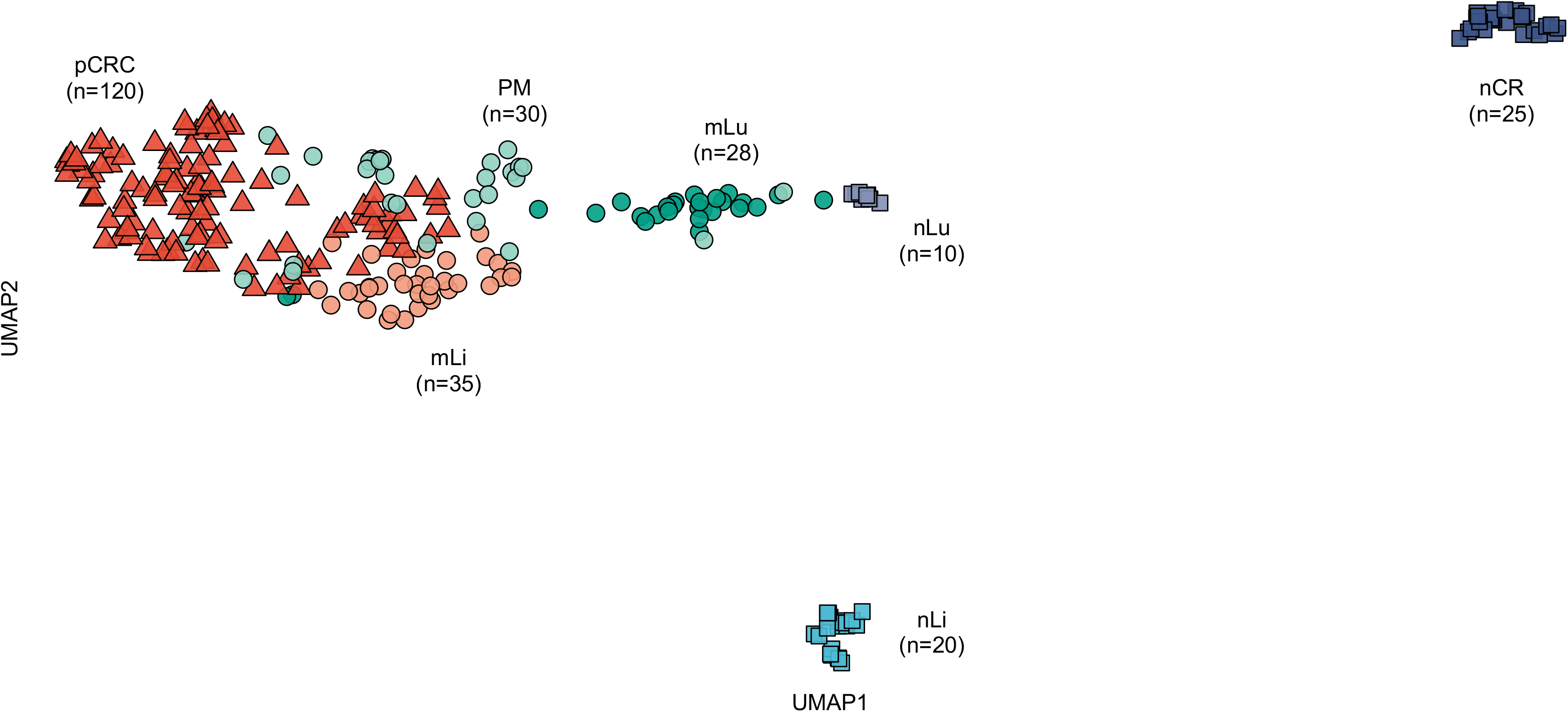
Global miRNA expression according to tissue of origin. UMAP cluster plot based on global miRNA expression, normalized by VST, from the DESeq2 bioconductor package, annotated by tissue of origin. Uniform approximation and projection for dimension reduction (UMAP); varianceStabilizingTransformation (VST); primary colorectal cancer (pCRC); normal colorectal tissue (nCR); CRC liver metastasis (mLi); normal adjacent liver tissue (nLi); CRC lung metastasis (mLu); normal adjacent lung tissue (nLu); CRC peritoneal metastasis (PM).

### Differential miRNA expression between pCRC and nCR

Thirty-two miRNAs were up-regulated and 35 miRNAs were down-regulated when comparing pCRC to nCR. Among the differentially expressed miRNAs were many well-known oncomiRs, including Mir-21_5p, multiple MIR-17 family members, Mir-31, Mir-221 (up-regulated) and Mir-8-P1b_3p (miR-141) (down-regulated). Several cell-type specific miRNAs were also detected at different levels in the two tissues, including higher levels of Mir-17-P1a/P1b_5p (mir-17; CD14+ monocytes) and Mir-223_3p (dendritic cells), and lower levels of Mir-486_5p and Mir-451_5P (red blood cells), the Mir-143_5p and Mir-145_5p (mesenchymal cells), Mir-150_5p (lymphocytes), Mir-375_3p and Mir-192-P1_5p (epithelial cells), and Mir-342_3p (dendritic cells, lymphocytes and macrophages)^22^. For a complete overview of differentially expressed miRNAs between pCRC and nCR, see Supplementary file 3.

### Differential expression analysis identifies miRNA signature of mCRC

After performing site specific differential expression analysis and correcting for background expression, a total of 26 miRNAs were identified as differentially expressed in one or more of the metastatic tissues compared to pCRC (Fig. 2 and Table 1). Two miRNAs, Mir-210_3p and Mir-191_5p, were up-regulated at all three metastatic sites. In addition, three miRNAs were up-regulated at two of the three sites; Mir-8-P1b_3p in mLi and mLu, Mir-1307_5p in mLi and PM, and Mir-155 in mLu and PM (Fig. 3). All these miRNAs were expressed well above the threshold for biological significance, with Mir-191_5p and Mir-8-P1b_3p being expressed at particularly high levels (>1000 RPM).

**Fig. 2.**
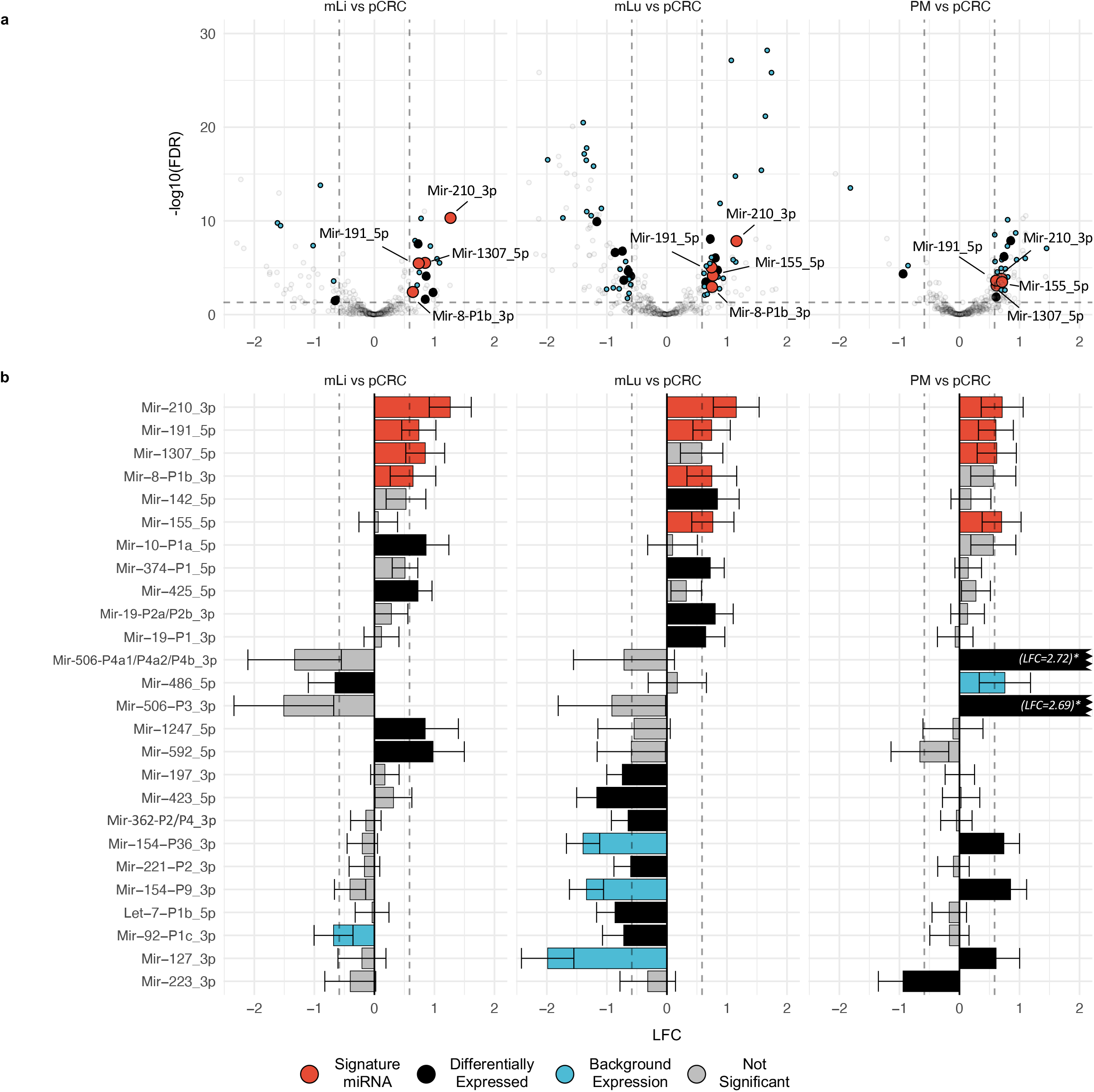
Differential expression analysis identifies miRNAs associated with metastatic colorectal cancer. **a** Volcano plots showing differentially expressed miRNAs in mLi, mLu and PM compared to pCRC. The horizontal axis shows LFC relative to pCRC, while the vertical axis shows-log10 FDR. The identified miRNAs were differentially expressed between pCRC and mCRC, with expression levels greater than 100 RPM in one of the tissues, and results were corrected for tumor adjacent background expression. **b** Bar plots of LFC of differentially expressed miRNAs in pCRC compared to the individual metastatic sites. Log2 fold change (LFC); false discovery rate (FDR); reads per million (RPM); primary colorectal cancer (pCRC); normal colorectal tissue (nCR); CRC liver metastasis (mLi); normal adjacent liver tissue (nLi); CRC lung metastasis (mLu); normal adjacent lung tissue (nLu); CRC peritoneal metastasis (PM).

**Table 1.** Differentially expressed miRNAs in mCRC compared to pCRC according to metastatic site.

| MirGeneDB ID (*miRBase ID*) | pCRC | mLi vs pCRC |  |  | mLu vs pCRC |  |  | PM vs pCRC |  |  |
| --- | --- | --- | --- | --- | --- | --- | --- | --- | --- | --- |
|  | RPM | RPM | LFC (SE) | FDR | RPM | LFC (SE) | FDR | RPM | LFC (SE) | FDR |
| Mir-191_5p | 13 630 | 23 253 | 0.74 (0.15) | 3.67E-06 | 26 989 | 0.74 (0.16) | 9.81E-06 | 20 429 | 0.61 (0.15) | 2.42E-04 |
| Mir-210_3p | 291 | 685 | 1.26 (0.18) | 4.73E-11 | 693 | 1.15 (0.19) | 1.55E-08 | 491 | 0.71 (0.18) | 3.37E-04 |
| Mir-1307_5p | 492 | 887 | 0.85 (0.17) | 2.99E-06 |  |  |  | 797 | 0.62 (0.17) | 8.29E-04 |
| Mir-8-P1b_3p (*mir-141-3p*) | 7518 | 11 957 | 0.64 (0.19) | 4.07E-03 | 15 111 | 0.75 (0.21) | 1.12E-03 |  |  |  |
| Mir-155_5p | 387 |  |  |  | 733 | 0.76 (0.18) | 7.24E-05 | 654 | 0.70 (0.17) | 1.59E-04 |
| Mir-10-P1a_5p (*mir-10a-5p*) | 97 123 | 164 925 | 0.86 (0.19) | 8.04E-05 |  |  |  |  |  |  |
| Mir-592_5p | 75 | 137 | 0.98 (0.27) | 4.88E-03 |  |  |  |  |  |  |
| Mir-1247_5p | 69 | 127 | 0.85 (0.28) | 2.58E-02 |  |  |  |  |  |  |
| Mir-425_5p | 700 | 1117 | 0.73 (0.12) | 2.90E-08 |  |  |  |  |  |  |
| Mir-486_5p | 2013 | 1489 | -0.66 (0.23) | 3.47E-02 |  |  |  |  |  |  |
| Mir-142_5p | 3453 |  |  |  | 7117 | 0.84 (0.18) | 2.01E-05 |  |  |  |
| Mir-19-P2a/P2b_3p | 1297 |  |  |  | 2345 | 0.80 (0.15) | 9.38E-07 |  |  |  |
| Mir-374-P1_5p | 115 |  |  |  | 217 | 0.72 (0.12) | 9.07E-09 |  |  |  |
| Mir-19-P1_3p | 379 |  |  |  | 597 | 0.65 (0.16) | 3.42E-04 |  |  |  |
| Mir-423_5p | 433 |  |  |  | 202 | -1.17 (0.17) | 1.35E-10 |  |  |  |
| Let-7-P1b_5p | 1057 |  |  |  | 650 | -0.86 (0.16) | 2.55E-07 |  |  |  |
| Mir-197_3p | 181 |  |  |  | 113 | -0.74 (0.13) | 1.76E-07 |  |  |  |
| Mir-92-P1c_3p | 1398 |  |  |  | 981 | -0.72 (0.18) | 2.22E-04 |  |  |  |
| Mir-362-P2/P4_3p | 441 |  |  |  | 302 | -0.65 (0.14) | 2.36E-05 |  |  |  |
| Mir-221-P2_3p | 1633 |  |  |  | 1101 | -0.60 (0.14) | 8.50E-05 |  |  |  |
| Mir-506-P3_3p | 6 |  |  |  |  |  |  | 112 | 2.69 (0.29) | 6.35E-09 |
| Mir-506 P4a1/P4a2/P4b_3p | 14 |  |  |  |  |  |  | 191 | 2.72 (0.29) | 5.75E-11 |
| Mir-154-P9_3p | 253 |  |  |  |  |  |  | 455 | 0.85 (0.14) | 1.43E-08 |
| Mir-127_3p | 2263 |  |  |  |  |  |  | 2888 | 0.61 (0.20) | 1.42E-02 |
| Mir-154-P36_3p | 195 |  |  |  |  |  |  | 330 | 0.74 (0.13) | 7.13E-07 |
| Mir-223_3p | 619 |  |  |  |  |  |  | 280 | -0.94 (0.21) | 4.62E-05 |

Reads per million (RPM); log 2 fold change (LFC) estimated using DESeq2; standard error (SE); primary colorectal cancer (pCRC); normal colorectal tissue (nCR); CRC liver metastasis (mLi); normal adjacent liver tissue (nLi); CRC lung metastasis (mLu); normal adjacent lung tissue (nLu); CRC peritoneal metastasis (PM); false discovery rate (FDR)

**Fig. 3.**
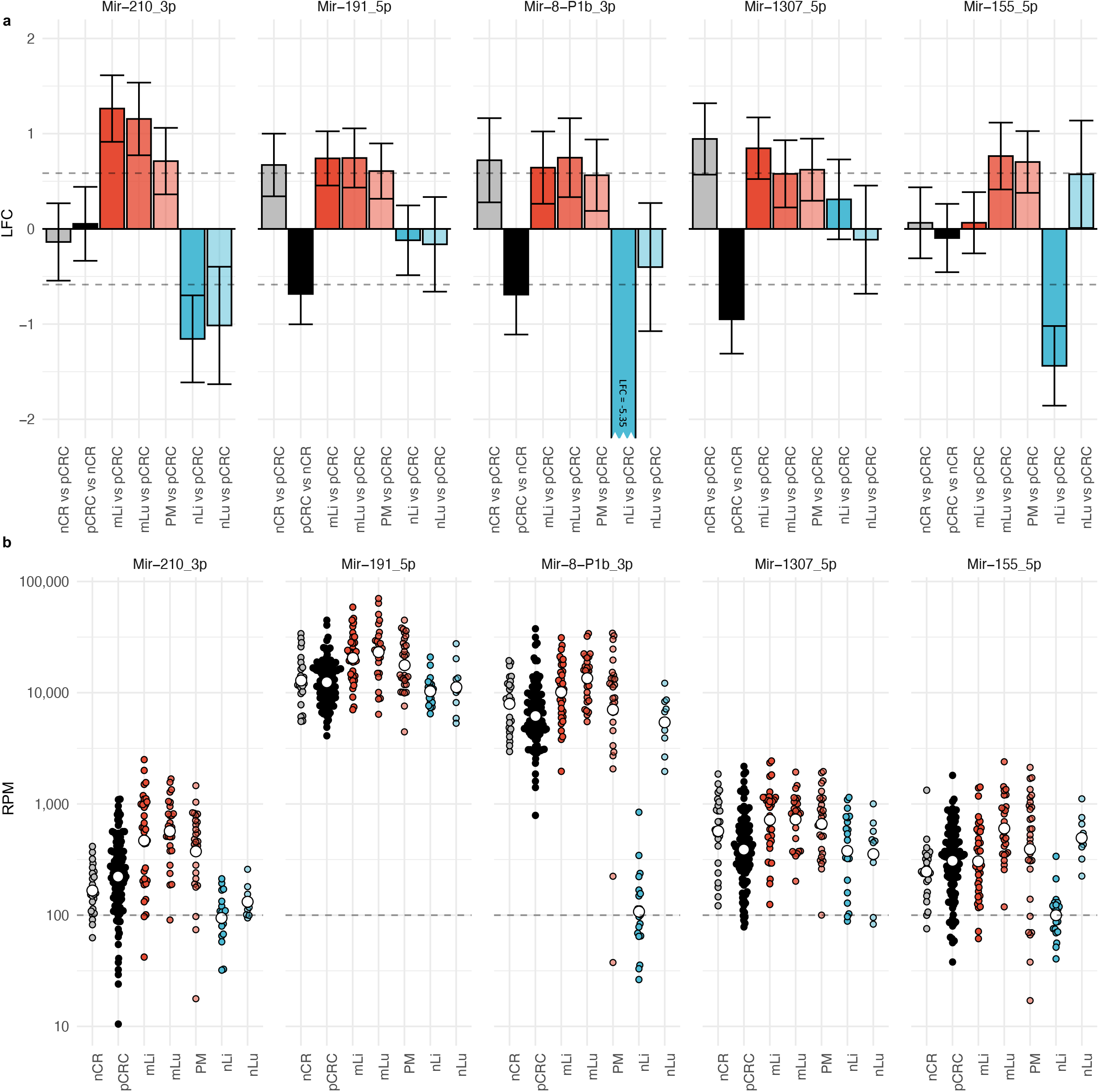
Differential expression analysis identifies a five-gene miRNA signature up-regulated at multiple metastatic sites. **a** Bar plots showing LFC of the five signature miRNAs that were differentially expressed between pCRC and at least two mCRC sites. Mir-210_3p and Mir-191_5p were up-regulated at all three metastatic sites, while, Mir-8-P1b_3p was up-regulated in mLi and mLu, Mir-1307_5p in mLi and PM, and Mir-155 in mLu and PM, after correcting for multiple hypothesis testing. Error bars represent 95% confidence intervals from DESeq2 estimated SE of the LFC. **b** Scatter plots of Log10 RPM in each tissue. Log2 fold change (LFC); reads per million (RPM); standard error (SE) primary colorectal cancer (pCRC); normal colorectal tissue (nCR); CRC liver metastasis (mLi); normal adjacent liver tissue (nLi); CRC lung metastasis (mLu); normal adjacent lung tissue (nLu); CRC peritoneal metastasis (PM).

In addition, several miRNAs were up- or down-regulated at individual metastatic sites. In mLi, four miRNAs were identified as up-regulated, Mir-10-P1a_5p, Mir-592_5p, Mir-1247_5p and Mir-425_5p, while Mir-486_5p was down-regulated. Of these, Mir-10-P1a_5p was expressed at exceptionally high levels, with 97 123 RPM in pCRC and 164 925 RPM in mLi. In mLu Mir-19-P1_3p, Mir-19-P2a/P2b_3p, Mir-374-P1_5p and Mir-142_5p were up-regulated while Mir-423_5p, Let-7-P1b_5p, Mir-197_3p, Mir-92-P1c_3p, Mir-362-P2/P4_3p and Mir-221_3p were down-regulated. In PM Mir-506-P3_3p, Mir-506-P4a1/P4a2/P4b_3p, Mir-154-P9_3p and Mir-127_3p, Mir-154-P36_3p were up-regulated and Mir-223_3p were down-regulated.

### qPCR validation

qPCR analysis of randomly selected mLi (n=11) and pCRC (n=11) samples not included in the previous NGS analysis replicated the findings from the NGS data, validating up-regulation of Mir-210_3p in mLi compared to pCRC. Welch two-sided t-test on dCq values, t-statistic = −2.25, degrees of freedom = 19.95, p-value = 0.036.

### Analysis of cell-type specific miRNAs

Of the 45 previously validated cell-type specific miRNAs, only 25 had a mean expression greater than 100 RPM in at least one of the tissues and formed the basis for the following analysis. The relative expression levels of cell-type specific miRNAs showed clear differences between the tissues, particularly prominent for the tumor adjacent tissues (Fig. 4A). For instance, hepatocyte specific Mir-122_5p was detected at high and moderate levels in nLi and mLi, respectively, and at extremely low levels in the other tissues. Also, Mir-143_3p and Mir-145_5p was detected at higher levels in nCR and nLu relative to the other datasets (p=4.38E-02 and p=3.42E-04, respectively). The epithelial cell specific Mir-8-P2a_3p and Mir-8-P2b_3p, were detected at higher levels in the intestinal epithelial-derived tissues compared to nLi and nLu (p=2.20E-16 and p=2.24E-16, respectively). In line with these findings, principal component analysis (PCA) (Fig. 4B) showed distinct clusters for nLi and nLu, whereas nCR clustered closer to pCRC and the metastatic tissues. The correlation circle (Fig. 4C) shows the loading of the PCA, indicating the direction and relative contribution of the 15 cell-type specific miRNAs that contributed most to the clustering of each tissue.

**Fig. 4.**
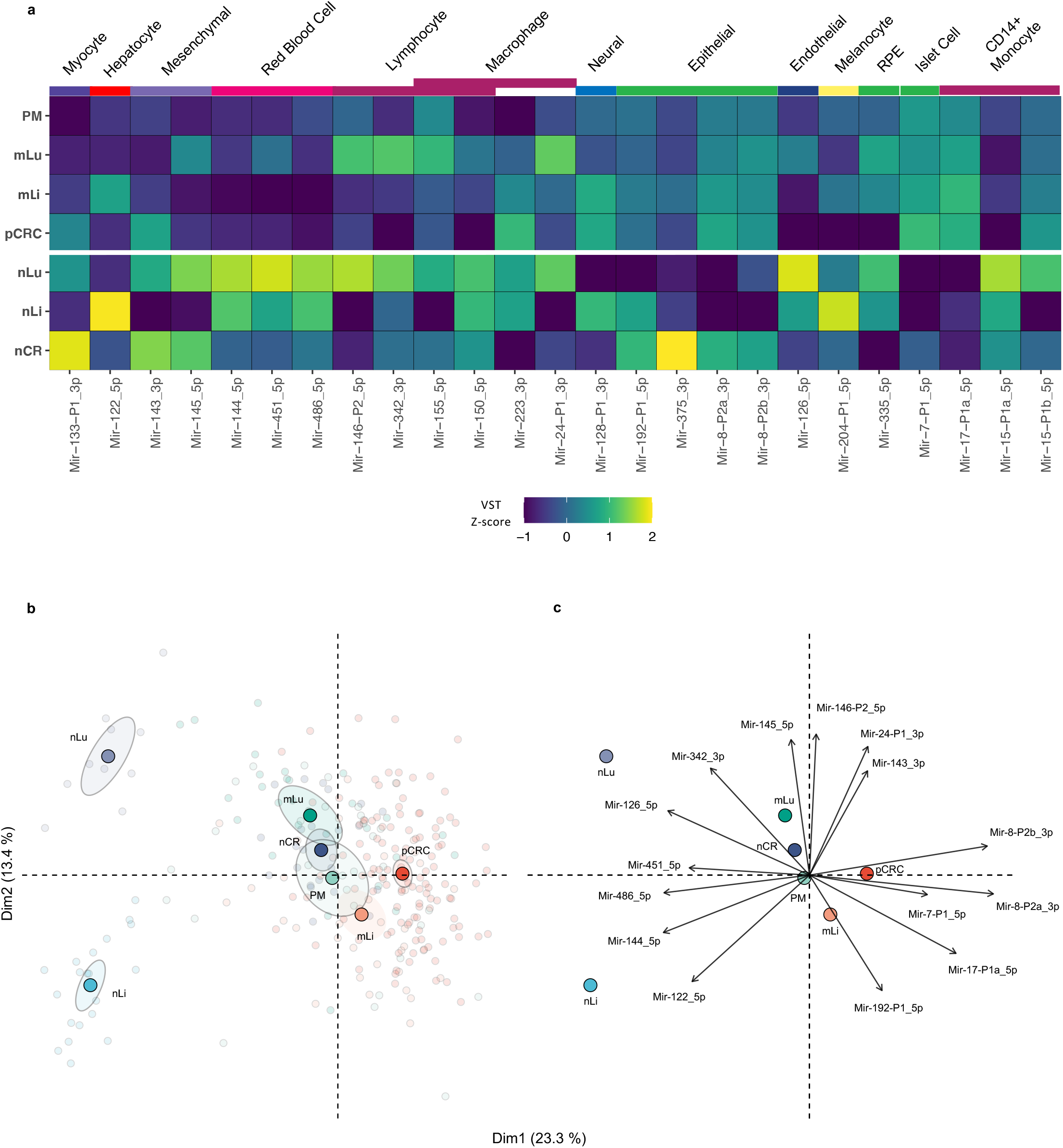
Cell-type specific miRNAs provide insight into the cellular composition of tissues. **a** Heatmap illustrating the expression of 25 cell-specific miRNAs for each tissue that had mean expression > 100 RPM in at least one tissue. The color scale indicates the z-score of RPM for each miRNA. **b** PCA plot based on analysis of 25 cell-specific miRNA with expression>100 reads per million, VST normalized, colored by tissue. Transparent points are individual samples, solid points represent the statistical mean and shaded ellipses is 95% confidence of the mean of each tissue. **c** The correlation circle shows PCA loadings, which are the direction and relative contribution the top 15 cell-type specific miRNAs had on clustering of each tissue. Reads per million (RPM); principal component analysis (PCA); varianceStabilizingTransformation (VST); primary colorectal cancer (pCRC); normal colorectal tissue (nCR); CRC liver metastasis (mLi); normal adjacent liver tissue (nLi); CRC lung metastasis (mLu); normal adjacent lung tissue (nLu); CRC peritoneal metastasis (PM).

### Gene set enrichment (GSE) analysis

RBiomirGS analysis revealed multiple GO terms and KEGG pathways that were predicted to be either more or less repressed because of differential expression of miRNAs between pCRC and mCRC. Among terms predicted to be less repressed, several were common to mLi and mLu. These included the five Molecular Function GO terms: Transcription Coregulator Activity, Transcription Coactivator Activity, Transcription Regulator Activity, mRNA Binding and Poly Purine Tract Binding. Also estimated to be less repressed were the eight Cellular Component GO terms, including Transcription Repressor Complex, Caveola, Transferase Complex, Golgi Aparatus, Glutamatergic Synapse, Catalytic Complex, Nuclear Envelope and Transcription Regulator Complex. Furthermore, 92 Biological Process GO terms, many of which are related to transcriptional activity, were also estimated to have reduced miRNA repression in both mLi and mLu. Among the KEGG pathways, only mLu had pathways predicted to be less repressed, including Long Term Potentiation, Axon Guidance, Long Term Depression, Ubiquitin Mediated Proteolysis, Neurotrophin Signaling Pathway, TGF Beta Signaling Pathway, Adherens Junction and Phosphatidylinositol Signaling System. One KEGG pathway, Systemic Lupus Erythematosus, was estimated to be more repressed in both mLi and mLu, while the Parkinsons Disease KEGG pathway was repressed only in mLu. Notably, no differentially repressed GO terms or KEGG pathways were shared between the PM datasets and the other two sites. There were also many GO terms and KEGG pathways that were differentially repressed specifically to individual sites. All results for the GSE are shown in Supplementary file 6.

## Discussion

The initial differential expression analysis comparing pCRC and nCR revealed 68 differentially expressed miRNAs, reflecting major and genome-wide changes of malignant transformation. Among the up-regulated miRNAs, a subset representing previously reported “oncoMirs” was identified, including Mir-21, and members of the three clusters Mir-17-92, Mir-31, and Mir-221. These miRNAs have validated mRNA targets in CRC, such as phosphatase and tensin homolog (PTEN), transforming growth factor beta receptor (TGFBR), and Smad^16,35,36^. A number of other miRNAs previously connected to CRC were also up-regulated, such as MIR-15 family member Mir-29-P2a/P2b (*miR-29b*), two MIR-96 family members, Mir-135-P3, Mir-130-P2a (*miR-301a*)^37,38^, Mir-181-P1c^39^, and Mir-224^16,35^. One up-regulated miRNA, Mir-95-P2 (*miR-421*), has not been reported for CRC previously, but its deregulation has been demonstrated in other cancers^40–42^. The remaining down-regulated miRNAs were all previously reported to be associated with CRC; for instance, the tumor suppressor MIR-10 family (4 genes), MIR-15, MIR-192 and MIR-194^16,35^ have been implicated in CRC progression. The identification of miRNAs previously reported to be deregulated in CRC and associated with relevant signaling pathways provides confidence in our analytical approach.

Using the same analytical strategy, a metastatic signature was identified, which included five miRNAs that were differentially expressed in at least two metastatic sites compared to pCRC. Of these, Mir-210_3p and Mir-191_5p were up-regulated at all three metastatic sites, which would suggest a strong metastasis-related significance. Mir-210_3p is in the literature known as the “hypoxamiR” because of its key involvement in the cellular response to hypoxia. The Mir-210_3p promoter has a binding site for hypoxia inducible transcription factors HIF-1*α* and HIF-2*α*^43^ which coordinate cellular responses to hypoxic stress, including regulation of key pathways in metastasis, such as angiogenesis, cell proliferation, differentiation and apoptosis^44^. Increased Mir-210_3p expression, which was additionally validated by qPCR in a separate set of samples, therefore suggests hypoxic stress to be a common feature of CRC metastasis, irrespective of metastatic site. In contrast, the role of Mir-191_5p in metastasis is less well established, and this miRNA is therefore an interesting candidate for further follow-up studies. Additionally, three miRNAs, Mir-8-P1b_3p, Mir-1307_5p, and Mir-155_5p, were up-regulated at two of three metastatic sites. Mir-8-P1b_3p (*miR-141*) is a member of the MIR-8 (*miR-200*) family, which is strongly involved in epithelial to mesenchymal transition (EMT) and targets transcription factors ZEB1 and ZEB2, which in turn suppress E-cadherin expression^45,46^. In our data, Mir-8-P1b_3p was down-regulated in pCRC relative to nCR, while up-regulated in mLi and mLu (Fig. 3). This fits well with the concept that tumors undergo EMT as part of tumorigenesis at the primary site, while in the established metastasis the inverse process, mesenchymal to epithelial transition, is necessary to establish growth in the new metastatic microenvironment. Mir-155_5p, which was preferentially expressed in mLu and PM, is reported to be specifically expressed in lymphocytes and macrophages^22^, possibly indicating higher abundance of these cell types in the metastases compared to pCRC. Less is known about the biological activity of Mir-1307_5p, although reports have suggested a role in lung adenocarcinoma proliferation^47^ and as a predictor of hepatocellular carcinoma metastasis^48^. The identification of this miRNA as highly expressed in mCRC points to its involvement in metastasis, and Mir-1307_5p therefore represents another target for further studies. Several of the signature miRNAs have previously been implicated in the metastatic process, providing evidence that their observed up-regulation represents adaptations to survival metastatic site.

To further investigate potential biological implications of miRNA expression, GSE analyses were performed to explore predicted effects on mRNA expression based on miRNAs that were differentially expressed at the metastatic sites. Through these analyses, GO terms related to transcription were estimated to be less repressed at the metastatic sites, and KEGG pathways “Axon Guidance”, “Long-Term Potentiation” and “Long-Term Depression” were estimated to be less repressed in mLu, possibly representing adaptations to challenges in this microenvironment. However, the biology of mRNA repression by miRNAs is complex. The seed sequence of any miRNA can usually target a large number of mRNAs, and an mRNA can potentially be targeted for repression by multiple miRNAs^5^. The functional role of these miRNAs will therefore be dependent on the local cellular contexture, and because the net effect on mRNA expression is highly unpredictable, GSE analysis cannot replace experimental validation. Furthermore, a miRNA may potentially have abnormal expression in multiple diseases, such as Mir-21, which has been shown to be up-regulated in 29 different diseases in addition to CRC^49^. Therefore, the functional role of the differentially expressed miRNAs is still not clear, but this work still represents an excellent starting point for further studies, as well as a basis for interpreting the, often contradictory, existing literature.

In order to obtain results that are physiologically relevant, the absolute expression level of a miRNA is an important consideration. A cut-off level of 100 RPM was therefore applied as a minimum expression level to be included in these analyses^20^. Several of the differentially expressed miRNAs were expressed at very high absolute levels, for instance with Mir-191_5p, Mir-8-P1b_3p and Mir-10-P1a_5p all being expressed at >1000 RPM in both pCRC and mCRC. Expression levels of this magnitude strongly suggest that alterations in expression of these miRNAs will influence target mRNA levels. Mir-10-P1a_5p is of particular interest, being highly and differentially expressed in mLi compared to pCRC (164 925 RPM and 97 123 RPM, respectively), suggesting specific adaptations to the unique conditions of this microenvironment. Previous experimental evidence suggests that the Mir-10-P1a_5p paralogue, Mir-10-P1b_5p, is associated with enhanced metastatic capability by down-regulation of metastasis suppressor Hoxd10^50–52^. Since Mir-10-P1a_5p and Mir-10-P1b_5p share the same seed sequence (ACCCUGU), the range of mRNA targets and functional roles would be expected to be similar.

In clinical cancer studies, the most commonly available material will be bulk tissue samples, and a tumor biopsy will therefore contain a variable amount of cancer cells of epithelial origin together with a mixture of cell types from the “host tissue” (such as fibroblasts, endothelial cells, myocytes, blood cells and immune cells). The importance of keeping cell specificity in mind when interpreting miRNA expression levels is illustrated by the high levels of hepatocyte specific Mir-122_5p in liver metastases compared to pCRC, which is likely due to the presence of hepatocytes in the metastatic tissues, and do not indicate that this is a metastasis biomarker. Similarly, Mir-143_3p/Mir-145_5p were previously suggested to be tumor suppressors due to apparent “down-regulation” in pCRC relative to nCR^53^. In our datasets, both miRNAs were detected at much lower levels in pCRC relative to nCR, but since these miRNAs are exclusively expressed in mesenchymal cells, the low expression in pCRC can only be explained by mesenchymal cells being less frequently present in the colorectal tumors than in the normal colorectal wall^54,55^. Expression of known cell-type specific miRNAs could also inform on the relative composition of cell types in the different tissues, and as expected, the three tumor adjacent tissues analyzed clustered separately and had distinct miRNA expression patterns. For instance, the epithelial specific miRNAs, such as Mir-8-P2a_3p (miR-200b) and Mir-8-P2b_3p (miR-200c), had similar levels in the malignant tissues, which is consistent with the intestinal epithelial origin of the cancer cells. This means that considering background expression was a necessary step to ensure that cell composition effects would not confound the differential expression analysis, and failure to do so would likely have led to incorrect identification of miRNAs not associated with metastasis. Taken together, when analyzing bulk tissue samples, it is therefore necessary to consider the composition of the “host tissue”, which in this study was handled by defining and correcting for background expression and by analyzing tumor samples from different metastatic sites as separate entities.

In addition to applying a 100 RPM cut-off level, correcting for background expression, and analyzing metastatic locations as separate entities, a number of other methodological improvements were implemented in this work. A systematic effort was made to ensure the quality of the analyses and making the data and the analytical pipeline transparent and available to other researchers in the interest of reproducibility. A novel quality assessment step was introduced in the analysis, using the recently developed miRTrace algorithm, resulting in the exclusion of two previous studies^14,29^. Furthermore, our recently established database MirGeneDB, containing only *bona fide* miRNA genes, was used as a reference for mapping miRNA reads to the human genome, representing a further level of precision compared to the previously more commonly used repository, miRBase^56^. Regretfully, miRBase has been shown to contain a proportion of incorrect miRNA annotations^19^, and including such sequences in the analyses could potentially lead to weakening the power of statistical analyses and result in misleading conclusions. Furthermore, to facilitate reproducibility, a common set of bioinformatics tools were used, and raw NGS datasets from all studies were run through an identical pipeline, from postprocessing and read alignment to miRNA gene counting, using the latest version of miRge^28^, a bioinformatics pipeline specifically developed with miRNAs in mind. Combining all these measures and specialized tools, the resulting set of differentially expressed miRNAs between pCRC and mCRC therefore highly likely represents a reliable mCRC signature.

In summary, this work represents the most comprehensive attempt to characterize miRNA expression in mCRC. An unbiased, stringent and transparent bioinformatics approach was developed and applied to a large compilation of new and previously published NGS datasets to identify miRNAs associated with metastatic progression in CRC. Comparison of pCRC and nCR replicated many previous findings of up- and down-regulation of well-known oncomiRs and tumor-suppressor miRNAs, supporting the analytical strategy. Correction for background expression was performed, and tumor samples from different metastatic sites were analyzed as separate entities using a 100 RPM expression cut-off level to ensure biological relevance. A metastatic signature containing five miRNAs that were up-regulated at multiple metastatic sites was identified along with a number of miRNAs differentially expressed at individual metastatic sites. Many of these miRNAs have previously been implicated as key players in the metastatic process, while for others, the involvement in tumor cell adaptations at the distant site represent novel findings. The use of open science practices and the biological relevance of the findings lend confidence in the resulting mCRC signature, which provides a starting-point for further elucidation of the role of miRNAs in metastatic progression. The pipeline developed for this analysis is freely available to other researchers to expand on the results presented in this work, as well for exploring other cancer entities and disease settings. This work could therefore represent a first step to generate a miRNA tissue expression “atlas” for researchers interested in miRNA biomarkers, for instance to identify miRNAs that are specifically deregulated in a disease of interest.

## Materials and Methods

### Patient samples

Tissue samples were obtained from study specific biobanks: pCRC and nCR samples were from the LARC-EX study (NCT02113384); mLi and nLi samples from the OSLO-COMET trial (NCT01516710); mLu and nLu samples from our lung metastasis biobank (S-06402b) and PM samples from the Peritoneal Surface Malignancies biobank (NCT02073500). The studies were approved by the Regional Ethics Committee of South-East Norway, and patients were included following written informed consent. Patient samples were collected at the time of surgery and were snap frozen in liquid nitrogen at the time of collection and stored at −80°C. Samples were prepared and processed as described in^57^.

### RNA extraction and NGS

RNA was extracted using Qiagen Allprep DNA/RNA/miRNA universal kit, which simultaneously isolates genomic DNA and total RNA. RNA concentration was evaluated using a NanoDrop spectrophotometer (ThermoFisher, Waltham, Massachusetts, USA) and RNA integrity was evaluated using the Bioanalyzer RNA 6000 Nano kit (Agilent Technologies, Santa Clara, California, USA). MiRNA NGS libraries were then prepared using TruSeq Small RNA Library protocol and sequences using HiSeq 2500 High Throughput Sequencer (all from Illumina, San Diego, California, USA).

### Identification of published NGS datasets

A literature search was conducted for the terms “microRNA + CRC + next generation sequencing” in different variations, and reviews were studied^16,35^. Publicly available datasets were downloaded from European Genome-phenome Archive (EGA), the Sequence Read Archive (SRA) and the Gene Expression Omnibus (GEO)^12–15,34^.

### Data processing, read alignment and gene counting

All datasets were processed using the same pipeline. miRTrace^27^ was used for preprocessing and quality control (QC) of raw data (FASTQ files). Briefly, low-quality reads, defined as reads where less than 50% of nucleotides had a Phread quality score greater than 20 were discarded. 3p adapter sequences were trimmed, and reads made up of repetitive elements and reads shorter than 18nt were removed. After miRTrace QC, samples were excluded if < 25% of reads were between 20 and 25 nt, if > 75% of reads were discarded, or if < 10% of reads were identified as miRNA. If > 50% of the datasets in a study failed the QC criteria, or if significant contamination was detected, it was excluded. After QC, raw data processing, read alignment and gene counting was performed using miRge3.0^28^, with MirGeneDB2.0^26^ as reference. To account for cross mapping of miRNA genes, miRge3.0 merges miRNA genes with very similar sequences into one annotation, reducing 537 human miRNA annotations in MirGeneDB2.0 to a total of 389 unique annotations.

### Analysis of global miRNA expression

For data visualization of global miRNA expression, read counts were normalized using the variance Stabilizing Transformation() (VST) function from the DESeq2 package^58^. UMAP was used to visualize the similarity of datasets on the global miRNA expression level. The UMAP algorithm reduces the 389 dimensions of unique miRNA genes into two dimensions for visualization^59^. The umap R package was used, and datasets were annotated by tissue.

### Differential expression analysis

The differential expression analysis can be viewed in the supplementary R-markdown file, (Supplementary file 3).

Differential expression analysis was performed using DESeq2 (version 1.26)^58^, which estimates LFC and its standard error (SE), using raw, non-normalized read counts as input. Hypothesis testing was performed by a Wald test against the null hypothesis LFC = 0, followed by the Benjamini-Hochberg procedure to correct for multiple hypothesis testing, applying a false discovery rate (FDR) threshold of < 0.05. The LFC shrinkage function in DESeq2, lfcShrink(), was enabled, to shrink fold changes for miRNAs with higher variance. Differentially expressed miRNAs were filtered for relevance by requiring an LFC > 0.58 or LFC < −0.58, and also requiring that at least one of the compared tissues had a mean expression > 100 RPM, a level previously suggested as minimal cut-off for physiological activity^20^. MiRNAs known to be cell-type specific^22^ were also labelled for each miRNA. Information regarding normal tissue background expression levels was obtained by analyzing differential expression between nCR versus nLi and nLu datasets, and pCRC versus nLi and nLu datasets. Then, in the mCRC versus pCRC differential expression analysis, miRNAs differentially expressed in the same direction in the corresponding normal tissue were not considered differentially expressed in the respective metastatic site. For the PM tissue datasets, where no tumor adjacent tissue was available, the union of nLi and nLu background expression was used.

### qPCR validation

Eleven additional randomly chosen mLi and 11 pCRC tissue samples were selected for qPCR validation of increased expression of Mir-210_3p in mLi compared to pCRC. Synthetic RNA Spike-Ins UniSp2, UniSp4 and UniSp5 (Qiagen, Düsseldorf, Germany Cat. No. 339390) were added pre-isolation. RT-PCR was done with miRCURY LNA RT Kit (Qiagen Cat. No. 339340), adding UniSp6 and cel-miR-39-3p RNA Spike-Ins (Qiagen Cat. No 339390). qPCR was done using Qiagen miRCURY SYBR Green Kit (Qiagen Cat. No. 339345) and miRCURY LNA miRNA PCR Assays (Qiagen Cat. No. 339306). Mir-103 (Qiagen primer Cat. No. YP00204306) was used as reference miRNA, while the Mir-210_3p primer was Qiagen Cat. No. YP00204333. Two PCR replicates were run per primer assay. Mir-210_3p Cq values were normalized to the reference miRNA, to obtain the dCq value. Welch two-sided t-test was used to compare the dCq values, to test if the expression levels were different between mLi and pCRC. (Supplementary file 5).

### Analysis of cell-type specific miRNAs

Two analyses were performed to assess differences in expression levels of 45 previously validated cell-type specific miRNAs^22^, and thereby infer differences in cell composition in the tissues. A heatmap using z-scores of RPM values per miRNA was made to illustrate the relative expression of each miRNA in the tissues (Fig. 4A). Principle component analysis (PCA) plots were made with the FactoMineR PCA() function to illustrate which miRNAs were more or less prevalent in each tissue. Welch two-sided t-test comparing mean VST values between two groups of tissues was done to assess if the observed differences in cell-type specific miRNA levels were statistically significant (Supplementary file 4).

### Gene set enrichment analysis

Gene set enrichment analysis (GSEA) was performed using RBiomirGS^60^ on GO Molecular Function, Cellular Component and Biological Process, and KEGG pathways, downloaded from https://www.gsea-msigdb.org/gsea/msigdb/collections.jsp. miRNA to mRNA predicted interactions were combined with data from the miRNA differential expression analysis (DESeq2 estimated LFC and FDR values for each mCRC site versus pCRC). The input values to RBiomirGS were corrected for normal background expression, reducing LFC towards 0 and increasing FDR towards 1, depending on the magnitude of the normal background. All miRNAs were then given an S_miRNA_ score -log10 p-value * sign(log_2_FC)., and for each mRNA, a S_mRNA_ score was calculated by summing up the S_miRNA_ scores of all the predicted miRNA to mRNA interactions. The S_mRNA_ scores were then used to perform logistic regression, which provides likelihoods of gene sets being more or less suppressed due to differential expression of miRNAs.

## Acknowledgements

The authors thank Michael Hackenberg for help with data acquisition and Marc Halushka for discussions. This work was supported by the South-Eastern Norway Regional Health Authority [grant numbers 2014041 to KF, 2018014 to EH], the Norwegian Cancer Society [grant number 4499184 to CLA] and the Research Council of Norway [grant number 218325].

## Competing interests

The authors have declared no competing interests.

## Availability

Data from current and previous studies can be found at EGA accession number EGAS00001001127 (14), GEO accession numbers: GSE57381 (31), GSE46622 (15) and GSE63119 (16), and SRA accession PRJNA397121 for datasets prepared in this study and (17). Bioinformatics pipeline can be found at (https://github.com/eirikhoye/mirna_pipeline)

## Supplementary material

Supplementary file 1: miRTrace QC reports (.html)

Supplementary file 2: miRge3.0 count matrix and sample metadata (.csv)

Supplementary file 3: R-markdown: DESeq2 differential expression analysis (.html)

Supplementary file 4: R-markdown: code for figures (.html)

Supplementary file 5: R-markdown qPCR analysis (.html)

Supplementary file 6: R-markdown GSE results (.html)

## Notes

https://github.com/eirikhoye/mirna_pipeline

### Summary of Updates

Major overhaul of manuscript following sequencing of 40 new smallRNA-Seq datasets, editing for clarity, and re-designed figures for improved visualisation.

## References

1 Ferlay J et al. Cancer incidence and mortality worldwide: sources, methods and major patterns in GLOBOCAN 2012. Int J Cancer 2015; 136: E359–86.

2 Nguyen DX, Bos PD, Massagué J. Metastasis: from dissemination to organ-specific colonization. Nat Rev Cancer 2009; 9: 274–284.

3 Riihimäki M, Hemminki A, Sundquist J, Hemminki K. Patterns of metastasis in colon and rectal cancer. Sci Rep 2016; 6: 29765.

4 Massague J, Obenauf AC. Metastatic colonization by circulating tumour cells. Nature 2016; 529: 298–306.

5 Flatmark K, Høye E, Fromm B. microRNAs as cancer biomarkers. Scandinavian Journal of Clinical and Laboratory Investigation. 2016; 76: S80–S83.

6 Arnold M, Sierra MS, Laversanne M, Soerjomataram I, Jemal A, Bray F. Global patterns and trends in colorectal cancer incidence and mortality. Gut 2017; 66: 683–691.

7 Bartel DP. Metazoan MicroRNAs. Cell 2018; 173: 20–51.

8 Dalmay T, Edwards DR. MicroRNAs and the hallmarks of cancer. Oncogene 2006; 25: 6170–6175.

9 Jung M et al. Robust microRNA stability in degraded RNA preparations from human tissue and cell samples. Clin Chem 2010; 56: 998–1006.

10 Volinia S et al. A microRNA expression signature of human solid tumors defines cancer gene targets. Proc Natl Acad Sci U S A 2006; 103: 2257–2261.

11 Baffa R et al. MicroRNA expression profiling of human metastatic cancers identifies cancer gene targets. J Pathol 2009; 219: 214–221.

12 Neerincx M et al. MiR expression profiles of paired primary colorectal cancer and metastases by next-generation sequencing. Oncogenesis 2015; 4: e170.

13 Röhr C et al. High-throughput miRNA and mRNA sequencing of paired colorectal normal, tumor and metastasis tissues and bioinformatic modeling of miRNA-1 therapeutic applications. PLoS One 2013; 8: e67461.

14 Goossens-Beumer IJ et al. MicroRNA classifier and nomogram for metastasis prediction in colon cancer. Cancer Epidemiol Biomarkers Prev 2015; 24: 187–197.

15 Schee K et al. Deep Sequencing the MicroRNA Transcriptome in Colorectal Cancer. PLoS ONE. 2013; 8: e66165.

16 Cekaite L, Eide PW, Lind GE, Skotheim RI, Lothe RA. MicroRNAs as growth regulators, their function and biomarker status in colorectal cancer. Oncotarget 2016; 7: 6476–6505.

17 To KK, Tong CW, Wu M, Cho WC. MicroRNAs in the prognosis and therapy of colorectal cancer: From bench to bedside. World J Gastroenterol 2018; 24: 2949–2973.

18 Git A et al. Systematic comparison of microarray profiling, real-time PCR, and next-generation sequencing technologies for measuring differential microRNA expression. RNA 2010; 16: 991–1006.

19 Fromm B et al. A Uniform System for the Annotation of Vertebrate microRNA Genes and the Evolution of the Human microRNAome. Annu Rev Genet 2015; 49: 213–242.

20 Mullokandov G et al. High-throughput assessment of microRNA activity and function using microRNA sensor and decoy libraries. Nat Methods 2012; 9: 840–846.

21 Witwer KW, Halushka MK. Toward the promise of microRNAs – Enhancing reproducibility and rigor in microRNA research. RNA Biol 2016; 13: 1103–1116.

22 McCall MN et al. Toward the human cellular microRNAome. Genome Res 2017; 27: 1769–1781.

23 Rie D de et al. An integrated expression atlas of miRNAs and their promoters in human and mouse. Nat Biotechnol 2017. doi:10.1038/nbt.3947.

24 Juzenas S et al. A comprehensive, cell specific microRNA catalogue of human peripheral blood. Nucleic Acids Research. 2017; 45: 9290–9301.

25 Mudduluru G et al. A Systematic Approach to Defining the microRNA Landscape in Metastasis. Cancer Res 2015; 75: 3010–3019.

26 Fromm B et al. MirGeneDB 2.0: the metazoan microRNA complement. Nucleic Acids Res 2020. doi:10.1093/nar/gkz885.

27 Kang W, Eldfjell Y, Fromm B, Estivill X, Biryukova I, Friedländer MR. miRTrace reveals the organismal origins of microRNA sequencing data. Genome Biol 2018; 19: 213.

28 Patil AH, Halushka MK. miRge3.0: a comprehensive microRNA and tRF sequencing analysis pipeline. Cold Spring Harbor Laboratory. 2021;: 2021.01.18.427129.

29 Rohr C et al. High-throughput miRNA and mRNA sequencing of paired colorectal normal, tumor and metastasis tissues and bioinformatic modeling of miRNA-1 therapeutic applications. PLoS One 2013; 8: e67461.

30 Sun Y, Wang L, Guo S-C, Wu X-B, Xu X-H. High-throughput sequencing to identify miRNA biomarkers in colorectal cancer patients. Oncology Letters. 2014; 8: 711–713.

31 Sun G, Cheng Y-W, Lai L, Huang T-C, Wang J, Wu X et al. Signature miRNAs in colorectal cancers were revealed using a bias reduction small RNA deep sequencing protocol. Oncotarget 2016; 7: 3857–3872.

32 Liang G et al. Deep sequencing reveals complex mechanisms of microRNA deregulation in colorectal cancer. Int J Oncol 2014; 45: 603–610.

33 Hamfjord J et al. Differential expression of miRNAs in colorectal cancer: comparison of paired tumor tissue and adjacent normal mucosa using high-throughput sequencing. PLoS One 2012; 7: e34150.

34 Selitsky SR et al. Transcriptomic Analysis of Chronic Hepatitis B and C and Liver Cancer Reveals MicroRNA-Mediated Control of Cholesterol Synthesis Programs. mBio. 2015; 6. doi:10.1128/mbio.01500-15.

35 Strubberg AM, Madison BB. MicroRNAs in the etiology of colorectal cancer: pathways and clinical implications. Dis Model Mech 2017; 10: 197–214.

36 Schee K, Boye K, Abrahamsen TW, Fodstad O, Flatmark K. Clinical relevance of microRNA miR-21, miR-31, miR-92a, miR-101, miR-106a and miR-145 in colorectal cancer. BMC Cancer 2012; 12: 505.

37 Zhang W et al. MicroRNA-301a promotes migration and invasion by targeting TGFBR2 in human colorectal cancer. J Exp Clin Cancer Res 2014; 33: 113.

38 Ma X et al. Modulation of tumorigenesis by the pro-inflammatory microRNA miR-301a in mouse models of lung cancer and colorectal cancer. Cell Discov 2015; 1: 15005.

39 Guo X et al. miR-181d and c-myc-mediated inhibition of CRY2 and FBXL3 reprograms metabolism in colorectal cancer. Cell Death Dis 2017; 8: e2958.

40 Scaravilli M et al. MiR-1247-5p is overexpressed in castration resistant prostate cancer and targets MYCBP2. The Prostate. 2015; 75: 798–805.

41 Chen L, Tang Y, Wang J, Yan Z, Xu R. miR-421 induces cell proliferation and apoptosis resistance in human nasopharyngeal carcinoma via downregulation of FOXO4. Biochem Biophys Res Commun 2013; 435: 745–750.

42 Zhao J et al. MicroRNA-7: a promising new target in cancer therapy. Cancer Cell Int 2015; 15: 103.

43 Chan YC, Banerjee J, Choi SY, Sen CK. miR-210: the master hypoxamir. Microcirculation 2012; 19: 215–223.

44 Rankin EB, Giaccia AJ. Hypoxic control of metastasis. Science 2016; 352: 175–180.

45 Pencheva N, Tavazoie SF. Control of metastatic progression by microRNA regulatory networks. Nat Cell Biol 2013; 15: 546–554.

46 Gregory PA et al. The miR-200 family and miR-205 regulate epithelial to mesenchymal transition by targeting ZEB1 and SIP1. Nat Cell Biol 2008; 10: 593–601.

47 Du X et al. MiR-1307-5p targeting TRAF3 upregulates the MAPK/NF-κB pathway and promotes lung adenocarcinoma proliferation. Cancer Cell Int 2020; 20: 502.

48 Eun JW et al. Circulating Exosomal MicroRNA-1307-5p as a Predictor for Metastasis in Patients with Hepatocellular Carcinoma. Cancers 2020; 12. doi:10.3390/cancers12123819.

49 Jenike AE, Halushka MK. miR-21: a non-specific biomarker of all maladies. Biomark Res 2021; 9: 18.

50 Ma L, Reinhardt F, Pan E, Soutschek J, Bhat B, Marcusson EG et al. Therapeutic silencing of miR-10b inhibits metastasis in a mouse mammary tumor model. Nat Biotechnol 2010; 28: 341–347.

51 Ma L, Teruya-Feldstein J, Weinberg RA. Tumour invasion and metastasis initiated by microRNA-10b in breast cancer. Nature 2007; 449: 682–688.

52 Wang Y, Li Z, Zhao X, Zuo X, Peng Z. miR-10b promotes invasion by targeting HOXD10 in colorectal cancer. Oncol Lett 2016; 12: 488–494.

53 Michael MZ, O’ Connor SM, van Holst Pellekaan NG, Young GP, James RJ. Reduced accumulation of specific microRNAs in colorectal neoplasia. Mol Cancer Res 2003; 1: 882–891.

54 Chivukula RR et al. An essential mesenchymal function for miR-143/145 in intestinal epithelial regeneration. Cell 2014; 157: 1104–1116.

55 Kent OA, McCall MN, Cornish TC, Halushka MK. Lessons from miR-143/145: the importance of cell-type localization of miRNAs. Nucleic Acids Res 2014; 42: 7528–7538.

56 Griffiths-Jones S, Grocock RJ, van Dongen S, Bateman A, Enright AJ. miRBase: microRNA sequences, targets and gene nomenclature. Nucleic Acids Res 2006; 34: D140–4.

57 Østrup O et al. Molecular signatures reflecting microenvironmental metabolism and chemotherapy-induced immunogenic cell death in colorectal liver metastases. Oncotarget. 2017; 8. doi:10.18632/oncotarget.19350.

58 Love MI, Huber W, Anders S. Moderated estimation of fold change and dispersion for RNA-seq data with DESeq2. Genome Biology. 2014; 15. doi:10.1186/s13059-014-0550-8.

59 McInnes L, Healy J, Melville J. UMAP: Uniform Manifold Approximation and Projection for Dimension Reduction. arXiv [stat.ML]. 2018.http://arxiv.org/abs/1802.03426.

60 Zhang J, Storey KB. RBiomirGS: an all-in-one miRNA gene set analysis solution featuring target mRNA mapping and expression profile integration. PeerJ 2018; 6: e4262.

