## Supplementary material for "A microRNA Signature of Metastatic Colorectal Cancer": Supplementary_file_3_DESeq2_differential_expression_analysis.html

comet\_analysis\_report\_automated\_no\_lfcthreshold\_08.12.20\_res\_alpha\_0.05\_with\_LFC\_shrinkage\_LFC\_0.58\_mirge3\_5.0.utf8


```
# import libraries
library("tidyverse")
library("DESeq2")
library("knitr")
library('kableExtra')
library("RColorBrewer")
library("pheatmap")
#library("scatterplot3d")
#library("gplots")
library("dplyr")
#library("rBLAST")
#library("Biostrings")
library("BiocParallel")
library("broman")
library("gridExtra")
library("ggpubr")
library("FactoMineR")
library("factoextra")
```

```
register(MulticoreParam(12))
```

```
"
Define threshold for signature miRNA, including effect size, 
significance therhold, and expression size
"
```

```
## [1] "\nDefine threshold for signature miRNA, including effect size, \nsignificance therhold, and expression size\n"
```

```
lfc.Threshold <- 0.5849625
rpm.Threshold <- 100
p.Threshold   <- 0.05
```

### Import MirGeneDB metadata

```
MirGeneDB_info <- read_delim('/Users/eirikhoy/Dropbox/projects/comet_analysis/data/hsa_MirGeneDB_to_miRBase.csv', delim = ';')
```

```
## Parsed with column specification:
## cols(
##   MirGeneDB_ID = col_character(),
##   MiRBase_ID = col_character(),
##   Family = col_character(),
##   Seed = col_character(),
##   `5p accession` = col_character(),
##   `3p accession` = col_character(),
##   Chromosome = col_character(),
##   Start = col_double(),
##   End = col_double(),
##   Strand = col_character(),
##   `Node of origin (locus)` = col_character(),
##   `Node of origin (family)` = col_character(),
##   `3' NTU` = col_character(),
##   ` UG ` = col_double(),
##   UGUG = col_double(),
##   CNNC = col_double()
## )
```

```
MirGeneDB_info <- MirGeneDB_info %>% filter(!grepl("-v[2-9]", MirGeneDB_ID)) # keep only -v1
MirGeneDB_info$MirGeneDB_ID <- str_replace_all(MirGeneDB_info$MirGeneDB_ID, "-v1", "")
```

### Functions

```
DeseqObject <- function(DESIGN, countdata, coldata, consensus="None", sample_type="None", Ref) {
  "
  Function to create DESeq2 object
  "
  
  dds <- DESeqDataSetFromMatrix(countData = countdata,
                                colData = coldata,
                                design = as.formula(paste("~", DESIGN)))
    # Kick out non-consensus samples
  if (!(consensus == "None")) {
    dds <- dds[, dds$paper %in% consensus]
  }

  # Kick out samples that are not bulk tissue
  if (!(sample_type == "None")) {
    dds <- dds[, dds$sample_type == sample_type]
  }

  dds$tissue.type <- relevel(dds$tissue.type, ref=ref)
  dds$tissue.type <- droplevels(dds$tissue.type)
  
  dds <- DESeq(dds,
               parallel=TRUE,
               BPPARAM=MulticoreParam(3)
               )

  return(dds)
}
```

```
DeseqResult <- function(dds, column, coef, tissue_type_A, tissue_type_B, 
                        lfc.Threshold, rpm.Threshold,
                        norm_adj_up       = "None",
                        norm_adj_down     = "None",
                        pCRC_adj_up   = "None",
                        pCRC_adj_down = "None"){
  "
  Function to return results from DESeq2 for different conditions, 
  including control for normal adjacent tissue, if available
  "
    p.threshold   <- p.Threshold
    lfc.threshold <- lfc.Threshold
    rpm.threshold <- rpm.Threshold
    samples_tissue_type_A <- colData(dds)[, column] == tissue_type_A
    samples_tissue_type_B <- colData(dds)[, column] == tissue_type_B
  res <- results(dds, name = coef, alpha = p.threshold)
  res <- lfcShrink(dds, coef=coef, res=res)
    rpm <- t(t(counts(dds)) / colSums(counts(dds))) * 1000000
    sig <- rownames(res[(abs(res$log2FoldChange) > lfc.threshold) &
                        (res$padj < p.threshold) &
                        !is.na(res$padj), ])
    sig <- sig[ (rowMeans(rpm[sig, samples_tissue_type_A]) > rpm.threshold) |
                (rowMeans(rpm[sig, samples_tissue_type_B]) > rpm.threshold)
                ]
    res_sig    <- res[sig, ]
    up_mirna   <- rownames(res_sig[res_sig$log2FoldChange > lfc.Threshold, ])
    down_mirna <- rownames(res_sig[res_sig$log2FoldChange < -lfc.Threshold, ])
  
    if (!(norm_adj_up == "None")) {
      up_mirna <- setdiff(up_mirna, norm_adj_up)
    }
    if (!(norm_adj_down == "None")) {
      down_mirna <- setdiff(down_mirna, norm_adj_down)
    }
    if (!(pCRC_adj_up == "None")) {
      up_mirna <- setdiff(up_mirna, pCRC_adj_up)
    }
    if (!(pCRC_adj_down == "None")) {
      down_mirna <- setdiff(down_mirna, pCRC_adj_down)
    }
        
    return_list <- list("rpm" = rpm, "res" = res, "sig"=sig, "res_sig"=res_sig, 
                        "down_mirna"=down_mirna, "up_mirna"=up_mirna)
    
    return(return_list)

}
```

```
SigList <- function(res, dds, tissue_type_A, tissue_type_B, coef,
                            norm_adj_up, norm_adj_down, 
                            pCRC_adj_up, pCRC_adj_down){
  "
  Function to create annotated lists of signature miRNA
  Return will print upregulated or downregulated miRNA, 
  by printing <signature_list>$up_mirna
  or          <signature_list>$down_mirna
  "
  group_A_rpm <- rowMeans(res$rpm[res$sig, dds$tissue.type == tissue_type_A])
  group_A_rpm_std <- rowSds(res$rpm[res$sig, dds$tissue.type == tissue_type_A])
  group_B_rpm <- rowMeans(res$rpm[res$sig, dds$tissue.type == tissue_type_B])
  group_B_rpm_std <- rowSds(res$rpm[res$sig, dds$tissue.type == tissue_type_B])
  lfc.deseq2  <- res$res[res$sig, ]$log2FoldChange
  lfcSE.deseq2<- res$res[res$sig, ]$lfcSE
  neg.log.10.adj.p <- format(res$res[res$sig, ]$padj, digits=3)
  signature_mirna <- res$sig
  sig_list <- dplyr::tibble(signature_mirna, lfc.deseq2, lfcSE.deseq2,
                            group_A_rpm, #group_A_rpm_std,
                            group_B_rpm, #group_B_rpm_std,
                            neg.log.10.adj.p)
  sig_list$signature_sub <- str_replace_all(signature_mirna, "/.*", "") %>% str_replace_all(., c("_5p" = "", "_3p" = ""))
  sig_list <- left_join(sig_list, MirGeneDB_info, by=c("signature_sub" = "MirGeneDB_ID"))
  
  # create list of upregulated mirna
  up_mirna <- sig_list %>%
    filter(lfc.deseq2 > lfc.Threshold) %>% 
    
    # Annotate which miRNA are cell markers
    mutate(
      cell_marker = ifelse(signature_mirna %in% names(cell_spec_dict_inv), cell_spec_dict_inv[signature_mirna], '')) %>%
    mutate(
      cell_marker = cell_spec(cell_marker, color = ifelse(cell_marker != '', 'white', 'black'),
                              background = ifelse(cell_marker != '', 'blue', 'white'),
                              bold = ifelse(cell_marker != '', F, F)))

    # Annotate which miRNA are in normal_adjacent
    if (norm_adj_up != "None") {
      up_mirna <- up_mirna %>%
        mutate(
          norm_adj = ifelse(signature_mirna %in% norm_adj_up, 'yes', '')) %>%
        mutate(
          norm_adj = cell_spec(norm_adj, color = ifelse(norm_adj == 'yes', 'white', 'black'),
                               background = ifelse(norm_adj == 'yes', 'black', 'white'),
                               bold = ifelse(norm_adj == 'yes', F, F))
        )
    }
    else up_mirna$norm_adj <- "na"
    
  # Annotate which miRNA are in pCRC_adjacent
    if (pCRC_adj_up != "None") {
      up_mirna <- up_mirna %>%
        mutate(
          pCRC_adj = ifelse(signature_mirna %in% pCRC_adj_up, 'yes', '')) %>%
        mutate(
          pCRC_adj = cell_spec(pCRC_adj, color = ifelse(pCRC_adj == 'yes', 'white', 'black'),
                               background = ifelse(pCRC_adj == 'yes', 'black', 'white'),
                               bold = ifelse(pCRC_adj == 'yes', F, F))
        )
    }
    else up_mirna$pCRC_adj <- "na"

    # number of upregulated miRNA
    number_upregulated <- dim(up_mirna)[1]
    
    # select only relevant rows
    up_mirna <- up_mirna %>% select(signature_mirna, lfc.deseq2, lfcSE.deseq2, neg.log.10.adj.p, 
                                    group_A_rpm,  #group_A_rpm_std,
                                    group_B_rpm, #group_B_rpm_std,
                                    MiRBase_ID, Family, Seed, Chromosome,
                                    cell_marker, norm_adj, pCRC_adj)
    
    
  # Create kable list with annotations
    up_mirna <- up_mirna %>%
      arrange(-lfc.deseq2) %>%
      arrange(desc(cell_marker)) %>%
      arrange(pCRC_adj) %>%
      arrange(norm_adj) %>%
      kable(col.names = c("miRNA", "LFC", "lfcSE", "FDR", 
                          paste('RPM', tissue_type_A), #paste('std', tissue_type_A), 
                          paste('RPM', tissue_type_B), #paste('std', tissue_type_B), 
                          "miRBase_ID", "Family", "Seed", "Chr",
                          "Cell-Type Specific", 'Norm Background', 'pCRC Background'),
            escape = F, booktabs = F, caption = paste("Upregulated in ", coef),
            digits = c(0, 2, 2, 3, 0, 0, 2, 3, 0, 0, 0, 0, 0, 0, 0)) %>%
      kable_styling(bootstrap_options = c("striped", "hover", "condensed"), full_width = T, 
                  fixed_thead = list(enabled = T)) %>%
      scroll_box(width = "2000px")
    
 
  # create list of downregulated miRNA
  down_mirna <- sig_list %>%
    filter(lfc.deseq2 < -lfc.Threshold) %>% 
    
    # Annotate which miRNA are cell markers
    mutate(
      cell_marker = ifelse(signature_mirna %in% names(cell_spec_dict_inv), cell_spec_dict_inv[signature_mirna], '')) %>%
    mutate(
      cell_marker = cell_spec(cell_marker, color = ifelse(cell_marker != '', 'white', 'black'),
                              background = ifelse(cell_marker != '', 'blue', 'white'),
                              bold = ifelse(cell_marker != '', F, F)))

    # Annotate which miRNA are in normal_adjacent
    if (norm_adj_down != "None") {
      down_mirna <- down_mirna %>%
        mutate(
          norm_adj = ifelse(signature_mirna %in% norm_adj_down, 'yes', '')) %>%
        mutate(
          norm_adj = cell_spec(norm_adj, color = ifelse(norm_adj == 'yes', 'white', 'black'),
                               background = ifelse(norm_adj == 'yes', 'black', 'white'),
                               bold = ifelse(norm_adj == 'yes', F, F))
        )
    }
    else down_mirna$norm_adj <- "na"

    # Annotate which miRNA are in pCRC_adjacent
    if (pCRC_adj_down != "None") {
      down_mirna <- down_mirna %>%
        mutate(
          pCRC_adj = ifelse(signature_mirna %in% pCRC_adj_down, 'yes', '')) %>%
        mutate(
          pCRC_adj = cell_spec(pCRC_adj, color = ifelse(pCRC_adj == 'yes', 'white', 'black'),
                               background = ifelse(pCRC_adj == 'yes', 'black', 'white'),
                               bold = ifelse(pCRC_adj == 'yes', F, F))
        )
    }
    else down_mirna$pCRC_adj <- "na"
  
    # number of upregulated miRNA
    number_downregulated <- dim(down_mirna)[1]

    down_mirna <- down_mirna %>% select(signature_mirna, lfc.deseq2, lfcSE.deseq2, neg.log.10.adj.p,
                                        group_A_rpm,  #group_A_rpm_std,
                                        group_B_rpm, #group_B_rpm_std,
                                        MiRBase_ID, Family, Seed, Chromosome,
                                        cell_marker, norm_adj, pCRC_adj)    
    
  # Create kable list with annotations    
    down_mirna <- down_mirna %>%
      arrange(lfc.deseq2) %>%
      arrange(desc(cell_marker)) %>%
      arrange(pCRC_adj) %>%
      arrange(norm_adj) %>%
      kable(col.names = c("miRNA", "LFC", "lfcSE", "FDR",
                          paste('RPM', tissue_type_A), #paste('std', tissue_type_A), 
                          paste('RPM', tissue_type_B), #paste('std', tissue_type_B), 
                          "miRBase_ID", "Family", "Seed", "Chr",
                           "Cell-Type Specific", 'Norm Background', 'pCRC Background'),
            escape = F, booktabs = F, caption = paste("Downregulated in ", coef),
            digits = c(0, 2, 2, 3, 0, 0, 2, 3, 0, 0, 0, 0, 0, 0, 0)) %>%
      kable_styling(bootstrap_options = c("striped", "hover", "condensed"), full_width = T, 
                  fixed_thead = list(enabled = T)) %>%
       scroll_box(width = "2000px")

  # Function return is to print kable, either upregulated or downregulated miRNA
  return_list = list("up_mirna" = up_mirna, "down_mirna" = down_mirna, 
                     "number_upregulated" = number_upregulated, 
                     "number_downregulated" = number_downregulated)
  return(return_list)
}
```

```
# Read the sample information into a data frame
sampleinfo <- read_delim("/Users/eirikhoy/Dropbox/projects/comet_analysis/data/sample_info_v9.csv", delim=';')
```

```
## Parsed with column specification:
## cols(
##   filename = col_character(),
##   paper = col_character(),
##   sample_name = col_character(),
##   type.tissue = col_character(),
##   type = col_character(),
##   tissue = col_character(),
##   paper_sample_name = col_character(),
##   `3p-adapter` = col_character(),
##   qc_report = col_character(),
##   malignant = col_character(),
##   new_old = col_character()
## )
```

```
sampleinfo <- sampleinfo %>%
  filter(qc_report == 'keep')
sampleinfo$filename <- str_remove(sampleinfo$filename, '.fasta.fas.gz.bam')
sampleinfo$filename <- str_remove(sampleinfo$filename, '.fasta.bam')
sampleinfo$filename <- str_replace_all(sampleinfo$filename, pattern = '\\.', replacement = '_')
sampleinfo$filename <- str_replace_all(sampleinfo$filename, pattern = '-', replacement = '_')
#sampleinfo$filename <- str_replace_all(sampleinfo$filename, pattern = '__', replacement = '_')

# Read the data into R
#seqdata <- read_delim("/Users/eirikhoy/Dropbox/projects/comet_analysis/data/count_matrix_08.12.20.csv", delim = ';')
seqdata_1 <- read_delim("/Users/eirikhoy/Dropbox/projects/mirge3/output_dir/miRge.2021-01-19_10-03-25/miR.Counts.csv", delim = ',')
```

```
## Parsed with column specification:
## cols(
##   .default = col_double(),
##   miRNA = col_character()
## )
```

```
## See spec(...) for full column specifications.
```

```
#seqdata_2 <- read_delim("/Users/eirikhoy/Dropbox/projects/mirge3/output_dir/miRge.2021-01-19_12-04-26/miR.Counts.csv", delim = ',')
seqdata_3 <- read_delim("/Users/eirikhoy/Dropbox/projects/mirge3/output_dir/miRge.2021-01-19_15-00-38/miR.Counts.csv", delim = ',')
```

```
## Parsed with column specification:
## cols(
##   miRNA = col_character(),
##   SRR1273998 = col_double(),
##   SRR1273999 = col_double(),
##   SRR1274000 = col_double(),
##   SRR1274001 = col_double()
## )
```

```
seqdata_4 <- read_delim("/Users/eirikhoy/Dropbox/projects/mirge3/output_dir/miRge.2021-01-19_15-33-44/miR.Counts.csv", delim = ',')
```

```
## Parsed with column specification:
## cols(
##   .default = col_double(),
##   miRNA = col_character()
## )
## See spec(...) for full column specifications.
```

```
seqdata_5 <- read_delim("/Users/eirikhoy/Dropbox/projects/mirge3/output_dir/miRge.2021-01-19_17-33-30/miR.Counts.csv", delim = ',')
```

```
## Parsed with column specification:
## cols(
##   .default = col_double(),
##   miRNA = col_character()
## )
## See spec(...) for full column specifications.
```

```
seqdata_6 <- read_delim("/Users/eirikhoy/Dropbox/projects/mirge3/output_dir/miRge.2021-02-04_08-16-37/miR.Counts.csv", delim = ',')
```

```
## Parsed with column specification:
## cols(
##   .default = col_double(),
##   miRNA = col_character()
## )
## See spec(...) for full column specifications.
```

```
seqdata_7 <- read_delim("/Users/eirikhoy/Dropbox/projects/mirge3/output_dir/miRge.2021-02-04_09-09-56/miR.Counts.csv", delim = ',')
```

```
## Parsed with column specification:
## cols(
##   .default = col_double(),
##   miRNA = col_character()
## )
## See spec(...) for full column specifications.
```

```
seqdata_8 <- read_delim("/Users/eirikhoy/Dropbox/projects/mirge3/output_dir/miRge.2021-02-04_12-11-38/miR.Counts.csv", delim = ',')
```

```
## Parsed with column specification:
## cols(
##   .default = col_double(),
##   miRNA = col_character()
## )
## See spec(...) for full column specifications.
```

```
seqdata <- inner_join(inner_join(inner_join(inner_join(inner_join(inner_join(seqdata_1, seqdata_3), seqdata_4), seqdata_5), seqdata_6), seqdata_7), seqdata_8)
```

```
## Joining, by = "miRNA"
```

```
## Joining, by = "miRNA"
## Joining, by = "miRNA"
## Joining, by = "miRNA"
## Joining, by = "miRNA"
## Joining, by = "miRNA"
```

```
colnames(seqdata) <- str_replace_all(colnames(seqdata), pattern='-', replacement = '_')
#colnames(seqdata) <- str_replace_all(colnames(seqdata), pattern='__', replacement = '_')

seqdata <- seqdata[- grep("\\*", seqdata$miRNA), ]
seqdata <- seqdata %>% filter(str_detect(miRNA , "chr", negate = TRUE))
#colnames(seqdata)[2:length(colnames(seqdata))] <- sampleinfo$sample

# Format the data
countdata <- seqdata %>%
  column_to_rownames("miRNA") %>%
  #rename_all(str_remove, ".bam") %>%
  select(sampleinfo$filename) %>%
  as.matrix()

# List consensus samples
consensus <- c("neerincx", "fromm", "schee", "selitsky")

# create the design formula
sampleinfo$tissue.type <- as.factor(paste(sampleinfo$type, sampleinfo$tissue, sep="."))
sampleinfo$type <- as.factor(sampleinfo$type)
design <- as.formula(~ tissue.type)
```

### Differential Expression

```
# Make a named list of signature miRNA
dict_sig_mirna <- c()
res_dict <- list()
```

```
ref <- 'normal.colorect'
dds <- DeseqObject(design, countdata, sampleinfo, "None", "None", ref)
```

```
# #datasets in total
dim(dds[, colData(dds)$type.tissue == 'pCRC'])
```

```
## [1] 389 120
```

```
dim(dds[, colData(dds)$type.tissue == 'mLi'])
```

```
## [1] 389  35
```

```
dim(dds[, colData(dds)$type.tissue == 'mLu'])
```

```
## [1] 389  28
```

```
dim(dds[, colData(dds)$type.tissue == 'nCR'])
```

```
## [1] 389  25
```

```
dim(dds[, colData(dds)$type.tissue == 'nLi'])
```

```
## [1] 389  20
```

```
dim(dds[, colData(dds)$type.tissue == 'nLu'])
```

```
## [1] 389  10
```

```
dim(dds[, colData(dds)$type.tissue == 'PM'])
```

```
## [1] 389  30
```

```
# #datasets for Fromm
dim(dds[, colData(dds)$type.tissue == 'pCRC' & colData(dds)$paper == 'fromm'])
```

```
## [1] 389   3
```

```
dim(dds[, colData(dds)$type.tissue == 'mLi' & colData(dds)$paper == 'fromm'])
```

```
## [1] 389  19
```

```
dim(dds[, colData(dds)$type.tissue == 'mLu' & colData(dds)$paper == 'fromm'])
```

```
## [1] 389  24
```

```
dim(dds[, colData(dds)$type.tissue == 'nCR' & colData(dds)$paper == 'fromm'])
```

```
## [1] 389   3
```

```
dim(dds[, colData(dds)$type.tissue == 'nLi' & colData(dds)$paper == 'fromm'])
```

```
## [1] 389   8
```

```
dim(dds[, colData(dds)$type.tissue == 'nLu' & colData(dds)$paper == 'fromm'])
```

```
## [1] 389   7
```

```
dim(dds[, colData(dds)$type.tissue == 'PM' & colData(dds)$paper == 'fromm'])
```

```
## [1] 389  18
```

```
# #datasets for Schee
dim(dds[, colData(dds)$type.tissue == 'pCRC' & colData(dds)$paper == 'schee'])
```

```
## [1] 389  83
```

```
dim(dds[, colData(dds)$type.tissue == 'mLi' & colData(dds)$paper == 'schee'])
```

```
## [1] 389   0
```

```
dim(dds[, colData(dds)$type.tissue == 'mLu' & colData(dds)$paper == 'schee'])
```

```
## [1] 389   0
```

```
dim(dds[, colData(dds)$type.tissue == 'nCR' & colData(dds)$paper == 'schee'])
```

```
## [1] 389   0
```

```
dim(dds[, colData(dds)$type.tissue == 'nLi' & colData(dds)$paper == 'schee'])
```

```
## [1] 389   0
```

```
dim(dds[, colData(dds)$type.tissue == 'nLu' & colData(dds)$paper == 'schee'])
```

```
## [1] 389   0
```

```
dim(dds[, colData(dds)$type.tissue == 'PM' & colData(dds)$paper == 'schee'])
```

```
## [1] 389   0
```

```
# #datasets for Schee
dim(dds[, colData(dds)$type.tissue == 'pCRC' & colData(dds)$paper == 'neerincx'])
```

```
## [1] 389  34
```

```
dim(dds[, colData(dds)$type.tissue == 'mLi' & colData(dds)$paper == 'neerincx'])
```

```
## [1] 389  16
```

```
dim(dds[, colData(dds)$type.tissue == 'mLu' & colData(dds)$paper == 'neerincx'])
```

```
## [1] 389   4
```

```
dim(dds[, colData(dds)$type.tissue == 'nCR' & colData(dds)$paper == 'neerincx'])
```

```
## [1] 389  22
```

```
dim(dds[, colData(dds)$type.tissue == 'nLi' & colData(dds)$paper == 'neerincx'])
```

```
## [1] 389   9
```

```
dim(dds[, colData(dds)$type.tissue == 'nLu' & colData(dds)$paper == 'neerincx'])
```

```
## [1] 389   3
```

```
dim(dds[, colData(dds)$type.tissue == 'PM' & colData(dds)$paper == 'neerincx'])
```

```
## [1] 389  12
```

```
# #datasets for Schee
dim(dds[, colData(dds)$type.tissue == 'pCRC' & colData(dds)$paper == 'selitsky'])
```

```
## [1] 389   0
```

```
dim(dds[, colData(dds)$type.tissue == 'mLi' & colData(dds)$paper == 'selitsky'])
```

```
## [1] 389   0
```

```
dim(dds[, colData(dds)$type.tissue == 'mLu' & colData(dds)$paper == 'selitsky'])
```

```
## [1] 389   0
```

```
dim(dds[, colData(dds)$type.tissue == 'nCR' & colData(dds)$paper == 'selitsky'])
```

```
## [1] 389   0
```

```
dim(dds[, colData(dds)$type.tissue == 'nLi' & colData(dds)$paper == 'selitsky'])
```

```
## [1] 389   3
```

```
dim(dds[, colData(dds)$type.tissue == 'nLu' & colData(dds)$paper == 'selitsky'])
```

```
## [1] 389   0
```

```
dim(dds[, colData(dds)$type.tissue == 'PM' & colData(dds)$paper == 'selitsky'])
```

```
## [1] 389   0
```

```
## Plot dispersion estimates
plotDispEsts(dds)
```

McCall, Matthew N; Kim, Min-Sik; Adil, Mohammed; Patil, Arun H; Lu, Yin; Mitchell, Christopher J; Leal-Rojas, Pamela; Xu, Jinchong; Kumar, Manoj; Dawson, Valina L; Dawson, Ted M; Baras, Alexander S; Rosenberg, Avi Z; Arking, Dan E; Burns, Kathleen H; Pandey, Akhilesh; Halushka, Marc K Toward the human cellular microRNAome Genome Res. October 2017

```
cell_spec_dict <- list(
  "CD14+ Monocyte"   = c("Hsa-Mir-15-P1a_5p","Hsa-Mir-15-P1b_5p", "Hsa-Mir-17-P1a_5p/P1b_5p"),
  
  "Dendritic Cell"   = c("Hsa-Mir-146-P2_5p", "Hsa-Mir-342_3p", "Hsa-Mir-142_3p",
                         "Hsa-Mir-223_3p"),
  
  "Endothelial Cell" = c("Hsa-Mir-126_5p"),
  
  "Epithelial Cell"  = c("Hsa-Mir-8-P2a_3p", "Hsa-Mir-8-P2b_3p", "Hsa0Mir-205-P1_5p",
                        "Hsa-Mir-192-P1_5p/P2_5p", "Hsa-Mir-375_3p"),
  
  "Islet Cell"       = c("Hsa-Mir-375_3p", "Hsa-Mir-154-P7_5p", "Hsa-Mir-7-P1_5p/P2_5p/P3_5p"),
  
  "Lymphocyte"       = c("Hsa-Mir-146-P2_5p", "Hsa-Mir-342_3p", "Hsa-Mir-150_5p",
                        "Hsa-Mir-155_5p"),
  
  "Macrophage"       = c("Hsa-Mir-342_3p", "Hsa-Mir-142_3p", "Hsa-Mir-223_3p", 
                         "Hsa-Mir-155_5p", "Hsa-Mir-24-P1_3p/P2_3p",
                         "Hsa-Mir-185_5p"),
  
  "Melanocyte"       = c("Hsa-Mir-185_5p", "Hsa-Mir-204-P2_5p"),
  
  "Mesenchymal"      = c("Hsa-Mir-185_5p", "Hsa-Mir-143_3p", "Hsa-Mir-145_5p"),
  
  "Neural"           = c("Hsa-Mir-375_3p", "Hsa-Mir-154-P7_5p", "Hsa-Mir-7-P1_5p/P2_5p/P3_5p",
                         "Hsa-Mir-128-P1_3p/P2_3p", "Hsa-Mir-129-P1_5p/P2_5p",
                         "Hsa-Mir-9-P1_5p/P2_5p/P3_5p","Hsa-Mir-430-P2_3p",
                         "Hsa-Mir-430-P4_3p"),
  
  "Platelet"         = c("Hsa-Mir-126_5p", "Hsa-Mir-486_5p"),
  
  "Red Blood Cell"   = c("Hsa-Mir-486_5p", "Hsa-Mir-451_5p", "Hsa-Mir-144_5p"),
  
  "Retinal Epithelial Cell" = c("Hsa-Mir-204-P1_5p", "Hsa-Mir-204-P2_5p", "Hsa-Mir-335_5p"),
  
  "Skeletal Myocyte" = c("Hsa-Mir-1-P1_3p/P2_3p", "Hsa-Mir-133-P1_3p/P2_3p/P3_3p"),
  
  "Stem Cell"        = c("Hsa-Mir-430-P2_3p", "Hsa-Mir-430-P4_3p", "Hsa-Mir-133-P1_3p/P2_3p/P3_3p"),
  
  "Hepatocyte"        = c("Hsa-Mir-122_5p")
  )

cell_spec_dict_inv <- topGO::inverseList(cell_spec_dict)
```

```
##
```

Pritchard, C C; Kroh, E; Wood, B; Arroyo, J D; Dougherty, K J; Miyaji, M M; Tait, J F; Tewari, M Blood Cell Origin of Circulating MicroRNAs: A Cautionary Note for Cancer Biomarker Studies Cancer Prevention Research 2012

```
blood.cell.mirna <- c("Hsa-Mir-223_3p",
                      "Hsa-Mir-15-P2a_5p",
                      "Hsa-Mir-15-P2b_5p",
                      "Hsa-Mir-126-v1_3p",
                      "Hsa-Mir-142-v1_3p",
                      "Hsa-Mir-21_5p",
                      "Hsa-Mir-24-P1_3p",
                      "Hsa-Mir-24-P2_3p",
                      "Hsa-Mir-19-P2a_3p",
                      "Hsa-Mir-19-P2b_3p",
                      "Hsa-Mir-103-P1_3p",
                      "Hsa-Mir-103-P2_3p",
                      "Hsa-Let-7-P1a_5p",
                      "Hsa-Let-7-P2a1_5p",
                      "Hsa-Let-7-P2a2_5p",
                      "Hsa-Mir-451_5p",
                      "Hsa-Mir-92-P1a_3p",
                      "Hsa-Mir-92-P1b_3p",
                      "Hsa-Mir-17-P1b_5p",
                      "Hsa-Mir-19-P1_3p",
                      "Hsa-Mir-30-P2c_5p",
                      "Hsa-Mir-17-P1a",
                      "Hsa-Mir-15-P1b_5p",
                      "Hsa-Mir-103-P3_3p",
                      "Hsa-Let-7-P2a3_5p",
                      "Hsa-Let-7-P2b1_5p",
                      "Hsa-Mir-221-P1_3p",
                      "Hsa-Mir-221-P2_3p",
                      "Hsa-Mir-17-P1c_5p",
                      "Hsa-Mir-30-P2a_5p",
                      "Hsa-Mir-30-P2b_5p",
                      "Hsa-Mir-28-P2_5p",
                      "Hsa-Mir-30-P1b_5p",
                      "Hsa-Mir-30-P1c_5p",
                      "Hsa-Mir-486_5p",
                      "Hsa-Mir-92-P2c_3p",
                      "Hsa-Mir-181-P1a_5p",
                      "Hsa-Mir-181-P1b_5p",
                      "Hsa-Mir-146-P1_5p",
                      "Hsa-Let-7-P2c1_5p",
                      "Hsa-Mir-197_3p",
                      "Hsa-Mir-17-P3c_5p",
                      "Hsa-Mir-17-P3c_3p",
                      "Hsa-Mir-148-P3_3p",
                      "Hsa-Mir-766_3p",
                      "Hsa-Mir-17-P3b_5p",
                      "Hsa-Mir-328_3p",
                      "Hsa-Mir-574_3p",
                      "Hsa-Mir-155_5p",
                      "Hsa-Mir-425_5p",
                      "Hsa-Mir-148-P1_3p",
                      "Hsa-Mir-29-P1a_3p",
                      "Hsa-Mir-8-P2b_3p",
                      "Hsa-Mir-92-P1c_3p",
                      "Hsa-Mir-192-P2_5p",
                      "Hsa-Mir-362-P2-v1_3p",
                      "Hsa-Mir-362-P5_5p"
                      )
```

#### nCR vs nLi

```
column='tissue.type'
tissue_type_A <- 'normal.liver'
tissue_type_B <- 'normal.colorect'
norm_adj_up   = "None"
norm_adj_down = "None"
pCRC_adj_up   = "None"
pCRC_adj_down = "None"

coef <- paste(column, tissue_type_A, 'vs', tissue_type_B, sep='_')
res <- DeseqResult(dds, column, coef, tissue_type_A, tissue_type_B,
                   lfc.Threshold, rpm.Threshold,
                   norm_adj_up,
                   norm_adj_down)
dict_sig_mirna[paste(coef, "up",   sep='_')] <- list(res$up_mirna)
dict_sig_mirna[paste(coef, "down", sep='_')] <- list(res$down_mirna)
res_res <- res$res
res_dict[coef] <- res_res
plotMA(res$res, alpha=0.05)
```

```
# Plot volcano plot
VolcanoPlot(res$res, coef, res$sig,
            res$up_mirna, res$down_mirna,
            norm_adj_up, norm_adj_down,
            pCRC_adj_up, pCRC_adj_down)
```

```
ExpressionPlot(res$res, res$rpm, coef, res$sig,
               tissue_type_A, tissue_type_B,
               res$up_mirna, res$down_mirna,
               norm_adj_up, norm_adj_down,
               pCRC_adj_up, pCRC_adj_down)
```

```
signature_mirnas <- SigList(res, dds, tissue_type_A, tissue_type_B, coef,
                            norm_adj_up, norm_adj_down, 
                            pCRC_adj_up, pCRC_adj_down)
# Print list upregulated miRNA
signature_mirnas$up_mirna
```

Upregulated in tissue.type\_normal.liver\_vs\_normal.colorect

| miRNA | LFC | lfcSE | FDR | RPM normal.liver | RPM normal.colorect | miRBase\_ID | Family | Seed | Chr | Cell-Type Specific | Norm Background | pCRC Background |
| --- | --- | --- | --- | --- | --- | --- | --- | --- | --- | --- | --- | --- |
| Hsa-Mir-204-P1\_5p | 1.56 | 0.51 | 4.46e-03 | 190 | 43 | hsa-mir-204 | MIR-204 | UCCCUUU | chr9 | Retinal Epithelial Cell | na | na |
| Hsa-Mir-335\_5p | 1.18 | 0.17 | 2.56e-11 | 205 | 79 | hsa-mir-335 | MIR-335 | CAAGAGC | chr7 | Retinal Epithelial Cell | na | na |
| Hsa-Mir-144\_5p | 1.53 | 0.33 | 5.21e-05 | 206 | 65 | hsa-mir-144 | MIR-144 | GAUAUCA | chr17 | Red Blood Cell | na | na |
| Hsa-Mir-128-P1\_3p/P2\_3p | 0.65 | 0.16 | 2.62e-04 | 194 | 110 | hsa-mir-128-1 | MIR-128 | CACAGUG | chr2 | Neural | na | na |
| Hsa-Mir-122\_5p | 10.58 | 0.78 | 7.97e-28 | 148842 | 26 | hsa-mir-122 | MIR-122 | GGAGUGU | chr18 | Hepatocyte | na | na |
| Hsa-Mir-486\_5p | 1.34 | 0.36 | 2.99e-03 | 10617 | 3961 | hsa-mir-486-1 | MIR-486 | CCUGUAC | chr8 | c(“Platelet”, “Red Blood Cell”) | na | na |
| Hsa-Mir-126\_5p | 1.25 | 0.21 | 1.87e-08 | 11148 | 4230 | hsa-mir-126 | MIR-126 | AUUAUUA | chr9 | c(“Endothelial Cell”, “Platelet”) | na | na |
| Hsa-Mir-885\_5p | 8.77 | 0.70 | 8.60e-24 | 510 | 0 | hsa-mir-885 | MIR-885 | CCAUUAC | chr3 |  | na | na |
| Hsa-Mir-483\_5p | 4.03 | 0.71 | 1.04e-10 | 115 | 1 | hsa-mir-483 | MIR-483 | AGACGGG | chr11 |  | na | na |
| Hsa-Let-7-P1c\_5p | 3.18 | 0.28 | 1.60e-27 | 4945 | 466 | hsa-let-7c | LET-7 | GAGGUAG | chr21 |  | na | na |
| Hsa-Mir-10-P2c\_5p | 3.08 | 0.33 | 6.34e-19 | 1202 | 114 | hsa-mir-99a | MIR-10 | ACCCGUA | chr21 |  | na | na |
| Hsa-Mir-455\_5p | 2.48 | 0.18 | 9.25e-41 | 279 | 44 | hsa-mir-455 | MIR-455 | AUGUGCC | chr9 |  | na | na |
| Hsa-Mir-193-P2a\_3p/P2b\_3p | 2.37 | 0.23 | 5.29e-22 | 219 | 37 | hsa-mir-365b | MIR-193 | AAUGCCC | chr17 |  | na | na |
| Hsa-Mir-193-P1b\_3p | 2.34 | 0.23 | 2.49e-22 | 514 | 90 | hsa-mir-193b | MIR-193 | ACUGGCC | chr16 |  | na | na |
| Hsa-Mir-15-P1d\_5p | 2.26 | 0.30 | 5.34e-14 | 247 | 40 | hsa-mir-424 | MIR-15 | AGCAGCA | chrX |  | na | na |
| Hsa-Mir-193-P1a\_5p | 2.14 | 0.28 | 3.66e-12 | 298 | 57 | hsa-mir-193a | MIR-193 | GGGUCUU | chr17 |  | na | na |
| Hsa-Mir-10-P3c\_5p | 2.12 | 0.30 | 6.48e-11 | 1620 | 326 | hsa-mir-125b-2 | MIR-10 | CCCUGAG | chr21 |  | na | na |
| Hsa-Mir-139\_5p | 2.00 | 0.26 | 1.06e-11 | 174 | 41 | hsa-mir-139 | MIR-139 | CUACAGU | chr11 |  | na | na |
| Hsa-Mir-148-P1\_3p | 2.00 | 0.24 | 8.34e-15 | 105086 | 21730 | hsa-mir-148a | MIR-148 | CAGUGCA | chr7 |  | na | na |
| Hsa-Mir-10-P3a\_5p | 1.93 | 0.31 | 1.24e-08 | 611 | 137 | hsa-mir-125b-1 | MIR-10 | CCCUGAG | chr11 |  | na | na |
| Hsa-Mir-423\_5p | 1.62 | 0.24 | 1.23e-09 | 1198 | 345 | hsa-mir-423 | MIR-423 | GAGGGGC | chr17 |  | na | na |
| Hsa-Mir-92-P1a\_3p/P1b\_3p | 1.60 | 0.22 | 1.98e-12 | 51173 | 13849 | hsa-mir-92a-1 | MIR-92 | AUUGCAC | chr13 |  | na | na |
| Hsa-Mir-101-P1\_3p/P2\_3p | 1.58 | 0.18 | 3.44e-17 | 14649 | 4197 | hsa-mir-101-1 | MIR-101 | UACAGUA | chr1 |  | na | na |
| Hsa-Mir-22-P1a\_3p | 1.58 | 0.17 | 1.76e-18 | 78932 | 23453 | hsa-mir-22 | MIR-22 | AGCUGCC | chr17 |  | na | na |
| Hsa-Mir-340\_5p | 1.42 | 0.16 | 3.45e-18 | 1030 | 337 | hsa-mir-340 | MIR-340 | UAUAAAG | chr5 |  | na | na |
| Hsa-Mir-30-P1a\_5p | 1.32 | 0.23 | 1.07e-07 | 15175 | 5150 | hsa-mir-30a | MIR-30 | GUAAACA | chr6 |  | na | na |
| Hsa-Mir-744\_5p | 1.31 | 0.21 | 1.28e-08 | 165 | 59 | hsa-mir-744 | MIR-744 | GCGGGGC | chr17 |  | na | na |
| Hsa-Mir-574\_3p | 1.21 | 0.19 | 1.27e-08 | 745 | 305 | hsa-mir-574 | MIR-574 | ACGCUCA | chr4 |  | na | na |
| Hsa-Mir-130-P1a\_3p | 1.09 | 0.20 | 4.15e-07 | 907 | 379 | hsa-mir-130a | MIR-130 | AGUGCAA | chr11 |  | na | na |
| Hsa-Mir-10-P2a\_5p | 1.07 | 0.32 | 3.39e-03 | 2634 | 986 | hsa-mir-100 | MIR-10 | ACCCGUA | chr11 |  | na | na |
| Hsa-Mir-30-P2a\_5p/P2b\_5p/P2c\_5p | 0.99 | 0.15 | 4.69e-10 | 5891 | 2677 | hsa-mir-30c-2 | MIR-30 | GUAAACA | chr6 |  | na | na |
| Hsa-Mir-154-P23\_3p | 0.89 | 0.20 | 5.21e-05 | 222 | 108 | hsa-mir-654 | MIR-154 | AUGUCUG | chr14 |  | na | na |
| Hsa-Mir-197\_3p | 0.79 | 0.19 | 2.06e-04 | 344 | 180 | hsa-mir-197 | MIR-197 | UCACCAC | chr1 |  | na | na |
| Hsa-Mir-214\_3p | 0.73 | 0.23 | 5.91e-03 | 137 | 74 | hsa-mir-214 | MIR-214 | CAGCAGG | chr1 |  | na | na |
| Hsa-Mir-30-P1b\_5p | 0.71 | 0.15 | 7.77e-06 | 11240 | 6046 | hsa-mir-30e | MIR-30 | GUAAACA | chr1 |  | na | na |
| Hsa-Mir-30-P1c\_5p | 0.70 | 0.16 | 7.79e-05 | 13680 | 7243 | hsa-mir-30d | MIR-30 | GUAAACA | chr8 |  | na | na |

```
# Number of upregulated miRNA
signature_mirnas$number_upregulated
```

```
## [1] 36
```

```
# Print list downregulated miRNA
signature_mirnas$down_mirna
```

Downregulated in tissue.type\_normal.liver\_vs\_normal.colorect

| miRNA | LFC | lfcSE | FDR | RPM normal.liver | RPM normal.colorect | miRBase\_ID | Family | Seed | Chr | Cell-Type Specific | Norm Background | pCRC Background |
| --- | --- | --- | --- | --- | --- | --- | --- | --- | --- | --- | --- | --- |
| Hsa-Mir-145\_5p | -2.51 | 0.32 | 5.55e-15 | 484 | 2832 | hsa-mir-145 | MIR-145 | UCCAGUU | chr5 | Mesenchymal | na | na |
| Hsa-Mir-143\_3p | -2.35 | 0.27 | 1.80e-17 | 37572 | 164812 | hsa-mir-143 | MIR-143 | GAGAUGA | chr5 | Mesenchymal | na | na |
| Hsa-Mir-24-P1\_3p/P2\_3p | -0.63 | 0.20 | 5.98e-03 | 319 | 413 | hsa-mir-24-2 | MIR-24 | GGCUCAG | chr19 | Macrophage | na | na |
| Hsa-Mir-8-P2b\_3p | -6.38 | 0.23 | 7.72e-155 | 31 | 2444 | hsa-mir-200c | MIR-8 | AAUACUG | chr12 | Epithelial Cell | na | na |
| Hsa-Mir-8-P2a\_3p | -4.47 | 0.24 | 4.08e-76 | 369 | 7741 | hsa-mir-200b | MIR-8 | AAUACUG | chr1 | Epithelial Cell | na | na |
| Hsa-Mir-192-P1\_5p/P2\_5p | -0.72 | 0.32 | 1.03e-02 | 29546 | 46010 | hsa-mir-192 | MIR-192 | UGACCUA | chr11 | Epithelial Cell | na | na |
| Hsa-Mir-15-P1b\_5p | -0.68 | 0.16 | 1.18e-04 | 114 | 168 | hsa-mir-15b | MIR-15 | AGCAGCA | chr3 | CD14+ Monocyte | na | na |
| Hsa-Mir-133-P1\_3p/P2\_3p/P3\_3p | -4.10 | 0.38 | 9.37e-29 | 22 | 460 | hsa-mir-133a-2 | MIR-133 | UUGGUCC | chr20 | c(“Skeletal Myocyte”, “Stem Cell”) | na | na |
| Hsa-Mir-155\_5p | -1.51 | 0.26 | 8.41e-08 | 113 | 295 | hsa-mir-155 | MIR-155 | UAAUGCU | chr21 | c(“Lymphocyte”, “Macrophage”) | na | na |
| Hsa-Mir-375\_3p | -2.65 | 0.37 | 1.69e-12 | 2553 | 17215 | hsa-mir-375 | MIR-375 | UUGUUCG | chr2 | c(“Epithelial Cell”, “Islet Cell”, “Neural”) | na | na |
| Hsa-Mir-196-P1\_5p/P2\_5p | -6.76 | 0.34 | 3.97e-79 | 2 | 250 | hsa-mir-196a-1 | MIR-196 | AGGUAGU | chr17 |  | na | na |
| Hsa-Mir-147\_3p | -6.57 | 0.38 | 2.57e-64 | 1 | 110 | hsa-mir-147b | MIR-147 | UGUGCGG | chr15 |  | na | na |
| Hsa-Mir-577\_5p | -6.02 | 0.34 | 1.65e-65 | 2 | 138 | hsa-mir-577 | MIR-577 | UAGAUAA | chr4 |  | na | na |
| Hsa-Mir-8-P1b\_3p | -6.00 | 0.30 | 3.97e-79 | 154 | 9180 | hsa-mir-141 | MIR-8 | AACACUG | chr12 |  | na | na |
| Hsa-Mir-196-P3\_5p | -5.53 | 0.37 | 2.24e-42 | 9 | 400 | hsa-mir-196b | MIR-196 | AGGUAGU | chr7 |  | na | na |
| Hsa-Mir-10-P1b\_5p | -4.80 | 0.36 | 6.31e-38 | 2704 | 75067 | hsa-mir-10b | MIR-10 | ACCCUGU | chr2 |  | na | na |
| Hsa-Mir-8-P3a\_3p | -4.32 | 0.24 | 2.00e-68 | 68 | 1282 | hsa-mir-429 | MIR-8 | AAUACUG | chr1 |  | na | na |
| Hsa-Mir-8-P1a\_3p | -4.10 | 0.24 | 4.38e-64 | 108 | 1754 | hsa-mir-200a | MIR-8 | AACACUG | chr1 |  | na | na |
| Hsa-Mir-190-P1\_5p | -3.71 | 0.26 | 4.48e-45 | 23 | 294 | hsa-mir-190a | MIR-190 | GAUAUGU | chr15 |  | na | na |
| Hsa-Mir-96-P3\_5p | -3.57 | 0.32 | 1.72e-22 | 27 | 224 | hsa-mir-183 | MIR-96 | AUGGCAC | chr7 |  | na | na |
| Hsa-Mir-203\_3p | -2.68 | 0.28 | 6.56e-18 | 171 | 922 | hsa-mir-203a | MIR-203 | UGAAAUG | chr14 |  | na | na |
| Hsa-Mir-96-P2\_5p | -2.65 | 0.24 | 2.99e-23 | 370 | 1867 | hsa-mir-182 | MIR-96 | UUGGCAA | chr7 |  | na | na |
| Hsa-Mir-221-P1\_3p | -2.30 | 0.19 | 5.81e-32 | 217 | 964 | hsa-mir-221 | MIR-221 | GCUACAU | chrX |  | na | na |
| Hsa-Mir-338-P1\_3p | -1.89 | 0.31 | 1.21e-08 | 48 | 166 | hsa-mir-338 | MIR-338 | CCAGCAU | chr17 |  | na | na |
| Hsa-Mir-221-P2\_3p | -1.87 | 0.20 | 9.25e-18 | 173 | 574 | hsa-mir-222 | MIR-221 | GCUACAU | chrX |  | na | na |
| Hsa-Mir-10-P1a\_5p | -1.77 | 0.30 | 1.66e-07 | 25828 | 70250 | hsa-mir-10a | MIR-10 | ACCCUGU | chr17 |  | na | na |
| Hsa-Mir-146-P1\_5p | -1.69 | 0.34 | 9.05e-06 | 351 | 891 | hsa-mir-146a | MIR-146 | GAGAACU | chr5 |  | na | na |
| Hsa-Mir-92-P1c\_3p | -1.32 | 0.26 | 4.00e-06 | 303 | 634 | hsa-mir-92b | MIR-92 | AUUGCAC | chr1 |  | na | na |
| Hsa-Mir-181-P1c\_5p | -1.25 | 0.21 | 6.34e-08 | 261 | 527 | hsa-mir-181c | MIR-181 | ACAUUCA | chr19 |  | na | na |
| Hsa-Mir-425\_5p | -1.16 | 0.19 | 6.36e-09 | 247 | 499 | hsa-mir-425 | MIR-425 | AUGACAC | chr3 |  | na | na |
| Hsa-Mir-362-P3\_3p | -1.15 | 0.36 | 2.88e-03 | 60 | 103 | hsa-mir-501 | MIR-362 | AUGCACC | chrX |  | na | na |
| Hsa-Mir-192-P1\_5p | -1.10 | 0.27 | 7.22e-05 | 109276 | 212293 | hsa-mir-192 | MIR-192 | UGACCUA | chr11 |  | na | na |
| Hsa-Mir-10-P2b\_5p | -1.08 | 0.36 | 6.81e-03 | 1478 | 2499 | hsa-mir-99b | MIR-10 | ACCCGUA | chr19 |  | na | na |
| Hsa-Mir-210\_3p | -1.07 | 0.28 | 1.53e-03 | 106 | 185 | hsa-mir-210 | MIR-210 | UGUGCGU | chr11 |  | na | na |
| Hsa-Mir-132-P1\_3p | -1.03 | 0.20 | 2.09e-06 | 61 | 117 | hsa-mir-132 | MIR-132 | AACAGUC | chr17 |  | na | na |
| Hsa-Mir-194-P1\_5p/P2\_5p | -0.94 | 0.27 | 5.19e-04 | 6323 | 11619 | hsa-mir-194-2 | MIR-194 | GUAACAG | chr11 |  | na | na |
| Hsa-Mir-130-P4a\_3p | -0.84 | 0.13 | 2.77e-10 | 70 | 111 | hsa-mir-454 | MIR-130 | AGUGCAA | chr17 |  | na | na |
| Hsa-Mir-188-P2\_5p | -0.82 | 0.19 | 9.56e-05 | 279 | 427 | hsa-mir-532 | MIR-188 | AUGCCUU | chrX |  | na | na |
| Hsa-Mir-191\_5p | -0.78 | 0.23 | 1.91e-03 | 10721 | 14608 | hsa-mir-191 | MIR-191 | AACGGAA | chr3 |  | na | na |
| Hsa-Mir-21\_5p | -0.76 | 0.17 | 1.54e-04 | 19240 | 28173 | hsa-mir-21 | MIR-21 | AGCUUAU | chr17 |  | na | na |
| Hsa-Mir-130-P2a\_3p | -0.71 | 0.20 | 3.57e-03 | 92 | 131 | hsa-mir-301a | MIR-130 | AGUGCAA | chr17 |  | na | na |
| Hsa-Mir-1307\_3p | -0.66 | 0.19 | 1.89e-03 | 89 | 127 | hsa-mir-1307 | MIR-1307 | CGACCGG | chr10 |  | na | na |
| Hsa-Mir-362-P2\_3p/P4\_3p | -0.61 | 0.20 | 7.29e-03 | 211 | 268 | hsa-mir-500a | MIR-362 | UGCACCU | chrX |  | na | na |
| Hsa-Mir-1307\_5p | -0.61 | 0.26 | 3.36e-02 | 502 | 713 | hsa-mir-1307 | MIR-1307 | CGACCGG | chr10 |  | na | na |

```
# Number of downregulated miRNA
signature_mirnas$number_downregulated
```

```
## [1] 44
```

#### nCR vs nLu

```
column='tissue.type'
tissue_type_A <- 'normal.lung'
tissue_type_B <- 'normal.colorect'
norm_adj_up       = "None"
norm_adj_down     = "None"
pCRC_adj_up   = "None"
pCRC_adj_down = "None"

coef <- paste(column, tissue_type_A, 'vs', tissue_type_B, sep='_')
res <- DeseqResult(dds, column, coef, tissue_type_A, tissue_type_B,
                   lfc.Threshold, rpm.Threshold,
                   norm_adj_up,
                   norm_adj_down)
dict_sig_mirna[paste(coef, "up",   sep='_')] <- list(res$up_mirna)
dict_sig_mirna[paste(coef, "down", sep='_')] <- list(res$down_mirna)
res_res <- res$res
res_dict[coef] <- res_res
plotMA(res$res, alpha=0.05)
```

```
# Plot volcano plot
VolcanoPlot(res$res, coef, res$sig,
            res$up_mirna, res$down_mirna,
            norm_adj_up, norm_adj_down,
            pCRC_adj_up, pCRC_adj_down)
```

```
ExpressionPlot(res$res, res$rpm, coef, res$sig,
               tissue_type_A, tissue_type_B,
               res$up_mirna, res$down_mirna,
               norm_adj_up, norm_adj_down,
               pCRC_adj_up, pCRC_adj_down)
```

```
signature_mirnas <- SigList(res, dds, tissue_type_A, tissue_type_B, coef,
                            norm_adj_up, norm_adj_down, 
                            pCRC_adj_up, pCRC_adj_down)
# Print list upregulated miRNA
signature_mirnas$up_mirna
```

Upregulated in tissue.type\_normal.lung\_vs\_normal.colorect

| miRNA | LFC | lfcSE | FDR | RPM normal.lung | RPM normal.colorect | miRBase\_ID | Family | Seed | Chr | Cell-Type Specific | Norm Background | pCRC Background |
| --- | --- | --- | --- | --- | --- | --- | --- | --- | --- | --- | --- | --- |
| Hsa-Mir-335\_5p | 1.48 | 0.21 | 3.73e-11 | 300 | 79 | hsa-mir-335 | MIR-335 | CAAGAGC | chr7 | Retinal Epithelial Cell | na | na |
| Hsa-Mir-144\_5p | 2.32 | 0.41 | 5.46e-07 | 451 | 65 | hsa-mir-144 | MIR-144 | GAUAUCA | chr17 | Red Blood Cell | na | na |
| Hsa-Mir-451\_5p | 1.90 | 0.47 | 1.06e-03 | 12496 | 2815 | hsa-mir-451a | MIR-451 | AACCGUU | chr17 | Red Blood Cell | na | na |
| Hsa-Mir-24-P1\_3p/P2\_3p | 0.86 | 0.25 | 1.88e-03 | 1084 | 413 | hsa-mir-24-2 | MIR-24 | GGCUCAG | chr19 | Macrophage | na | na |
| Hsa-Mir-486\_5p | 2.04 | 0.45 | 1.33e-04 | 22081 | 3961 | hsa-mir-486-1 | MIR-486 | CCUGUAC | chr8 | c(“Platelet”, “Red Blood Cell”) | na | na |
| Hsa-Mir-126\_5p | 2.57 | 0.26 | 4.06e-21 | 33620 | 4230 | hsa-mir-126 | MIR-126 | AUUAUUA | chr9 | c(“Endothelial Cell”, “Platelet”) | na | na |
| Hsa-Mir-146-P2\_5p | 1.65 | 0.41 | 1.94e-04 | 17435 | 3259 | hsa-mir-146b | MIR-146 | GAGAACU | chr10 | c(“Dendritic Cell”, “Lymphocyte”) | na | na |
| Hsa-Mir-342\_3p | 1.00 | 0.31 | 6.94e-03 | 681 | 260 | hsa-mir-342 | MIR-342 | CUCACAC | chr14 | c(“Dendritic Cell”, “Lymphocyte”, “Macrophage”) | na | na |
| Hsa-Mir-34-P2b\_5p | 5.29 | 0.43 | 1.18e-33 | 748 | 11 | hsa-mir-34c | MIR-34 | GGCAGUG | chr11 |  | na | na |
| Hsa-Mir-34-P2a\_5p | 4.98 | 0.49 | 3.17e-22 | 116 | 2 | hsa-mir-34b | MIR-34 | GGCAGUG | chr11 |  | na | na |
| Hsa-Mir-184\_3p | 4.75 | 0.92 | 5.98e-06 | 111 | 1 | hsa-mir-184 | MIR-184 | GGACGGA | chr15 |  | na | na |
| Hsa-Mir-30-P1a\_5p | 3.04 | 0.28 | 6.17e-24 | 60860 | 5150 | hsa-mir-30a | MIR-30 | GUAAACA | chr6 |  | na | na |
| Hsa-Mir-218-P1\_5p/P2\_5p | 2.39 | 0.33 | 7.14e-11 | 374 | 52 | hsa-mir-218-1 | MIR-218 | UGUGCUU | chr4 |  | na | na |
| Hsa-Mir-10-P2c\_5p | 2.23 | 0.41 | 4.40e-07 | 804 | 114 | hsa-mir-99a | MIR-10 | ACCCGUA | chr21 |  | na | na |
| Hsa-Mir-181-P1a\_5p/P1b\_5p | 2.19 | 0.24 | 1.16e-18 | 80063 | 12321 | hsa-mir-181a-1 | MIR-181 | ACAUUCA | chr1 |  | na | na |
| Hsa-Mir-181-P2a\_5p/P2b\_5p | 2.06 | 0.23 | 2.25e-18 | 3799 | 650 | hsa-mir-181b-1 | MIR-181 | ACAUUCA | chr1 |  | na | na |
| Hsa-Let-7-P1c\_5p | 1.97 | 0.35 | 1.57e-07 | 2606 | 466 | hsa-let-7c | LET-7 | GAGGUAG | chr21 |  | na | na |
| Hsa-Mir-130-P1a\_3p | 1.92 | 0.25 | 5.89e-13 | 1941 | 379 | hsa-mir-130a | MIR-130 | AGUGCAA | chr11 |  | na | na |
| Hsa-Mir-30-P1c\_5p | 1.74 | 0.20 | 4.37e-16 | 34088 | 7243 | hsa-mir-30d | MIR-30 | GUAAACA | chr8 |  | na | na |
| Hsa-Mir-101-P1\_3p/P2\_3p | 1.45 | 0.22 | 1.42e-09 | 16084 | 4197 | hsa-mir-101-1 | MIR-101 | UACAGUA | chr1 |  | na | na |
| Hsa-Mir-338-P1\_3p | 1.45 | 0.39 | 2.15e-03 | 605 | 166 | hsa-mir-338 | MIR-338 | CCAGCAU | chr17 |  | na | na |
| Hsa-Mir-181-P2c\_5p | 1.42 | 0.28 | 1.14e-06 | 200 | 51 | hsa-mir-181d | MIR-181 | ACAUUCA | chr19 |  | na | na |
| Hsa-Mir-10-P2a\_5p | 1.40 | 0.40 | 2.08e-03 | 4056 | 986 | hsa-mir-100 | MIR-10 | ACCCGUA | chr11 |  | na | na |
| Hsa-Mir-10-P3a\_5p | 1.37 | 0.38 | 1.96e-03 | 507 | 137 | hsa-mir-125b-1 | MIR-10 | CCCUGAG | chr11 |  | na | na |
| Hsa-Mir-10-P3b\_5p | 1.34 | 0.34 | 7.55e-04 | 8561 | 2392 | hsa-mir-125a | MIR-10 | CCCUGAG | chr19 |  | na | na |
| Hsa-Mir-181-P1c\_5p | 1.25 | 0.26 | 5.14e-06 | 1790 | 527 | hsa-mir-181c | MIR-181 | ACAUUCA | chr19 |  | na | na |
| Hsa-Mir-140\_3p | 1.25 | 0.19 | 1.88e-09 | 2195 | 686 | hsa-mir-140 | MIR-140 | CCACAGG | chr16 |  | na | na |
| Hsa-Let-7-P2b1\_5p | 1.10 | 0.15 | 1.06e-11 | 6253 | 2110 | hsa-let-7f-1 | LET-7 | GAGGUAG | chr9 |  | na | na |
| Hsa-Mir-30-P2a\_5p/P2b\_5p/P2c\_5p | 1.08 | 0.19 | 9.04e-08 | 7603 | 2677 | hsa-mir-30c-2 | MIR-30 | GUAAACA | chr6 |  | na | na |
| Hsa-Mir-221-P1\_3p | 0.95 | 0.23 | 1.40e-04 | 2502 | 964 | hsa-mir-221 | MIR-221 | GCUACAU | chrX |  | na | na |
| Hsa-Mir-92-P1c\_3p | 0.90 | 0.32 | 1.56e-02 | 1737 | 634 | hsa-mir-92b | MIR-92 | AUUGCAC | chr1 |  | na | na |
| Hsa-Mir-15-P2c\_5p | 0.86 | 0.30 | 2.04e-02 | 1552 | 672 | hsa-mir-195 | MIR-15 | AGCAGCA | chr17 |  | na | na |
| Hsa-Mir-374-P2\_5p | 0.80 | 0.25 | 3.59e-03 | 148 | 65 | hsa-mir-374b | MIR-374 | UAUAAUA | chrX |  | na | na |
| Hsa-Mir-221-P2\_3p | 0.77 | 0.25 | 5.86e-03 | 1346 | 574 | hsa-mir-222 | MIR-221 | GCUACAU | chrX |  | na | na |
| Hsa-Let-7-P2c2\_5p | 0.76 | 0.21 | 1.16e-03 | 5088 | 2218 | hsa-let-7i | LET-7 | GAGGUAG | chr12 |  | na | na |
| Hsa-Mir-130-P2a\_3p | 0.75 | 0.25 | 6.48e-03 | 305 | 131 | hsa-mir-301a | MIR-130 | AGUGCAA | chr17 |  | na | na |
| Hsa-Mir-652\_3p | 0.71 | 0.20 | 1.38e-03 | 187 | 86 | hsa-mir-652 | MIR-652 | AUGGCGC | chrX |  | na | na |
| Hsa-Mir-26-P1\_5p/P2\_5p | 0.68 | 0.17 | 3.54e-04 | 123947 | 56268 | hsa-mir-26b | MIR-26 | UCAAGUA | chr2 |  | na | na |
| Hsa-Mir-28-P2\_3p | 0.63 | 0.21 | 9.02e-03 | 6007 | 2638 | hsa-mir-151a | MIR-28 | CGAGGAG | chr8 |  | na | na |

```
# Number of upregulated miRNA
signature_mirnas$number_upregulated
```

```
## [1] 39
```

```
# Print list downregulated miRNA
signature_mirnas$down_mirna
```

Downregulated in tissue.type\_normal.lung\_vs\_normal.colorect

| miRNA | LFC | lfcSE | FDR | RPM normal.lung | RPM normal.colorect | miRBase\_ID | Family | Seed | Chr | Cell-Type Specific | Norm Background | pCRC Background |
| --- | --- | --- | --- | --- | --- | --- | --- | --- | --- | --- | --- | --- |
| Hsa-Mir-143\_3p | -0.81 | 0.34 | 2.00e-02 | 133600 | 164812 | hsa-mir-143 | MIR-143 | GAGAUGA | chr5 | Mesenchymal | na | na |
| Hsa-Mir-192-P1\_5p/P2\_5p | -8.51 | 0.40 | 3.64e-97 | 142 | 46010 | hsa-mir-192 | MIR-192 | UGACCUA | chr11 | Epithelial Cell | na | na |
| Hsa-Mir-8-P2a\_3p | -4.00 | 0.29 | 6.74e-40 | 618 | 7741 | hsa-mir-200b | MIR-8 | AAUACUG | chr1 | Epithelial Cell | na | na |
| Hsa-Mir-8-P2b\_3p | -2.06 | 0.29 | 1.04e-11 | 766 | 2444 | hsa-mir-200c | MIR-8 | AAUACUG | chr12 | Epithelial Cell | na | na |
| Hsa-Mir-17-P1a\_5p/P1b\_5p | -0.77 | 0.24 | 9.07e-03 | 184 | 225 | hsa-mir-17 | MIR-17 | AAAGUGC | chr13 | CD14+ Monocyte | na | na |
| Hsa-Mir-133-P1\_3p/P2\_3p/P3\_3p | -2.33 | 0.47 | 9.21e-08 | 92 | 460 | hsa-mir-133a-2 | MIR-133 | UUGGUCC | chr20 | c(“Skeletal Myocyte”, “Stem Cell”) | na | na |
| Hsa-Mir-375\_3p | -3.01 | 0.47 | 2.15e-10 | 2346 | 17215 | hsa-mir-375 | MIR-375 | UUGUUCG | chr2 | c(“Epithelial Cell”, “Islet Cell”, “Neural”) | na | na |
| Hsa-Mir-194-P1\_5p/P2\_5p | -7.89 | 0.34 | 2.93e-114 | 58 | 11619 | hsa-mir-194-2 | MIR-194 | GUAACAG | chr11 |  | na | na |
| Hsa-Mir-192-P1\_5p | -7.28 | 0.34 | 3.81e-95 | 1684 | 212293 | hsa-mir-192 | MIR-192 | UGACCUA | chr11 |  | na | na |
| Hsa-Mir-196-P1\_5p/P2\_5p | -6.68 | 0.41 | 1.42e-54 | 3 | 250 | hsa-mir-196a-1 | MIR-196 | AGGUAGU | chr17 |  | na | na |
| Hsa-Mir-196-P3\_5p | -5.95 | 0.47 | 9.59e-33 | 7 | 400 | hsa-mir-196b | MIR-196 | AGGUAGU | chr7 |  | na | na |
| Hsa-Mir-577\_5p | -5.95 | 0.41 | 4.39e-45 | 3 | 138 | hsa-mir-577 | MIR-577 | UAGAUAA | chr4 |  | na | na |
| Hsa-Mir-147\_3p | -5.82 | 0.43 | 5.54e-40 | 2 | 110 | hsa-mir-147b | MIR-147 | UGUGCGG | chr15 |  | na | na |
| Hsa-Mir-190-P1\_5p | -4.03 | 0.32 | 1.35e-34 | 22 | 294 | hsa-mir-190a | MIR-190 | GAUAUGU | chr15 |  | na | na |
| Hsa-Mir-378\_3p | -3.17 | 0.24 | 2.10e-38 | 2189 | 14834 | hsa-mir-378a | MIR-378 | CUGGACU | chr5 |  | na | na |
| Hsa-Mir-127\_3p | -2.96 | 0.40 | 4.55e-12 | 327 | 1967 | hsa-mir-127 | MIR-127 | CGGAUCC | chr14 |  | na | na |
| Hsa-Mir-10-P1b\_5p | -2.67 | 0.45 | 5.72e-09 | 14125 | 75067 | hsa-mir-10b | MIR-10 | ACCCUGU | chr2 |  | na | na |
| Hsa-Mir-8-P1a\_3p | -2.66 | 0.29 | 3.55e-18 | 353 | 1754 | hsa-mir-200a | MIR-8 | AACACUG | chr1 |  | na | na |
| Hsa-Mir-8-P3a\_3p | -2.65 | 0.30 | 1.86e-17 | 262 | 1282 | hsa-mir-429 | MIR-8 | AAUACUG | chr1 |  | na | na |
| Hsa-Mir-154-P23\_3p | -1.79 | 0.25 | 3.73e-11 | 41 | 108 | hsa-mir-654 | MIR-154 | AUGUCUG | chr14 |  | na | na |
| Hsa-Mir-425\_5p | -1.66 | 0.23 | 3.12e-11 | 213 | 499 | hsa-mir-425 | MIR-425 | AUGACAC | chr3 |  | na | na |
| Hsa-Mir-154-P9\_3p | -1.37 | 0.26 | 6.98e-07 | 135 | 265 | hsa-mir-381 | MIR-154 | AUACAAG | chr14 |  | na | na |
| Hsa-Mir-154-P13\_5p | -1.35 | 0.27 | 3.42e-06 | 161 | 311 | hsa-mir-411 | MIR-154 | AGUAGAC | chr14 |  | na | na |
| Hsa-Mir-28-P1\_3p | -1.24 | 0.23 | 3.62e-07 | 2385 | 3956 | hsa-mir-28 | MIR-28 | ACUAGAU | chr3 |  | na | na |
| Hsa-Mir-17-P3a\_5p | -1.23 | 0.29 | 2.53e-04 | 194 | 322 | hsa-mir-20a | MIR-17 | AAAGUGC | chr13 |  | na | na |
| Hsa-Mir-1307\_3p | -1.09 | 0.24 | 2.47e-05 | 80 | 127 | hsa-mir-1307 | MIR-1307 | CGACCGG | chr10 |  | na | na |
| Hsa-Mir-8-P1b\_3p | -1.06 | 0.38 | 1.07e-02 | 6145 | 9180 | hsa-mir-141 | MIR-8 | AACACUG | chr12 |  | na | na |
| Hsa-Mir-1307\_5p | -1.03 | 0.32 | 4.10e-03 | 444 | 713 | hsa-mir-1307 | MIR-1307 | CGACCGG | chr10 |  | na | na |
| Hsa-Mir-362-P3\_3p | -1.00 | 0.45 | 4.09e-02 | 82 | 103 | hsa-mir-501 | MIR-362 | AUGCACC | chrX |  | na | na |
| Hsa-Mir-210\_3p | -0.93 | 0.35 | 3.36e-02 | 138 | 185 | hsa-mir-210 | MIR-210 | UGUGCGU | chr11 |  | na | na |
| Hsa-Mir-136\_3p | -0.89 | 0.27 | 2.95e-03 | 88 | 127 | hsa-mir-136 | MIR-136 | AUCAUCG | chr14 |  | na | na |
| Hsa-Mir-21\_5p | -0.86 | 0.22 | 6.21e-04 | 21349 | 28173 | hsa-mir-21 | MIR-21 | AGCUUAU | chr17 |  | na | na |
| Hsa-Mir-30-P1b\_5p | -0.86 | 0.18 | 8.68e-06 | 4513 | 6046 | hsa-mir-30e | MIR-30 | GUAAACA | chr1 |  | na | na |
| Hsa-Mir-191\_5p | -0.82 | 0.28 | 1.02e-02 | 12597 | 14608 | hsa-mir-191 | MIR-191 | AACGGAA | chr3 |  | na | na |
| Hsa-Mir-148-P1\_3p | -0.72 | 0.30 | 4.78e-02 | 18874 | 21730 | hsa-mir-148a | MIR-148 | CAGUGCA | chr7 |  | na | na |

```
# Number of downregulated miRNA
signature_mirnas$number_downregulated
```

```
## [1] 35
```

#### nCR vs pCRC

```
column='tissue.type'
tissue_type_A <- 'tumor.colorect'
tissue_type_B <- 'normal.colorect'
norm_adj_up       = "None"
norm_adj_down     = "None"
pCRC_adj_up   = "None"
pCRC_adj_down = "None"

coef <- paste(column, tissue_type_A, 'vs', tissue_type_B, sep='_')
res <- DeseqResult(dds, column, coef, tissue_type_A, tissue_type_B,
                   lfc.Threshold, rpm.Threshold,
                   norm_adj_up,
                   norm_adj_down)
dict_sig_mirna[paste(coef, "up",   sep='_')] <- list(res$up_mirna)
dict_sig_mirna[paste(coef, "down", sep='_')] <- list(res$down_mirna)
res_res <- res$res
res_dict[coef] <- res_res
plotMA(res$res, alpha=0.05)
```

```
# Plot volcano plot
VolcanoPlot(res$res, coef, res$sig,
            res$up_mirna, res$down_mirna,
            norm_adj_up, norm_adj_down,
            pCRC_adj_up, pCRC_adj_down)
```

```
ExpressionPlot(res$res, res$rpm, coef, res$sig,
               tissue_type_A, tissue_type_B,
               res$up_mirna, res$down_mirna,
               norm_adj_up, norm_adj_down,
               pCRC_adj_up, pCRC_adj_down)
```

```
signature_mirnas <- SigList(res, dds, tissue_type_A, tissue_type_B, coef,
                            norm_adj_up, norm_adj_down, 
                            pCRC_adj_up, pCRC_adj_down)
# Print list upregulated miRNA
signature_mirnas$up_mirna
```

Upregulated in tissue.type\_tumor.colorect\_vs\_normal.colorect

| miRNA | LFC | lfcSE | FDR | RPM tumor.colorect | RPM normal.colorect | miRBase\_ID | Family | Seed | Chr | Cell-Type Specific | Norm Background | pCRC Background |
| --- | --- | --- | --- | --- | --- | --- | --- | --- | --- | --- | --- | --- |
| Hsa-Mir-17-P1a\_5p/P1b\_5p | 1.42 | 0.14 | 1.30e-22 | 900 | 225 | hsa-mir-17 | MIR-17 | AAAGUGC | chr13 | CD14+ Monocyte | na | na |
| Hsa-Mir-7-P1\_5p/P2\_5p/P3\_5p | 2.32 | 0.25 | 8.96e-18 | 199 | 22 | hsa-mir-7-1 | MIR-7 | GGAAGAC | chr9 | c(“Islet Cell”, “Neural”) | na | na |
| Hsa-Mir-223\_3p | 1.22 | 0.24 | 2.50e-06 | 619 | 177 | hsa-mir-223 | MIR-223 | GUCAGUU | chrX | c(“Dendritic Cell”, “Macrophage”) | na | na |
| Hsa-Mir-31\_5p | 4.32 | 0.33 | 1.40e-44 | 627 | 6 | hsa-mir-31 | MIR-31 | GGCAAGA | chr9 |  | na | na |
| Hsa-Mir-135-P3\_5p | 4.10 | 0.23 | 1.68e-69 | 155 | 4 | hsa-mir-135b | MIR-135 | AUGGCUU | chr1 |  | na | na |
| Hsa-Mir-224\_5p | 2.48 | 0.19 | 2.72e-36 | 395 | 43 | hsa-mir-224 | MIR-224 | AAGUCAC | chrX |  | na | na |
| Hsa-Mir-584\_5p | 1.94 | 0.21 | 2.16e-19 | 110 | 17 | hsa-mir-584 | MIR-584 | UAUGGUU | chr5 |  | na | na |
| Hsa-Mir-15-P1d\_5p | 1.87 | 0.21 | 1.15e-17 | 235 | 40 | hsa-mir-424 | MIR-15 | AGCAGCA | chrX |  | na | na |
| Hsa-Mir-96-P2\_5p | 1.84 | 0.17 | 6.44e-24 | 10417 | 1867 | hsa-mir-182 | MIR-96 | UUGGCAA | chr7 |  | na | na |
| Hsa-Mir-17-P3a\_5p | 1.62 | 0.16 | 2.85e-21 | 1513 | 322 | hsa-mir-20a | MIR-17 | AAAGUGC | chr13 |  | na | na |
| Hsa-Mir-96-P3\_5p | 1.53 | 0.23 | 1.96e-10 | 1063 | 224 | hsa-mir-183 | MIR-96 | AUGGCAC | chr7 |  | na | na |
| Hsa-Mir-19-P1\_3p | 1.47 | 0.17 | 1.43e-17 | 379 | 87 | hsa-mir-19a | MIR-19 | GUGCAAA | chr13 |  | na | na |
| Hsa-Mir-21\_5p | 1.35 | 0.13 | 1.01e-24 | 105735 | 28173 | hsa-mir-21 | MIR-21 | AGCUUAU | chr17 |  | na | na |
| Hsa-Mir-17-P2a\_5p | 1.33 | 0.19 | 6.62e-11 | 168 | 44 | hsa-mir-18a | MIR-17 | AAGGUGC | chr13 |  | na | na |
| Hsa-Mir-181-P2c\_5p | 1.33 | 0.16 | 1.40e-15 | 196 | 51 | hsa-mir-181d | MIR-181 | ACAUUCA | chr19 |  | na | na |
| Hsa-Mir-95-P2\_3p | 1.17 | 0.14 | 1.40e-15 | 142 | 41 | hsa-mir-421 | MIR-95 | UCAACAG | chrX |  | na | na |
| Hsa-Mir-19-P2a\_3p/P2b\_3p | 1.14 | 0.16 | 1.59e-12 | 1297 | 389 | hsa-mir-19b-1 | MIR-19 | GUGCAAA | chr13 |  | na | na |
| Hsa-Mir-181-P1c\_5p | 1.11 | 0.15 | 1.06e-12 | 1724 | 527 | hsa-mir-181c | MIR-181 | ACAUUCA | chr19 |  | na | na |
| Hsa-Mir-29-P2a\_3p/P2b\_3p | 1.08 | 0.17 | 5.43e-10 | 380 | 124 | hsa-mir-29b-1 | MIR-29 | AGCACCA | chr7 |  | na | na |
| Hsa-Mir-130-P2a\_3p | 1.05 | 0.15 | 8.09e-12 | 409 | 131 | hsa-mir-301a | MIR-130 | AGUGCAA | chr17 |  | na | na |
| Hsa-Mir-17-P3c\_5p | 0.97 | 0.12 | 1.00e-15 | 370 | 130 | hsa-mir-106b | MIR-17 | AAAGUGC | chr7 |  | na | na |
| Hsa-Mir-221-P2\_3p | 0.94 | 0.15 | 1.32e-09 | 1633 | 574 | hsa-mir-222 | MIR-221 | GCUACAU | chrX |  | na | na |
| Hsa-Mir-92-P1a\_3p/P1b\_3p | 0.90 | 0.16 | 5.66e-08 | 39320 | 13849 | hsa-mir-92a-1 | MIR-92 | AUUGCAC | chr13 |  | na | na |
| Hsa-Mir-17-P1c\_5p | 0.88 | 0.10 | 2.69e-17 | 2287 | 843 | hsa-mir-93 | MIR-17 | AAAGUGC | chr7 |  | na | na |
| Hsa-Mir-221-P1\_3p | 0.83 | 0.13 | 4.23e-09 | 2529 | 964 | hsa-mir-221 | MIR-221 | GCUACAU | chrX |  | na | na |
| Hsa-Mir-203\_3p | 0.74 | 0.20 | 9.13e-04 | 2307 | 922 | hsa-mir-203a | MIR-203 | UGAAAUG | chr14 |  | na | na |
| Hsa-Let-7-P2c2\_5p | 0.74 | 0.12 | 1.50e-08 | 5432 | 2218 | hsa-let-7i | LET-7 | GAGGUAG | chr12 |  | na | na |
| Hsa-Let-7-P2b3\_5p | 0.68 | 0.14 | 4.33e-06 | 2149 | 937 | hsa-mir-98 | LET-7 | GAGGUAG | chrX |  | na | na |
| Hsa-Mir-29-P1a\_3p | 0.62 | 0.12 | 1.40e-06 | 3667 | 1692 | hsa-mir-29a | MIR-29 | AGCACCA | chr7 |  | na | na |
| Hsa-Mir-92-P2c\_3p | 0.62 | 0.10 | 4.70e-09 | 3274 | 1443 | hsa-mir-25 | MIR-92 | AUUGCAC | chr7 |  | na | na |
| Hsa-Mir-92-P1c\_3p | 0.60 | 0.18 | 4.90e-03 | 1398 | 634 | hsa-mir-92b | MIR-92 | AUUGCAC | chr1 |  | na | na |
| Hsa-Mir-769\_5p | 0.59 | 0.11 | 1.18e-06 | 439 | 195 | hsa-mir-769 | MIR-769 | GAGACCU | chr19 |  | na | na |

```
# Number of upregulated miRNA
signature_mirnas$number_upregulated
```

```
## [1] 32
```

```
# Print list downregulated miRNA
signature_mirnas$down_mirna
```

Downregulated in tissue.type\_tumor.colorect\_vs\_normal.colorect

| miRNA | LFC | lfcSE | FDR | RPM tumor.colorect | RPM normal.colorect | miRBase\_ID | Family | Seed | Chr | Cell-Type Specific | Norm Background | pCRC Background |
| --- | --- | --- | --- | --- | --- | --- | --- | --- | --- | --- | --- | --- |
| Hsa-Mir-451\_5p | -1.19 | 0.26 | 3.69e-05 | 1565 | 2815 | hsa-mir-451a | MIR-451 | AACCGUU | chr17 | Red Blood Cell | na | na |
| Hsa-Mir-145\_5p | -1.97 | 0.22 | 1.63e-17 | 840 | 2832 | hsa-mir-145 | MIR-145 | UCCAGUU | chr5 | Mesenchymal | na | na |
| Hsa-Mir-143\_3p | -0.60 | 0.19 | 1.39e-03 | 148431 | 164812 | hsa-mir-143 | MIR-143 | GAGAUGA | chr5 | Mesenchymal | na | na |
| Hsa-Mir-150\_5p | -1.36 | 0.24 | 3.58e-07 | 280 | 586 | hsa-mir-150 | MIR-150 | CUCCCAA | chr19 | Lymphocyte | na | na |
| Hsa-Mir-192-P1\_5p/P2\_5p | -2.05 | 0.22 | 3.04e-19 | 13717 | 46010 | hsa-mir-192 | MIR-192 | UGACCUA | chr11 | Epithelial Cell | na | na |
| Hsa-Mir-133-P1\_3p/P2\_3p/P3\_3p | -1.98 | 0.26 | 6.98e-16 | 116 | 460 | hsa-mir-133a-2 | MIR-133 | UUGGUCC | chr20 | c(“Skeletal Myocyte”, “Stem Cell”) | na | na |
| Hsa-Mir-486\_5p | -1.18 | 0.25 | 2.02e-05 | 2013 | 3961 | hsa-mir-486-1 | MIR-486 | CCUGUAC | chr8 | c(“Platelet”, “Red Blood Cell”) | na | na |
| Hsa-Mir-375\_3p | -1.86 | 0.26 | 2.92e-12 | 5263 | 17215 | hsa-mir-375 | MIR-375 | UUGUUCG | chr2 | c(“Epithelial Cell”, “Islet Cell”, “Neural”) | na | na |
| Hsa-Mir-126\_5p | -0.72 | 0.15 | 1.61e-05 | 3455 | 4230 | hsa-mir-126 | MIR-126 | AUUAUUA | chr9 | c(“Endothelial Cell”, “Platelet”) | na | na |
| Hsa-Mir-342\_3p | -1.26 | 0.18 | 1.59e-10 | 145 | 260 | hsa-mir-342 | MIR-342 | CUCACAC | chr14 | c(“Dendritic Cell”, “Lymphocyte”, “Macrophage”) | na | na |
| Hsa-Mir-147\_3p | -1.92 | 0.23 | 8.04e-17 | 33 | 110 | hsa-mir-147b | MIR-147 | UGUGCGG | chr15 |  | na | na |
| Hsa-Mir-15-P2c\_5p | -1.76 | 0.17 | 6.42e-23 | 257 | 672 | hsa-mir-195 | MIR-15 | AGCAGCA | chr17 |  | na | na |
| Hsa-Mir-378\_3p | -1.72 | 0.14 | 8.99e-34 | 6426 | 14834 | hsa-mir-378a | MIR-378 | CUGGACU | chr5 |  | na | na |
| Hsa-Mir-15-P1c\_5p | -1.62 | 0.16 | 5.86e-22 | 143 | 337 | hsa-mir-497 | MIR-15 | AGCAGCA | chr17 |  | na | na |
| Hsa-Mir-190-P1\_5p | -1.47 | 0.18 | 1.43e-15 | 140 | 294 | hsa-mir-190a | MIR-190 | GAUAUGU | chr15 |  | na | na |
| Hsa-Mir-194-P1\_5p/P2\_5p | -1.31 | 0.19 | 3.24e-11 | 6167 | 11619 | hsa-mir-194-2 | MIR-194 | GUAACAG | chr11 |  | na | na |
| Hsa-Mir-30-P1a\_5p | -1.21 | 0.16 | 7.30e-12 | 3165 | 5150 | hsa-mir-30a | MIR-30 | GUAAACA | chr6 |  | na | na |
| Hsa-Mir-26-P3\_5p | -1.21 | 0.14 | 3.12e-16 | 3467 | 5898 | hsa-mir-26a-1 | MIR-26 | UCAAGUA | chr3 |  | na | na |
| Hsa-Mir-338-P1\_3p | -1.18 | 0.22 | 8.59e-07 | 98 | 166 | hsa-mir-338 | MIR-338 | CCAGCAU | chr17 |  | na | na |
| Hsa-Mir-26-P1\_5p/P2\_5p | -1.15 | 0.10 | 4.50e-29 | 35584 | 56268 | hsa-mir-26b | MIR-26 | UCAAGUA | chr2 |  | na | na |
| Hsa-Mir-10-P1b\_5p | -1.11 | 0.25 | 4.63e-06 | 39286 | 75067 | hsa-mir-10b | MIR-10 | ACCCUGU | chr2 |  | na | na |
| Hsa-Mir-192-P1\_5p | -1.11 | 0.20 | 2.37e-08 | 132000 | 212293 | hsa-mir-192 | MIR-192 | UGACCUA | chr11 |  | na | na |
| Hsa-Mir-148-P2\_3p | -1.04 | 0.21 | 5.54e-06 | 137 | 210 | hsa-mir-152 | MIR-148 | CAGUGCA | chr17 |  | na | na |
| Hsa-Mir-15-P2a\_5p/P2b\_5p | -1.02 | 0.12 | 8.21e-15 | 4522 | 6926 | hsa-mir-16-1 | MIR-15 | AGCAGCA | chr13 |  | na | na |
| Hsa-Mir-10-P3b\_5p | -0.97 | 0.19 | 4.09e-06 | 1649 | 2392 | hsa-mir-125a | MIR-10 | CCCUGAG | chr19 |  | na | na |
| Hsa-Mir-28-P1\_3p | -0.95 | 0.13 | 6.44e-12 | 2915 | 3956 | hsa-mir-28 | MIR-28 | ACUAGAU | chr3 |  | na | na |
| Hsa-Mir-1307\_5p | -0.95 | 0.18 | 2.27e-06 | 492 | 713 | hsa-mir-1307 | MIR-1307 | CGACCGG | chr10 |  | na | na |
| Hsa-Mir-29-P1b\_3p | -0.93 | 0.15 | 1.11e-09 | 331 | 464 | hsa-mir-29c | MIR-29 | AGCACCA | chr1 |  | na | na |
| Hsa-Mir-10-P3c\_5p | -0.80 | 0.21 | 1.44e-03 | 239 | 326 | hsa-mir-125b-2 | MIR-10 | CCCUGAG | chr21 |  | na | na |
| Hsa-Mir-574\_3p | -0.78 | 0.14 | 1.83e-07 | 240 | 305 | hsa-mir-574 | MIR-574 | ACGCUCA | chr4 |  | na | na |
| Hsa-Mir-8-P1b\_3p | -0.69 | 0.21 | 2.87e-03 | 7518 | 9180 | hsa-mir-141 | MIR-8 | AACACUG | chr12 |  | na | na |
| Hsa-Mir-191\_5p | -0.68 | 0.16 | 1.46e-04 | 13630 | 14608 | hsa-mir-191 | MIR-191 | AACGGAA | chr3 |  | na | na |
| Hsa-Mir-362-P3\_3p | -0.62 | 0.25 | 1.92e-02 | 108 | 103 | hsa-mir-501 | MIR-362 | AUGCACC | chrX |  | na | na |
| Hsa-Mir-142\_5p | -0.60 | 0.19 | 4.90e-03 | 3453 | 3817 | hsa-mir-142 | MIR-142 | AUAAAGU | chr17 |  | na | na |
| Hsa-Mir-154-P9\_3p | -0.59 | 0.15 | 1.81e-04 | 253 | 265 | hsa-mir-381 | MIR-154 | AUACAAG | chr14 |  | na | na |

```
# Number of downregulated miRNA
signature_mirnas$number_downregulated
```

```
## [1] 35
```

```
ref <- 'tumor.colorect'
dds <- DeseqObject(design, countdata, sampleinfo, "None", "None", ref)
#
```

```
# #datasets in total
dim(dds[, colData(dds)$type.tissue == 'pCRC'])
```

```
## [1] 389 120
```

```
dim(dds[, colData(dds)$type.tissue == 'mLi'])
```

```
## [1] 389  35
```

```
dim(dds[, colData(dds)$type.tissue == 'mLu'])
```

```
## [1] 389  28
```

```
dim(dds[, colData(dds)$type.tissue == 'nCR'])
```

```
## [1] 389  25
```

```
dim(dds[, colData(dds)$type.tissue == 'nLi'])
```

```
## [1] 389  20
```

```
dim(dds[, colData(dds)$type.tissue == 'nLu'])
```

```
## [1] 389  10
```

```
dim(dds[, colData(dds)$type.tissue == 'PM'])
```

```
## [1] 389  30
```

```
# #datasets for Fromm
dim(dds[, colData(dds)$type.tissue == 'pCRC' & colData(dds)$paper == 'fromm'])
```

```
## [1] 389   3
```

```
dim(dds[, colData(dds)$type.tissue == 'mLi' & colData(dds)$paper == 'fromm'])
```

```
## [1] 389  19
```

```
dim(dds[, colData(dds)$type.tissue == 'mLu' & colData(dds)$paper == 'fromm'])
```

```
## [1] 389  24
```

```
dim(dds[, colData(dds)$type.tissue == 'nCR' & colData(dds)$paper == 'fromm'])
```

```
## [1] 389   3
```

```
dim(dds[, colData(dds)$type.tissue == 'nLi' & colData(dds)$paper == 'fromm'])
```

```
## [1] 389   8
```

```
dim(dds[, colData(dds)$type.tissue == 'nLu' & colData(dds)$paper == 'fromm'])
```

```
## [1] 389   7
```

```
dim(dds[, colData(dds)$type.tissue == 'PM' & colData(dds)$paper == 'fromm'])
```

```
## [1] 389  18
```

```
# #datasets for Schee
dim(dds[, colData(dds)$type.tissue == 'pCRC' & colData(dds)$paper == 'schee'])
```

```
## [1] 389  83
```

```
dim(dds[, colData(dds)$type.tissue == 'mLi' & colData(dds)$paper == 'schee'])
```

```
## [1] 389   0
```

```
dim(dds[, colData(dds)$type.tissue == 'mLu' & colData(dds)$paper == 'schee'])
```

```
## [1] 389   0
```

```
dim(dds[, colData(dds)$type.tissue == 'nCR' & colData(dds)$paper == 'schee'])
```

```
## [1] 389   0
```

```
dim(dds[, colData(dds)$type.tissue == 'nLi' & colData(dds)$paper == 'schee'])
```

```
## [1] 389   0
```

```
dim(dds[, colData(dds)$type.tissue == 'nLu' & colData(dds)$paper == 'schee'])
```

```
## [1] 389   0
```

```
dim(dds[, colData(dds)$type.tissue == 'PM' & colData(dds)$paper == 'schee'])
```

```
## [1] 389   0
```

```
# #datasets for Schee
dim(dds[, colData(dds)$type.tissue == 'pCRC' & colData(dds)$paper == 'neerincx'])
```

```
## [1] 389  34
```

```
dim(dds[, colData(dds)$type.tissue == 'mLi' & colData(dds)$paper == 'neerincx'])
```

```
## [1] 389  16
```

```
dim(dds[, colData(dds)$type.tissue == 'mLu' & colData(dds)$paper == 'neerincx'])
```

```
## [1] 389   4
```

```
dim(dds[, colData(dds)$type.tissue == 'nCR' & colData(dds)$paper == 'neerincx'])
```

```
## [1] 389  22
```

```
dim(dds[, colData(dds)$type.tissue == 'nLi' & colData(dds)$paper == 'neerincx'])
```

```
## [1] 389   9
```

```
dim(dds[, colData(dds)$type.tissue == 'nLu' & colData(dds)$paper == 'neerincx'])
```

```
## [1] 389   3
```

```
dim(dds[, colData(dds)$type.tissue == 'PM' & colData(dds)$paper == 'neerincx'])
```

```
## [1] 389  12
```

```
# #datasets for Schee
dim(dds[, colData(dds)$type.tissue == 'pCRC' & colData(dds)$paper == 'selitsky'])
```

```
## [1] 389   0
```

```
dim(dds[, colData(dds)$type.tissue == 'mLi' & colData(dds)$paper == 'selitsky'])
```

```
## [1] 389   0
```

```
dim(dds[, colData(dds)$type.tissue == 'mLu' & colData(dds)$paper == 'selitsky'])
```

```
## [1] 389   0
```

```
dim(dds[, colData(dds)$type.tissue == 'nCR' & colData(dds)$paper == 'selitsky'])
```

```
## [1] 389   0
```

```
dim(dds[, colData(dds)$type.tissue == 'nLi' & colData(dds)$paper == 'selitsky'])
```

```
## [1] 389   3
```

```
dim(dds[, colData(dds)$type.tissue == 'nLu' & colData(dds)$paper == 'selitsky'])
```

```
## [1] 389   0
```

```
dim(dds[, colData(dds)$type.tissue == 'PM' & colData(dds)$paper == 'selitsky'])
```

```
## [1] 389   0
```

```
plotDispEsts(dds)
```

#### pCRC vs nLi

```
column='tissue.type'
tissue_type_A <- 'normal.liver'
tissue_type_B <- 'tumor.colorect'
norm_adj_up       = "None"
norm_adj_down     = "None"
pCRC_adj_up   = "None"
pCRC_adj_down = "None"

coef <- paste(column, tissue_type_A, 'vs', tissue_type_B, sep='_')
res <- DeseqResult(dds, column, coef, tissue_type_A, tissue_type_B,
                   lfc.Threshold, rpm.Threshold,
                   norm_adj_up,
                   norm_adj_down)
dict_sig_mirna[paste(coef, "up",   sep='_')] <- list(res$up_mirna)
dict_sig_mirna[paste(coef, "down", sep='_')] <- list(res$down_mirna)
res_res <- res$res
res_dict[coef] <- res_res
plotMA(res$res, alpha=0.05)
```

```
# Plot volcano plot
VolcanoPlot(res$res, coef, res$sig,
            res$up_mirna, res$down_mirna,
            norm_adj_up, norm_adj_down,
            pCRC_adj_up, pCRC_adj_down)
```

```
ExpressionPlot(res$res, res$rpm, coef, res$sig,
               tissue_type_A, tissue_type_B,
               res$up_mirna, res$down_mirna,
               norm_adj_up, norm_adj_down,
               pCRC_adj_up, pCRC_adj_down)
```

```
signature_mirnas <- SigList(res, dds, tissue_type_A, tissue_type_B, coef,
                            norm_adj_up, norm_adj_down, 
                            pCRC_adj_up, pCRC_adj_down)
# Print list upregulated miRNA
signature_mirnas$up_mirna
```

Upregulated in tissue.type\_normal.liver\_vs\_tumor.colorect

| miRNA | LFC | lfcSE | FDR | RPM normal.liver | RPM tumor.colorect | miRBase\_ID | Family | Seed | Chr | Cell-Type Specific | Norm Background | pCRC Background |
| --- | --- | --- | --- | --- | --- | --- | --- | --- | --- | --- | --- | --- |
| Hsa-Mir-204-P1\_5p | 1.03 | 0.46 | 1.25e-04 | 190 | 62 | hsa-mir-204 | MIR-204 | UCCCUUU | chr9 | Retinal Epithelial Cell | na | na |
| Hsa-Mir-335\_5p | 0.81 | 0.14 | 1.89e-08 | 205 | 150 | hsa-mir-335 | MIR-335 | CAAGAGC | chr7 | Retinal Epithelial Cell | na | na |
| Hsa-Mir-144\_5p | 2.22 | 0.28 | 3.68e-15 | 206 | 57 | hsa-mir-144 | MIR-144 | GAUAUCA | chr17 | Red Blood Cell | na | na |
| Hsa-Mir-451\_5p | 1.48 | 0.32 | 7.08e-06 | 3358 | 1565 | hsa-mir-451a | MIR-451 | AACCGUU | chr17 | Red Blood Cell | na | na |
| Hsa-Mir-150\_5p | 2.07 | 0.29 | 5.24e-13 | 1053 | 280 | hsa-mir-150 | MIR-150 | CUCCCAA | chr19 | Lymphocyte | na | na |
| Hsa-Mir-122\_5p | 11.53 | 0.78 | 4.04e-55 | 148842 | 9 | hsa-mir-122 | MIR-122 | GGAGUGU | chr18 | Hepatocyte | na | na |
| Hsa-Mir-192-P1\_5p/P2\_5p | 1.31 | 0.27 | 2.62e-06 | 29546 | 13717 | hsa-mir-192 | MIR-192 | UGACCUA | chr11 | Epithelial Cell | na | na |
| Hsa-Mir-15-P1a\_5p | 0.62 | 0.12 | 1.17e-06 | 555 | 453 | hsa-mir-15a | MIR-15 | AGCAGCA | chr13 | CD14+ Monocyte | na | na |
| Hsa-Mir-486\_5p | 2.48 | 0.30 | 8.95e-16 | 10617 | 2013 | hsa-mir-486-1 | MIR-486 | CCUGUAC | chr8 | c(“Platelet”, “Red Blood Cell”) | na | na |
| Hsa-Mir-126\_5p | 1.95 | 0.17 | 7.32e-30 | 11148 | 3455 | hsa-mir-126 | MIR-126 | AUUAUUA | chr9 | c(“Endothelial Cell”, “Platelet”) | na | na |
| Hsa-Mir-342\_3p | 1.05 | 0.21 | 5.60e-07 | 254 | 145 | hsa-mir-342 | MIR-342 | CUCACAC | chr14 | c(“Dendritic Cell”, “Lymphocyte”, “Macrophage”) | na | na |
| Hsa-Mir-885\_5p | 9.48 | 0.68 | 8.47e-41 | 510 | 0 | hsa-mir-885 | MIR-885 | CCAUUAC | chr3 |  | na | na |
| Hsa-Mir-139\_5p | 4.20 | 0.21 | 6.49e-84 | 174 | 11 | hsa-mir-139 | MIR-139 | CUACAGU | chr11 |  | na | na |
| Hsa-Let-7-P1c\_5p | 3.25 | 0.23 | 3.14e-43 | 4945 | 606 | hsa-let-7c | LET-7 | GAGGUAG | chr21 |  | na | na |
| Hsa-Mir-10-P2c\_5p | 3.24 | 0.28 | 2.36e-31 | 1202 | 142 | hsa-mir-99a | MIR-10 | ACCCGUA | chr21 |  | na | na |
| Hsa-Mir-10-P3c\_5p | 2.87 | 0.25 | 2.44e-30 | 1620 | 239 | hsa-mir-125b-2 | MIR-10 | CCCUGAG | chr21 |  | na | na |
| Hsa-Mir-30-P1a\_5p | 2.51 | 0.19 | 3.24e-40 | 15175 | 3165 | hsa-mir-30a | MIR-30 | GUAAACA | chr6 |  | na | na |
| Hsa-Mir-193-P1a\_5p | 2.48 | 0.23 | 3.82e-25 | 298 | 65 | hsa-mir-193a | MIR-193 | GGGUCUU | chr17 |  | na | na |
| Hsa-Mir-193-P2a\_3p/P2b\_3p | 2.33 | 0.19 | 1.61e-32 | 219 | 56 | hsa-mir-365b | MIR-193 | AAUGCCC | chr17 |  | na | na |
| Hsa-Mir-10-P3a\_5p | 2.29 | 0.26 | 3.08e-18 | 611 | 145 | hsa-mir-125b-1 | MIR-10 | CCCUGAG | chr11 |  | na | na |
| Hsa-Mir-193-P1b\_3p | 2.17 | 0.19 | 2.30e-29 | 514 | 145 | hsa-mir-193b | MIR-193 | ACUGGCC | chr16 |  | na | na |
| Hsa-Mir-455\_5p | 2.10 | 0.15 | 6.57e-45 | 279 | 83 | hsa-mir-455 | MIR-455 | AUGUGCC | chr9 |  | na | na |
| Hsa-Mir-574\_3p | 1.99 | 0.16 | 1.44e-34 | 745 | 240 | hsa-mir-574 | MIR-574 | ACGCUCA | chr4 |  | na | na |
| Hsa-Mir-101-P1\_3p/P2\_3p | 1.98 | 0.15 | 6.40e-41 | 14649 | 4638 | hsa-mir-101-1 | MIR-101 | UACAGUA | chr1 |  | na | na |
| Hsa-Mir-148-P1\_3p | 1.86 | 0.20 | 1.75e-19 | 105086 | 34704 | hsa-mir-148a | MIR-148 | CAGUGCA | chr7 |  | na | na |
| Hsa-Mir-423\_5p | 1.83 | 0.20 | 1.11e-18 | 1198 | 433 | hsa-mir-423 | MIR-423 | GAGGGGC | chr17 |  | na | na |
| Hsa-Mir-22-P1a\_3p | 1.79 | 0.14 | 1.08e-36 | 78932 | 29018 | hsa-mir-22 | MIR-22 | AGCUGCC | chr17 |  | na | na |
| Hsa-Mir-744\_5p | 1.79 | 0.17 | 1.28e-23 | 165 | 61 | hsa-mir-744 | MIR-744 | GCGGGGC | chr17 |  | na | na |
| Hsa-Mir-30-P2a\_5p/P2b\_5p/P2c\_5p | 1.45 | 0.12 | 7.16e-32 | 5891 | 2793 | hsa-mir-30c-2 | MIR-30 | GUAAACA | chr6 |  | na | na |
| Hsa-Mir-130-P1a\_3p | 1.43 | 0.16 | 1.39e-17 | 907 | 421 | hsa-mir-130a | MIR-130 | AGUGCAA | chr11 |  | na | na |
| Hsa-Mir-378\_3p | 1.35 | 0.16 | 1.07e-16 | 12736 | 6426 | hsa-mir-378a | MIR-378 | CUGGACU | chr5 |  | na | na |
| Hsa-Mir-148-P2\_3p | 1.34 | 0.25 | 1.57e-07 | 289 | 137 | hsa-mir-152 | MIR-148 | CAGUGCA | chr17 |  | na | na |
| Hsa-Mir-197\_3p | 1.28 | 0.15 | 5.68e-16 | 344 | 181 | hsa-mir-197 | MIR-197 | UCACCAC | chr1 |  | na | na |
| Hsa-Mir-15-P1c\_5p | 1.25 | 0.19 | 3.90e-11 | 276 | 143 | hsa-mir-497 | MIR-15 | AGCAGCA | chr17 |  | na | na |
| Hsa-Mir-26-P3\_5p | 1.24 | 0.16 | 2.22e-14 | 6787 | 3467 | hsa-mir-26a-1 | MIR-26 | UCAAGUA | chr3 |  | na | na |
| Hsa-Mir-15-P2c\_5p | 1.19 | 0.20 | 1.78e-09 | 496 | 257 | hsa-mir-195 | MIR-15 | AGCAGCA | chr17 |  | na | na |
| Hsa-Mir-331\_3p | 1.16 | 0.21 | 6.75e-08 | 110 | 60 | hsa-mir-331 | MIR-331 | CCCCUGG | chr12 |  | na | na |
| Hsa-Mir-26-P1\_5p/P2\_5p | 1.10 | 0.11 | 1.13e-22 | 62274 | 35584 | hsa-mir-26b | MIR-26 | UCAAGUA | chr2 |  | na | na |
| Hsa-Mir-10-P3b\_5p | 1.09 | 0.23 | 2.99e-06 | 3040 | 1649 | hsa-mir-125a | MIR-10 | CCCUGAG | chr19 |  | na | na |
| Hsa-Mir-30-P1b\_5p | 1.08 | 0.12 | 5.44e-19 | 11240 | 6833 | hsa-mir-30e | MIR-30 | GUAAACA | chr1 |  | na | na |
| Hsa-Mir-340\_5p | 1.05 | 0.13 | 1.20e-15 | 1030 | 657 | hsa-mir-340 | MIR-340 | UAUAAAG | chr5 |  | na | na |
| Hsa-Mir-27-P1\_3p/P2\_3p | 1.04 | 0.12 | 4.13e-17 | 50009 | 29444 | hsa-mir-27a | MIR-27 | UCACAGU | chr19 |  | na | na |
| Hsa-Mir-10-P2a\_5p | 1.00 | 0.27 | 4.80e-04 | 2634 | 1724 | hsa-mir-100 | MIR-10 | ACCCGUA | chr11 |  | na | na |
| Hsa-Mir-29-P1b\_3p | 0.98 | 0.17 | 1.55e-08 | 516 | 331 | hsa-mir-29c | MIR-29 | AGCACCA | chr1 |  | na | na |
| Hsa-Mir-154-P23\_3p | 0.94 | 0.16 | 3.85e-08 | 222 | 148 | hsa-mir-654 | MIR-154 | AUGUCUG | chr14 |  | na | na |
| Hsa-Mir-15-P2a\_5p/P2b\_5p | 0.87 | 0.14 | 9.10e-10 | 6884 | 4522 | hsa-mir-16-1 | MIR-15 | AGCAGCA | chr13 |  | na | na |
| Hsa-Mir-30-P1c\_5p | 0.84 | 0.13 | 9.49e-10 | 13680 | 9703 | hsa-mir-30d | MIR-30 | GUAAACA | chr8 |  | na | na |
| Hsa-Mir-154-P13\_5p | 0.78 | 0.18 | 2.94e-05 | 442 | 328 | hsa-mir-411 | MIR-154 | AGUAGAC | chr14 |  | na | na |
| Hsa-Mir-28-P2\_5p | 0.76 | 0.15 | 1.01e-06 | 3098 | 2244 | hsa-mir-151a | MIR-28 | CGAGGAG | chr8 |  | na | na |
| Hsa-Mir-92-P1a\_3p/P1b\_3p | 0.71 | 0.18 | 2.33e-04 | 51173 | 39320 | hsa-mir-92a-1 | MIR-92 | AUUGCAC | chr13 |  | na | na |
| Hsa-Mir-136\_3p | 0.65 | 0.18 | 4.46e-04 | 208 | 173 | hsa-mir-136 | MIR-136 | AUCAUCG | chr14 |  | na | na |

```
# Number of upregulated miRNA
signature_mirnas$number_upregulated
```

```
## [1] 51
```

```
# Print list downregulated miRNA
signature_mirnas$down_mirna
```

Downregulated in tissue.type\_normal.liver\_vs\_tumor.colorect

| miRNA | LFC | lfcSE | FDR | RPM normal.liver | RPM tumor.colorect | miRBase\_ID | Family | Seed | Chr | Cell-Type Specific | Norm Background | pCRC Background |
| --- | --- | --- | --- | --- | --- | --- | --- | --- | --- | --- | --- | --- |
| Hsa-Mir-143\_3p | -1.72 | 0.23 | 4.38e-14 | 37572 | 148431 | hsa-mir-143 | MIR-143 | GAGAUGA | chr5 | Mesenchymal | na | na |
| Hsa-Mir-24-P1\_3p/P2\_3p | -0.77 | 0.16 | 8.36e-06 | 319 | 626 | hsa-mir-24-2 | MIR-24 | GGCUCAG | chr19 | Macrophage | na | na |
| Hsa-Mir-8-P2b\_3p | -6.07 | 0.19 | 3.90e-210 | 31 | 2727 | hsa-mir-200c | MIR-8 | AAUACUG | chr12 | Epithelial Cell | na | na |
| Hsa-Mir-8-P2a\_3p | -4.36 | 0.19 | 2.71e-109 | 369 | 10330 | hsa-mir-200b | MIR-8 | AAUACUG | chr1 | Epithelial Cell | na | na |
| Hsa-Mir-17-P1a\_5p/P1b\_5p | -1.32 | 0.16 | 4.02e-16 | 278 | 900 | hsa-mir-17 | MIR-17 | AAAGUGC | chr13 | CD14+ Monocyte | na | na |
| Hsa-Mir-15-P1b\_5p | -0.92 | 0.13 | 1.30e-11 | 114 | 273 | hsa-mir-15b | MIR-15 | AGCAGCA | chr3 | CD14+ Monocyte | na | na |
| Hsa-Mir-133-P1\_3p/P2\_3p/P3\_3p | -2.03 | 0.32 | 1.91e-10 | 22 | 116 | hsa-mir-133a-2 | MIR-133 | UUGGUCC | chr20 | c(“Skeletal Myocyte”, “Stem Cell”) | na | na |
| Hsa-Mir-155\_5p | -1.44 | 0.21 | 1.42e-10 | 113 | 387 | hsa-mir-155 | MIR-155 | UAAUGCU | chr21 | c(“Lymphocyte”, “Macrophage”) | na | na |
| Hsa-Mir-7-P1\_5p/P2\_5p/P3\_5p | -5.51 | 0.32 | 2.03e-63 | 3 | 199 | hsa-mir-7-1 | MIR-7 | GGAAGAC | chr9 | c(“Islet Cell”, “Neural”) | na | na |
| Hsa-Mir-375\_3p | -0.84 | 0.32 | 2.21e-02 | 2553 | 5263 | hsa-mir-375 | MIR-375 | UUGUUCG | chr2 | c(“Epithelial Cell”, “Islet Cell”, “Neural”) | na | na |
| Hsa-Mir-196-P1\_5p/P2\_5p | -6.71 | 0.30 | 2.31e-108 | 2 | 319 | hsa-mir-196a-1 | MIR-196 | AGGUAGU | chr17 |  | na | na |
| Hsa-Mir-135-P3\_5p | -6.12 | 0.31 | 3.71e-82 | 2 | 155 | hsa-mir-135b | MIR-135 | AUGGCUU | chr1 |  | na | na |
| Hsa-Mir-196-P3\_5p | -6.11 | 0.32 | 1.17e-77 | 9 | 828 | hsa-mir-196b | MIR-196 | AGGUAGU | chr7 |  | na | na |
| Hsa-Mir-31\_5p | -5.76 | 0.45 | 3.20e-37 | 7 | 627 | hsa-mir-31 | MIR-31 | GGCAAGA | chr9 |  | na | na |
| Hsa-Mir-577\_5p | -5.70 | 0.29 | 1.37e-80 | 2 | 142 | hsa-mir-577 | MIR-577 | UAGAUAA | chr4 |  | na | na |
| Hsa-Mir-8-P1b\_3p | -5.35 | 0.25 | 1.29e-93 | 154 | 7518 | hsa-mir-141 | MIR-8 | AACACUG | chr12 |  | na | na |
| Hsa-Mir-96-P3\_5p | -5.08 | 0.27 | 2.84e-74 | 27 | 1063 | hsa-mir-183 | MIR-96 | AUGGCAC | chr7 |  | na | na |
| Hsa-Mir-8-P3a\_3p | -4.60 | 0.20 | 5.55e-118 | 68 | 2303 | hsa-mir-429 | MIR-8 | AAUACUG | chr1 |  | na | na |
| Hsa-Mir-96-P2\_5p | -4.47 | 0.20 | 4.46e-107 | 370 | 10417 | hsa-mir-182 | MIR-96 | UUGGCAA | chr7 |  | na | na |
| Hsa-Mir-10-P1b\_5p | -3.68 | 0.31 | 1.37e-31 | 2704 | 39286 | hsa-mir-10b | MIR-10 | ACCCUGU | chr2 |  | na | na |
| Hsa-Mir-8-P1a\_3p | -3.64 | 0.19 | 9.42e-76 | 108 | 1772 | hsa-mir-200a | MIR-8 | AACACUG | chr1 |  | na | na |
| Hsa-Mir-203\_3p | -3.42 | 0.24 | 2.77e-45 | 171 | 2307 | hsa-mir-203a | MIR-203 | UGAAAUG | chr14 |  | na | na |
| Hsa-Mir-221-P1\_3p | -3.13 | 0.15 | 5.83e-92 | 217 | 2529 | hsa-mir-221 | MIR-221 | GCUACAU | chrX |  | na | na |
| Hsa-Mir-224\_5p | -2.99 | 0.22 | 2.60e-39 | 36 | 395 | hsa-mir-224 | MIR-224 | AAGUCAC | chrX |  | na | na |
| Hsa-Mir-221-P2\_3p | -2.81 | 0.17 | 1.32e-61 | 173 | 1633 | hsa-mir-222 | MIR-221 | GCUACAU | chrX |  | na | na |
| Hsa-Mir-181-P2c\_5p | -2.59 | 0.19 | 7.58e-42 | 26 | 196 | hsa-mir-181d | MIR-181 | ACAUUCA | chr19 |  | na | na |
| Hsa-Mir-181-P1c\_5p | -2.35 | 0.17 | 3.48e-42 | 261 | 1724 | hsa-mir-181c | MIR-181 | ACAUUCA | chr19 |  | na | na |
| Hsa-Mir-584\_5p | -2.25 | 0.25 | 4.29e-18 | 18 | 110 | hsa-mir-584 | MIR-584 | UAUGGUU | chr5 |  | na | na |
| Hsa-Mir-190-P1\_5p | -2.24 | 0.22 | 4.84e-24 | 23 | 140 | hsa-mir-190a | MIR-190 | GAUAUGU | chr15 |  | na | na |
| Hsa-Mir-17-P2a\_5p | -2.22 | 0.23 | 1.80e-21 | 26 | 168 | hsa-mir-18a | MIR-17 | AAGGUGC | chr13 |  | na | na |
| Hsa-Mir-21\_5p | -2.10 | 0.14 | 5.68e-48 | 19240 | 105735 | hsa-mir-21 | MIR-21 | AGCUUAU | chr17 |  | na | na |
| Hsa-Mir-92-P1c\_3p | -1.90 | 0.21 | 3.08e-18 | 303 | 1398 | hsa-mir-92b | MIR-92 | AUUGCAC | chr1 |  | na | na |
| Hsa-Mir-10-P1a\_5p | -1.79 | 0.25 | 2.08e-11 | 25828 | 97123 | hsa-mir-10a | MIR-10 | ACCCUGU | chr17 |  | na | na |
| Hsa-Mir-146-P1\_5p | -1.77 | 0.29 | 4.42e-09 | 351 | 1453 | hsa-mir-146a | MIR-146 | GAGAACU | chr5 |  | na | na |
| Hsa-Mir-130-P2a\_3p | -1.75 | 0.17 | 1.02e-24 | 92 | 409 | hsa-mir-301a | MIR-130 | AGUGCAA | chr17 |  | na | na |
| Hsa-Mir-29-P2a\_3p/P2b\_3p | -1.48 | 0.19 | 9.97e-14 | 107 | 380 | hsa-mir-29b-1 | MIR-29 | AGCACCA | chr7 |  | na | na |
| Hsa-Mir-17-P3c\_5p | -1.47 | 0.13 | 8.74e-29 | 100 | 370 | hsa-mir-106b | MIR-17 | AAAGUGC | chr7 |  | na | na |
| Hsa-Mir-17-P1c\_5p | -1.45 | 0.11 | 7.11e-37 | 647 | 2287 | hsa-mir-93 | MIR-17 | AAAGUGC | chr7 |  | na | na |
| Hsa-Mir-95-P2\_3p | -1.42 | 0.16 | 7.76e-18 | 40 | 142 | hsa-mir-421 | MIR-95 | UCAACAG | chrX |  | na | na |
| Hsa-Mir-17-P3a\_5p | -1.41 | 0.19 | 1.99e-13 | 437 | 1513 | hsa-mir-20a | MIR-17 | AAAGUGC | chr13 |  | na | na |
| Hsa-Mir-188-P2\_5p | -1.33 | 0.15 | 3.22e-17 | 279 | 914 | hsa-mir-532 | MIR-188 | AUGCCUU | chrX |  | na | na |
| Hsa-Let-7-P2c2\_5p | -1.26 | 0.14 | 4.74e-19 | 1738 | 5432 | hsa-let-7i | LET-7 | GAGGUAG | chr12 |  | na | na |
| Hsa-Let-7-P2b3\_5p | -1.22 | 0.15 | 7.97e-15 | 760 | 2149 | hsa-mir-98 | LET-7 | GAGGUAG | chrX |  | na | na |
| Hsa-Mir-210\_3p | -1.16 | 0.23 | 5.10e-06 | 106 | 291 | hsa-mir-210 | MIR-210 | UGUGCGU | chr11 |  | na | na |
| Hsa-Mir-425\_5p | -1.15 | 0.15 | 2.77e-13 | 247 | 700 | hsa-mir-425 | MIR-425 | AUGACAC | chr3 |  | na | na |
| Hsa-Mir-19-P1\_3p | -1.12 | 0.19 | 1.87e-08 | 134 | 379 | hsa-mir-19a | MIR-19 | GUGCAAA | chr13 |  | na | na |
| Hsa-Mir-132-P1\_3p | -1.06 | 0.17 | 6.29e-10 | 61 | 165 | hsa-mir-132 | MIR-132 | AACAGUC | chr17 |  | na | na |
| Hsa-Mir-130-P4a\_3p | -0.99 | 0.10 | 5.70e-21 | 70 | 177 | hsa-mir-454 | MIR-130 | AGUGCAA | chr17 |  | na | na |
| Hsa-Mir-652\_3p | -0.91 | 0.13 | 2.45e-11 | 44 | 106 | hsa-mir-652 | MIR-652 | AUGGCGC | chrX |  | na | na |
| Hsa-Mir-769\_5p | -0.81 | 0.13 | 9.49e-10 | 190 | 439 | hsa-mir-769 | MIR-769 | GAGACCU | chr19 |  | na | na |
| Hsa-Mir-362-P2\_3p/P4\_3p | -0.69 | 0.17 | 7.46e-05 | 211 | 441 | hsa-mir-500a | MIR-362 | UGCACCU | chrX |  | na | na |
| Hsa-Mir-181-P2a\_5p/P2b\_5p | -0.67 | 0.15 | 1.77e-05 | 611 | 1251 | hsa-mir-181b-1 | MIR-181 | ACAUUCA | chr1 |  | na | na |
| Hsa-Mir-17-P3c\_3p | -0.64 | 0.14 | 5.82e-06 | 105 | 212 | hsa-mir-106b | MIR-17 | AAAGUGC | chr7 |  | na | na |
| Hsa-Mir-19-P2a\_3p/P2b\_3p | -0.60 | 0.18 | 1.90e-03 | 677 | 1297 | hsa-mir-19b-1 | MIR-19 | GUGCAAA | chr13 |  | na | na |

```
# Number of downregulated miRNA
signature_mirnas$number_downregulated
```

```
## [1] 54
```

#### pCRC vs nLu

```
column='tissue.type'
tissue_type_A <- 'normal.lung'
tissue_type_B <- 'tumor.colorect'
norm_adj_up       = "None"
norm_adj_down     = "None"
pCRC_adj_up   = "None"
pCRC_adj_down = "None"

coef <- paste(column, tissue_type_A, 'vs', tissue_type_B, sep='_')
res <- DeseqResult(dds, column, coef, tissue_type_A, tissue_type_B,
                   lfc.Threshold, rpm.Threshold,
                   norm_adj_up,
                   norm_adj_down)

dict_sig_mirna[paste(coef, "up",   sep='_')] <- list(res$up_mirna)
dict_sig_mirna[paste(coef, "down", sep='_')] <- list(res$down_mirna)
res_res <- res$res
res_dict[coef] <- res_res
plotMA(res$res, alpha=0.05)
```

```
# Plot volcano plot
VolcanoPlot(res$res, coef, res$sig,
            res$up_mirna, res$down_mirna,
            norm_adj_up, norm_adj_down,
            pCRC_adj_up, pCRC_adj_down)
```

```
ExpressionPlot(res$res, res$rpm, coef, res$sig,
               tissue_type_A, tissue_type_B,
               res$up_mirna, res$down_mirna,
               norm_adj_up, norm_adj_down,
               pCRC_adj_up, pCRC_adj_down)
```

```
signature_mirnas <- SigList(res, dds, tissue_type_A, tissue_type_B, coef,
                            norm_adj_up, norm_adj_down, 
                            pCRC_adj_up, pCRC_adj_down)
# Print list upregulated miRNA
signature_mirnas$up_mirna
```

Upregulated in tissue.type\_normal.lung\_vs\_tumor.colorect

| miRNA | LFC | lfcSE | FDR | RPM normal.lung | RPM tumor.colorect | miRBase\_ID | Family | Seed | Chr | Cell-Type Specific | Norm Background | pCRC Background |
| --- | --- | --- | --- | --- | --- | --- | --- | --- | --- | --- | --- | --- |
| Hsa-Mir-335\_5p | 1.10 | 0.19 | 1.98e-08 | 300 | 150 | hsa-mir-335 | MIR-335 | CAAGAGC | chr7 | Retinal Epithelial Cell | na | na |
| Hsa-Mir-451\_5p | 3.02 | 0.43 | 1.09e-11 | 12496 | 1565 | hsa-mir-451a | MIR-451 | AACCGUU | chr17 | Red Blood Cell | na | na |
| Hsa-Mir-144\_5p | 2.99 | 0.37 | 6.58e-15 | 451 | 57 | hsa-mir-144 | MIR-144 | GAUAUCA | chr17 | Red Blood Cell | na | na |
| Hsa-Mir-145\_5p | 1.49 | 0.36 | 9.83e-05 | 2445 | 840 | hsa-mir-145 | MIR-145 | UCCAGUU | chr5 | Mesenchymal | na | na |
| Hsa-Mir-24-P1\_3p/P2\_3p | 0.73 | 0.22 | 2.23e-03 | 1084 | 626 | hsa-mir-24-2 | MIR-24 | GGCUCAG | chr19 | Macrophage | na | na |
| Hsa-Mir-150\_5p | 1.73 | 0.39 | 1.07e-05 | 973 | 280 | hsa-mir-150 | MIR-150 | CUCCCAA | chr19 | Lymphocyte | na | na |
| Hsa-Mir-15-P1a\_5p | 0.96 | 0.17 | 2.82e-08 | 854 | 453 | hsa-mir-15a | MIR-15 | AGCAGCA | chr13 | CD14+ Monocyte | na | na |
| Hsa-Mir-486\_5p | 3.15 | 0.41 | 6.09e-14 | 22081 | 2013 | hsa-mir-486-1 | MIR-486 | CCUGUAC | chr8 | c(“Platelet”, “Red Blood Cell”) | na | na |
| Hsa-Mir-126\_5p | 3.27 | 0.23 | 4.58e-44 | 33620 | 3455 | hsa-mir-126 | MIR-126 | AUUAUUA | chr9 | c(“Endothelial Cell”, “Platelet”) | na | na |
| Hsa-Mir-146-P2\_5p | 1.39 | 0.37 | 3.72e-04 | 17435 | 6279 | hsa-mir-146b | MIR-146 | GAGAACU | chr10 | c(“Dendritic Cell”, “Lymphocyte”) | na | na |
| Hsa-Mir-342\_3p | 2.20 | 0.28 | 2.02e-14 | 681 | 145 | hsa-mir-342 | MIR-342 | CUCACAC | chr14 | c(“Dendritic Cell”, “Lymphocyte”, “Macrophage”) | na | na |
| Hsa-Mir-184\_3p | 4.90 | 0.90 | 1.36e-07 | 111 | 1 | hsa-mir-184 | MIR-184 | GGACGGA | chr15 |  | na | na |
| Hsa-Mir-34-P2a\_5p | 4.41 | 0.45 | 6.03e-22 | 116 | 5 | hsa-mir-34b | MIR-34 | GGCAGUG | chr11 |  | na | na |
| Hsa-Mir-34-P2b\_5p | 4.40 | 0.39 | 1.18e-28 | 748 | 33 | hsa-mir-34c | MIR-34 | GGCAGUG | chr11 |  | na | na |
| Hsa-Mir-30-P1a\_5p | 4.21 | 0.25 | 3.44e-60 | 60860 | 3165 | hsa-mir-30a | MIR-30 | GUAAACA | chr6 |  | na | na |
| Hsa-Mir-218-P1\_5p/P2\_5p | 3.31 | 0.30 | 3.30e-27 | 374 | 37 | hsa-mir-218-1 | MIR-218 | UGUGCUU | chr4 |  | na | na |
| Hsa-Mir-15-P2c\_5p | 2.58 | 0.26 | 1.28e-21 | 1552 | 257 | hsa-mir-195 | MIR-15 | AGCAGCA | chr17 |  | na | na |
| Hsa-Mir-338-P1\_3p | 2.57 | 0.35 | 1.28e-12 | 605 | 98 | hsa-mir-338 | MIR-338 | CCAGCAU | chr17 |  | na | na |
| Hsa-Mir-10-P2c\_5p | 2.37 | 0.37 | 4.44e-10 | 804 | 142 | hsa-mir-99a | MIR-10 | ACCCGUA | chr21 |  | na | na |
| Hsa-Mir-10-P3b\_5p | 2.28 | 0.30 | 5.11e-13 | 8561 | 1649 | hsa-mir-125a | MIR-10 | CCCUGAG | chr19 |  | na | na |
| Hsa-Mir-130-P1a\_3p | 2.26 | 0.22 | 6.52e-23 | 1941 | 421 | hsa-mir-130a | MIR-130 | AGUGCAA | chr11 |  | na | na |
| Hsa-Let-7-P1c\_5p | 2.02 | 0.31 | 4.62e-10 | 2606 | 606 | hsa-let-7c | LET-7 | GAGGUAG | chr21 |  | na | na |
| Hsa-Mir-181-P1a\_5p/P1b\_5p | 1.97 | 0.21 | 1.65e-19 | 80063 | 21233 | hsa-mir-181a-1 | MIR-181 | ACAUUCA | chr1 |  | na | na |
| Hsa-Mir-30-P1c\_5p | 1.87 | 0.18 | 4.93e-24 | 34088 | 9703 | hsa-mir-30d | MIR-30 | GUAAACA | chr8 |  | na | na |
| Hsa-Mir-101-P1\_3p/P2\_3p | 1.85 | 0.20 | 1.47e-19 | 16084 | 4638 | hsa-mir-101-1 | MIR-101 | UACAGUA | chr1 |  | na | na |
| Hsa-Mir-26-P1\_5p/P2\_5p | 1.82 | 0.15 | 1.88e-32 | 123947 | 35584 | hsa-mir-26b | MIR-26 | UCAAGUA | chr2 |  | na | na |
| Hsa-Mir-10-P3a\_5p | 1.72 | 0.35 | 2.27e-06 | 507 | 145 | hsa-mir-125b-1 | MIR-10 | CCCUGAG | chr11 |  | na | na |
| Hsa-Mir-181-P2a\_5p/P2b\_5p | 1.68 | 0.20 | 4.86e-16 | 3799 | 1251 | hsa-mir-181b-1 | MIR-181 | ACAUUCA | chr1 |  | na | na |
| Hsa-Mir-10-P3c\_5p | 1.60 | 0.34 | 3.99e-06 | 822 | 239 | hsa-mir-125b-2 | MIR-10 | CCCUGAG | chr21 |  | na | na |
| Hsa-Mir-140\_3p | 1.56 | 0.17 | 1.01e-18 | 2195 | 784 | hsa-mir-140 | MIR-140 | CCACAGG | chr16 |  | na | na |
| Hsa-Mir-15-P1c\_5p | 1.56 | 0.25 | 2.16e-09 | 405 | 143 | hsa-mir-497 | MIR-15 | AGCAGCA | chr17 |  | na | na |
| Hsa-Mir-30-P2a\_5p/P2b\_5p/P2c\_5p | 1.54 | 0.17 | 1.46e-19 | 7603 | 2793 | hsa-mir-30c-2 | MIR-30 | GUAAACA | chr6 |  | na | na |
| Hsa-Mir-92-P2b\_3p | 1.53 | 0.36 | 8.68e-05 | 137 | 47 | hsa-mir-363 | MIR-92 | AUUGCAC | chrX |  | na | na |
| Hsa-Mir-15-P2a\_5p/P2b\_5p | 1.48 | 0.19 | 3.24e-14 | 12836 | 4522 | hsa-mir-16-1 | MIR-15 | AGCAGCA | chr13 |  | na | na |
| Hsa-Mir-26-P3\_5p | 1.37 | 0.22 | 1.19e-09 | 9029 | 3467 | hsa-mir-26a-1 | MIR-26 | UCAAGUA | chr3 |  | na | na |
| Hsa-Mir-10-P2a\_5p | 1.32 | 0.36 | 6.59e-04 | 4056 | 1724 | hsa-mir-100 | MIR-10 | ACCCGUA | chr11 |  | na | na |
| Hsa-Let-7-P2b1\_5p | 1.23 | 0.14 | 9.88e-19 | 6253 | 2784 | hsa-let-7f-1 | LET-7 | GAGGUAG | chr9 |  | na | na |
| Hsa-Mir-331\_3p | 1.11 | 0.29 | 2.07e-04 | 128 | 60 | hsa-mir-331 | MIR-331 | CCCCUGG | chr12 |  | na | na |
| Hsa-Mir-10-P2b\_5p | 1.06 | 0.42 | 2.28e-02 | 5968 | 2594 | hsa-mir-99b | MIR-10 | ACCCGUA | chr19 |  | na | na |
| Hsa-Mir-148-P2\_3p | 0.98 | 0.34 | 6.10e-03 | 270 | 137 | hsa-mir-152 | MIR-148 | CAGUGCA | chr17 |  | na | na |
| Hsa-Mir-29-P1b\_3p | 0.96 | 0.23 | 6.29e-05 | 607 | 331 | hsa-mir-29c | MIR-29 | AGCACCA | chr1 |  | na | na |
| Hsa-Mir-652\_3p | 0.88 | 0.18 | 1.77e-06 | 187 | 106 | hsa-mir-652 | MIR-652 | AUGGCGC | chrX |  | na | na |
| Hsa-Mir-744\_5p | 0.88 | 0.24 | 5.22e-04 | 109 | 61 | hsa-mir-744 | MIR-744 | GCGGGGC | chr17 |  | na | na |
| Hsa-Let-7-P2b2\_5p | 0.84 | 0.17 | 3.85e-06 | 6575 | 3787 | hsa-let-7b | LET-7 | GAGGUAG | chr22 |  | na | na |
| Hsa-Mir-27-P1\_3p/P2\_3p | 0.80 | 0.16 | 4.13e-06 | 50854 | 29444 | hsa-mir-27a | MIR-27 | UCACAGU | chr19 |  | na | na |
| Hsa-Mir-374-P2\_5p | 0.74 | 0.22 | 1.39e-03 | 148 | 95 | hsa-mir-374b | MIR-374 | UAUAAUA | chrX |  | na | na |
| Hsa-Mir-23-P1\_3p/P2\_3p | 0.68 | 0.17 | 1.35e-04 | 5786 | 3766 | hsa-mir-23a | MIR-23 | UCACAUU | chr19 |  | na | na |
| Hsa-Mir-28-P2\_3p | 0.65 | 0.19 | 1.33e-03 | 6007 | 3830 | hsa-mir-151a | MIR-28 | CGAGGAG | chr8 |  | na | na |

```
# Number of upregulated miRNA
signature_mirnas$number_upregulated
```

```
## [1] 48
```

```
# Print list downregulated miRNA
signature_mirnas$down_mirna
```

Downregulated in tissue.type\_normal.lung\_vs\_tumor.colorect

| miRNA | LFC | lfcSE | FDR | RPM normal.lung | RPM tumor.colorect | miRBase\_ID | Family | Seed | Chr | Cell-Type Specific | Norm Background | pCRC Background |
| --- | --- | --- | --- | --- | --- | --- | --- | --- | --- | --- | --- | --- |
| Hsa-Mir-128-P1\_3p/P2\_3p | -0.86 | 0.18 | 6.72e-06 | 116 | 233 | hsa-mir-128-1 | MIR-128 | CACAGUG | chr2 | Neural | na | na |
| Hsa-Mir-192-P1\_5p/P2\_5p | -6.50 | 0.36 | 4.31e-69 | 142 | 13717 | hsa-mir-192 | MIR-192 | UGACCUA | chr11 | Epithelial Cell | na | na |
| Hsa-Mir-8-P2a\_3p | -3.88 | 0.26 | 9.46e-48 | 618 | 10330 | hsa-mir-200b | MIR-8 | AAUACUG | chr1 | Epithelial Cell | na | na |
| Hsa-Mir-8-P2b\_3p | -1.75 | 0.26 | 7.14e-11 | 766 | 2727 | hsa-mir-200c | MIR-8 | AAUACUG | chr12 | Epithelial Cell | na | na |
| Hsa-Mir-17-P1a\_5p/P1b\_5p | -2.17 | 0.21 | 9.51e-23 | 184 | 900 | hsa-mir-17 | MIR-17 | AAAGUGC | chr13 | CD14+ Monocyte | na | na |
| Hsa-Mir-7-P1\_5p/P2\_5p/P3\_5p | -5.51 | 0.42 | 5.13e-38 | 3 | 199 | hsa-mir-7-1 | MIR-7 | GGAAGAC | chr9 | c(“Islet Cell”, “Neural”) | na | na |
| Hsa-Mir-375\_3p | -1.22 | 0.43 | 1.11e-02 | 2346 | 5263 | hsa-mir-375 | MIR-375 | UUGUUCG | chr2 | c(“Epithelial Cell”, “Islet Cell”, “Neural”) | na | na |
| Hsa-Mir-196-P1\_5p/P2\_5p | -6.60 | 0.38 | 7.65e-66 | 3 | 319 | hsa-mir-196a-1 | MIR-196 | AGGUAGU | chr17 |  | na | na |
| Hsa-Mir-194-P1\_5p/P2\_5p | -6.59 | 0.31 | 7.31e-100 | 58 | 6167 | hsa-mir-194-2 | MIR-194 | GUAACAG | chr11 |  | na | na |
| Hsa-Mir-196-P3\_5p | -6.49 | 0.43 | 9.67e-50 | 7 | 828 | hsa-mir-196b | MIR-196 | AGGUAGU | chr7 |  | na | na |
| Hsa-Mir-192-P1\_5p | -6.17 | 0.31 | 1.45e-85 | 1684 | 132000 | hsa-mir-192 | MIR-192 | UGACCUA | chr11 |  | na | na |
| Hsa-Mir-577\_5p | -5.61 | 0.37 | 3.74e-49 | 3 | 142 | hsa-mir-577 | MIR-577 | UAGAUAA | chr4 |  | na | na |
| Hsa-Mir-31\_5p | -4.10 | 0.59 | 3.10e-12 | 24 | 627 | hsa-mir-31 | MIR-31 | GGCAAGA | chr9 |  | na | na |
| Hsa-Mir-8-P3a\_3p | -2.93 | 0.27 | 4.00e-27 | 262 | 2303 | hsa-mir-429 | MIR-8 | AAUACUG | chr1 |  | na | na |
| Hsa-Mir-17-P3a\_5p | -2.82 | 0.26 | 3.30e-27 | 194 | 1513 | hsa-mir-20a | MIR-17 | AAAGUGC | chr13 |  | na | na |
| Hsa-Mir-135-P3\_5p | -2.73 | 0.37 | 6.36e-13 | 20 | 155 | hsa-mir-135b | MIR-135 | AUGGCUU | chr1 |  | na | na |
| Hsa-Mir-127\_3p | -2.70 | 0.36 | 7.71e-13 | 327 | 2263 | hsa-mir-127 | MIR-127 | CGGAUCC | chr14 |  | na | na |
| Hsa-Mir-190-P1\_5p | -2.56 | 0.29 | 1.90e-17 | 22 | 140 | hsa-mir-190a | MIR-190 | GAUAUGU | chr15 |  | na | na |
| Hsa-Mir-224\_5p | -2.55 | 0.30 | 2.30e-16 | 59 | 395 | hsa-mir-224 | MIR-224 | AAGUCAC | chrX |  | na | na |
| Hsa-Mir-154-P36\_3p | -2.22 | 0.23 | 2.04e-21 | 39 | 195 | hsa-mir-409 | MIR-154 | AAUGUUG | chr14 |  | na | na |
| Hsa-Mir-8-P1a\_3p | -2.20 | 0.26 | 7.55e-16 | 353 | 1772 | hsa-mir-200a | MIR-8 | AACACUG | chr1 |  | na | na |
| Hsa-Mir-21\_5p | -2.20 | 0.19 | 1.45e-28 | 21349 | 105735 | hsa-mir-21 | MIR-21 | AGCUUAU | chr17 |  | na | na |
| Hsa-Mir-15-P1d\_5p | -2.15 | 0.34 | 9.48e-10 | 46 | 235 | hsa-mir-424 | MIR-15 | AGCAGCA | chrX |  | na | na |
| Hsa-Mir-17-P2a\_5p | -2.09 | 0.31 | 5.05e-11 | 33 | 168 | hsa-mir-18a | MIR-17 | AAGGUGC | chr13 |  | na | na |
| Hsa-Mir-130-P1b\_3p | -1.94 | 0.22 | 2.67e-18 | 28 | 116 | hsa-mir-130b | MIR-130 | AGUGCAA | chr22 |  | na | na |
| Hsa-Mir-96-P3\_5p | -1.94 | 0.37 | 3.95e-07 | 291 | 1063 | hsa-mir-183 | MIR-96 | AUGGCAC | chr7 |  | na | na |
| Hsa-Mir-96-P2\_5p | -1.91 | 0.27 | 1.11e-11 | 2685 | 10417 | hsa-mir-182 | MIR-96 | UUGGCAA | chr7 |  | na | na |
| Hsa-Mir-154-P23\_3p | -1.74 | 0.22 | 1.04e-13 | 41 | 148 | hsa-mir-654 | MIR-154 | AUGUCUG | chr14 |  | na | na |
| Hsa-Mir-19-P1\_3p | -1.65 | 0.26 | 1.09e-09 | 108 | 379 | hsa-mir-19a | MIR-19 | GUGCAAA | chr13 |  | na | na |
| Hsa-Mir-425\_5p | -1.65 | 0.21 | 2.02e-14 | 213 | 700 | hsa-mir-425 | MIR-425 | AUGACAC | chr3 |  | na | na |
| Hsa-Mir-154-P12\_3p | -1.63 | 0.24 | 1.28e-10 | 33 | 115 | hsa-mir-410 | MIR-154 | AUAUAAC | chr14 |  | na | na |
| Hsa-Mir-10-P1b\_5p | -1.55 | 0.41 | 4.05e-04 | 14125 | 39286 | hsa-mir-10b | MIR-10 | ACCCUGU | chr2 |  | na | na |
| Hsa-Mir-378\_3p | -1.45 | 0.22 | 9.02e-11 | 2189 | 6426 | hsa-mir-378a | MIR-378 | CUGGACU | chr5 |  | na | na |
| Hsa-Mir-584\_5p | -1.38 | 0.34 | 1.18e-04 | 39 | 110 | hsa-mir-584 | MIR-584 | UAUGGUU | chr5 |  | na | na |
| Hsa-Mir-92-P1a\_3p/P1b\_3p | -1.27 | 0.25 | 8.94e-07 | 15850 | 39320 | hsa-mir-92a-1 | MIR-92 | AUUGCAC | chr13 |  | na | na |
| Hsa-Mir-17-P3c\_5p | -1.21 | 0.18 | 2.57e-11 | 143 | 370 | hsa-mir-106b | MIR-17 | AAAGUGC | chr7 |  | na | na |
| Hsa-Mir-17-P1c\_5p | -1.06 | 0.15 | 2.20e-11 | 1024 | 2287 | hsa-mir-93 | MIR-17 | AAAGUGC | chr7 |  | na | na |
| Hsa-Mir-203\_3p | -1.03 | 0.32 | 2.53e-03 | 1068 | 2307 | hsa-mir-203a | MIR-203 | UGAAAUG | chr14 |  | na | na |
| Hsa-Mir-210\_3p | -1.01 | 0.31 | 4.02e-03 | 138 | 291 | hsa-mir-210 | MIR-210 | UGUGCGU | chr11 |  | na | na |
| Hsa-Mir-95-P2\_3p | -0.99 | 0.22 | 1.35e-05 | 65 | 142 | hsa-mir-421 | MIR-95 | UCAACAG | chrX |  | na | na |
| Hsa-Mir-154-P13\_5p | -0.93 | 0.24 | 3.43e-04 | 161 | 328 | hsa-mir-411 | MIR-154 | AGUAGAC | chr14 |  | na | na |
| Hsa-Mir-193-P1b\_3p | -0.92 | 0.26 | 8.53e-04 | 72 | 145 | hsa-mir-193b | MIR-193 | ACUGGCC | chr16 |  | na | na |
| Hsa-Mir-19-P2a\_3p/P2b\_3p | -0.91 | 0.24 | 5.22e-04 | 645 | 1297 | hsa-mir-19b-1 | MIR-19 | GUGCAAA | chr13 |  | na | na |
| Hsa-Mir-148-P1\_3p | -0.86 | 0.27 | 3.44e-03 | 18874 | 34704 | hsa-mir-148a | MIR-148 | CAGUGCA | chr7 |  | na | na |
| Hsa-Mir-146-P1\_5p | -0.85 | 0.39 | 4.76e-02 | 786 | 1453 | hsa-mir-146a | MIR-146 | GAGAACU | chr5 |  | na | na |
| Hsa-Mir-136\_3p | -0.81 | 0.24 | 1.54e-03 | 88 | 173 | hsa-mir-136 | MIR-136 | AUCAUCG | chr14 |  | na | na |
| Hsa-Mir-10-P1a\_5p | -0.79 | 0.34 | 4.36e-02 | 61989 | 97123 | hsa-mir-10a | MIR-10 | ACCCUGU | chr17 |  | na | na |
| Hsa-Mir-154-P9\_3p | -0.78 | 0.23 | 1.67e-03 | 135 | 253 | hsa-mir-381 | MIR-154 | AUACAAG | chr14 |  | na | na |
| Hsa-Mir-214\_3p | -0.74 | 0.26 | 8.05e-03 | 77 | 137 | hsa-mir-214 | MIR-214 | CAGCAGG | chr1 |  | na | na |
| Hsa-Mir-1307\_3p | -0.74 | 0.21 | 1.32e-03 | 80 | 139 | hsa-mir-1307 | MIR-1307 | CGACCGG | chr10 |  | na | na |
| Hsa-Let-7-P2b3\_5p | -0.72 | 0.21 | 1.18e-03 | 1289 | 2149 | hsa-mir-98 | LET-7 | GAGGUAG | chrX |  | na | na |
| Hsa-Mir-199-P1\_3p/P2\_3p/P3\_3p | -0.64 | 0.22 | 5.97e-03 | 3382 | 5772 | hsa-mir-199b | MIR-199 | CAGUAGU | chr9 |  | na | na |

```
# Number of downregulated miRNA
signature_mirnas$number_downregulated
```

```
## [1] 52
```

```
SubtractLFC <- function(x, y){
  z = x
  if ( is.na(x) | is.na(y) ){ return( z ) }
  else if (sign(x) == sign(y)){ z = x - y }
  if (sign(z) != sign(x)) { z = 0 }
  return( z )
}

SubtractAdjP <- function(x , y, xP, yP){
  z = xP
  if ( is.na(xP) | is.na(yP) ){ return( z ) }
  if ( sign(x) == sign(y) ){ 
    z = (xP + ( 1 - yP )) }
  if (z > 1) {z = 1}
  return(z)
}
```

#### pCRC vs mLi

```
#pCRC versus liver metastasis, control also with pCRC versus normal liver

column='tissue.type'
tissue_type_A <- 'metastasis.liver'
tissue_type_B <- 'tumor.colorect'
norm_adj_up       = dict_sig_mirna$tissue.type_normal.liver_vs_normal.colorect_up
norm_adj_down     = dict_sig_mirna$tissue.type_normal.liver_vs_normal.colorect_down
pCRC_adj_up   = dict_sig_mirna$tissue.type_normal.liver_vs_tumor.colorect_up
pCRC_adj_down = dict_sig_mirna$tissue.type_normal.liver_vs_tumor.colorect_down
palette <- 'jco'

coef <- paste(column, tissue_type_A, 'vs', tissue_type_B, sep='_')
res <- DeseqResult(dds, column, coef, tissue_type_A, tissue_type_B,
                   lfc.Threshold, rpm.Threshold,
                   norm_adj_up,
                   norm_adj_down,
                   pCRC_adj_up,
                   pCRC_adj_down)

dict_sig_mirna[paste(coef, "up",   sep='_')] <- list(res$up_mirna)
dict_sig_mirna[paste(coef, "down", sep='_')] <- list(res$down_mirna)
res_res <- res$res
res_dict[coef] <- res_res
plotMA(res$res, alpha=0.05)
```

```
# Plot volcano plot
VolcanoPlot(res$res, coef, res$sig,
            res$up_mirna, res$down_mirna,
            norm_adj_up, norm_adj_down,
            pCRC_adj_up, pCRC_adj_down)
```

```
ExpressionPlot(res$res, res$rpm, coef, res$sig,
               tissue_type_A, tissue_type_B,
               res$up_mirna, res$down_mirna,
               norm_adj_up, norm_adj_down,
               pCRC_adj_up, pCRC_adj_down)
```

```
signature_mirnas <- SigList(res, dds, tissue_type_A, tissue_type_B, coef,
                            norm_adj_up, norm_adj_down, 
                            pCRC_adj_up, pCRC_adj_down)
# Print list upregulated miRNA
signature_mirnas$up_mirna
```

Upregulated in tissue.type\_metastasis.liver\_vs\_tumor.colorect

| miRNA | LFC | lfcSE | FDR | RPM metastasis.liver | RPM tumor.colorect | miRBase\_ID | Family | Seed | Chr | Cell-Type Specific | Norm Background | pCRC Background |
| --- | --- | --- | --- | --- | --- | --- | --- | --- | --- | --- | --- | --- |
| Hsa-Mir-210\_3p | 1.26 | 0.18 | 4.73e-11 | 685 | 291 | hsa-mir-210 | MIR-210 | UGUGCGU | chr11 |  |  |  |
| Hsa-Mir-592\_5p | 0.98 | 0.27 | 4.88e-03 | 137 | 75 | hsa-mir-592 | MIR-592 | UGUGUCA | chr7 |  |  |  |
| Hsa-Mir-10-P1a\_5p | 0.86 | 0.19 | 8.04e-05 | 164925 | 97123 | hsa-mir-10a | MIR-10 | ACCCUGU | chr17 |  |  |  |
| Hsa-Mir-1307\_5p | 0.85 | 0.17 | 2.99e-06 | 887 | 492 | hsa-mir-1307 | MIR-1307 | CGACCGG | chr10 |  |  |  |
| Hsa-Mir-1247\_5p | 0.85 | 0.28 | 2.58e-02 | 127 | 69 | hsa-mir-1247 | MIR-1247 | CCCGUCC | chr14 |  |  |  |
| Hsa-Mir-191\_5p | 0.74 | 0.15 | 3.67e-06 | 23253 | 13630 | hsa-mir-191 | MIR-191 | AACGGAA | chr3 |  |  |  |
| Hsa-Mir-425\_5p | 0.73 | 0.12 | 2.90e-08 | 1117 | 700 | hsa-mir-425 | MIR-425 | AUGACAC | chr3 |  |  |  |
| Hsa-Mir-8-P1b\_3p | 0.64 | 0.19 | 4.07e-03 | 11957 | 7518 | hsa-mir-141 | MIR-8 | AACACUG | chr12 |  |  |  |
| Hsa-Mir-150\_5p | 1.08 | 0.22 | 2.99e-06 | 721 | 280 | hsa-mir-150 | MIR-150 | CUCCCAA | chr19 | Lymphocyte |  | yes |
| Hsa-Mir-342\_3p | 0.93 | 0.16 | 5.32e-08 | 320 | 145 | hsa-mir-342 | MIR-342 | CUCACAC | chr14 | c(“Dendritic Cell”, “Lymphocyte”, “Macrophage”) |  | yes |
| Hsa-Mir-331\_3p | 0.75 | 0.16 | 3.16e-05 | 107 | 60 | hsa-mir-331 | MIR-331 | CCCCUGG | chr12 |  |  | yes |
| Hsa-Mir-15-P2a\_5p/P2b\_5p | 0.68 | 0.11 | 1.34e-08 | 7588 | 4522 | hsa-mir-16-1 | MIR-15 | AGCAGCA | chr13 |  |  | yes |
| Hsa-Mir-204-P1\_5p | 1.04 | 0.32 | 1.21e-06 | 260 | 62 | hsa-mir-204 | MIR-204 | UCCCUUU | chr9 | Retinal Epithelial Cell | yes | yes |
| Hsa-Mir-335\_5p | 0.78 | 0.11 | 5.33e-11 | 261 | 150 | hsa-mir-335 | MIR-335 | CAAGAGC | chr7 | Retinal Epithelial Cell | yes | yes |
| Hsa-Mir-122\_5p | 5.14 | 0.43 | 2.78e-29 | 3668 | 9 | hsa-mir-122 | MIR-122 | GGAGUGU | chr18 | Hepatocyte | yes | yes |
| Hsa-Mir-10-P3c\_5p | 0.71 | 0.19 | 7.30e-04 | 480 | 239 | hsa-mir-125b-2 | MIR-10 | CCCUGAG | chr21 |  | yes | yes |

```
# Number of upregulated miRNA
signature_mirnas$number_upregulated
```

```
## [1] 16
```

```
# Print list downregulated miRNA
signature_mirnas$down_mirna
```

Downregulated in tissue.type\_metastasis.liver\_vs\_tumor.colorect

| miRNA | LFC | lfcSE | FDR | RPM metastasis.liver | RPM tumor.colorect | miRBase\_ID | Family | Seed | Chr | Cell-Type Specific | Norm Background | pCRC Background |
| --- | --- | --- | --- | --- | --- | --- | --- | --- | --- | --- | --- | --- |
| Hsa-Mir-486\_5p | -0.66 | 0.23 | 3.47e-02 | 1489 | 2013 | hsa-mir-486-1 | MIR-486 | CCUGUAC | chr8 | c(“Platelet”, “Red Blood Cell”) |  |  |
| Hsa-Mir-31\_5p | -2.34 | 0.31 | 4.05e-13 | 88 | 627 | hsa-mir-31 | MIR-31 | GGCAAGA | chr9 |  |  | yes |
| Hsa-Let-7-P2c2\_5p | -0.90 | 0.11 | 1.69e-14 | 2840 | 5432 | hsa-let-7i | LET-7 | GAGGUAG | chr12 |  |  | yes |
| Hsa-Mir-143\_3p | -1.02 | 0.17 | 4.70e-08 | 72945 | 148431 | hsa-mir-143 | MIR-143 | GAGAUGA | chr5 | Mesenchymal | yes | yes |
| Hsa-Mir-133-P1\_3p/P2\_3p/P3\_3p | -1.56 | 0.24 | 3.43e-10 | 38 | 116 | hsa-mir-133a-2 | MIR-133 | UUGGUCC | chr20 | c(“Skeletal Myocyte”, “Stem Cell”) | yes | yes |
| Hsa-Mir-10-P1b\_5p | -1.62 | 0.23 | 1.76e-10 | 11785 | 39286 | hsa-mir-10b | MIR-10 | ACCCUGU | chr2 |  | yes | yes |
| Hsa-Mir-92-P1c\_3p | -0.68 | 0.17 | 2.83e-04 | 879 | 1398 | hsa-mir-92b | MIR-92 | AUUGCAC | chr1 |  | yes | yes |
| Hsa-Mir-146-P1\_5p | -0.63 | 0.22 | 1.91e-02 | 909 | 1453 | hsa-mir-146a | MIR-146 | GAGAACU | chr5 |  | yes | yes |

```
# Number of downregulated miRNA
signature_mirnas$number_downregulated
```

```
## [1] 8
```

```
res_tibble <- res$res
res_tibble$miRNA <- rownames(res_tibble)
res_tibble <- as_tibble(res_tibble)

metslfc <- res_dict$tissue.type_metastasis.liver_vs_tumor.colorect$log2FoldChange
normlfc <- res_dict$tissue.type_normal.liver_vs_normal.colorect$log2FoldChange

res_tibble$LFC_adj_background <- mapply(SubtractLFC, metslfc, normlfc)

metsP <- res_dict$tissue.type_metastasis.liver_vs_tumor.colorect$padj
normP <- res_dict$tissue.type_normal.liver_vs_normal.colorect$padj

res_tibble$padj_subt_normal <- mapply( SubtractAdjP, metslfc, normlfc, metsP, normP )

res_tibble %>% select(miRNA, log2FoldChange, lfcSE, LFC_adj_background, padj_subt_normal, baseMean, stat, pvalue, padj) %>% write_csv(path = '/Users/eirikhoy/Dropbox/projects/comet_analysis/data/Deseq_result_clm_vs_pcrc.csv')
```

#### pCRC vs mLu

```
#pCRC versus lung metastasis, control also with pCRC versus normal liver

column='tissue.type'
tissue_type_A <- 'metastasis.lung'
tissue_type_B <- 'tumor.colorect'
norm_adj_up       = dict_sig_mirna$tissue.type_normal.lung_vs_normal.colorect_up
norm_adj_down     = dict_sig_mirna$tissue.type_normal.lung_vs_normal.colorect_down
pCRC_adj_up   = dict_sig_mirna$tissue.type_normal.lung_vs_tumor.colorect_up
pCRC_adj_down = dict_sig_mirna$tissue.type_normal.lung_vs_tumor.colorect_down

coef <- paste(column, tissue_type_A, 'vs', tissue_type_B, sep='_')
res <- DeseqResult(dds, column, coef, tissue_type_A, tissue_type_B,
                   lfc.Threshold, rpm.Threshold,
                   norm_adj_up,
                   norm_adj_down,
                   pCRC_adj_up,
                   pCRC_adj_down)
dict_sig_mirna[paste(coef, "up",   sep='_')] <- list(res$up_mirna)
dict_sig_mirna[paste(coef, "down", sep='_')] <- list(res$down_mirna)
res_res <- res$res
res_dict[coef] <- res_res
plotMA(res$res, alpha=0.05)
```

```
# Plot volcano plot
VolcanoPlot(res$res, coef, res$sig,
            res$up_mirna, res$down_mirna,
            norm_adj_up, norm_adj_down,
            pCRC_adj_up, pCRC_adj_down)
```

```
ExpressionPlot(res$res, res$rpm, coef, res$sig,
               tissue_type_A, tissue_type_B,
               res$up_mirna, res$down_mirna,
               norm_adj_up, norm_adj_down,
               pCRC_adj_up, pCRC_adj_down)
```

```
signature_mirnas <- SigList(res, dds, tissue_type_A, tissue_type_B, coef,
                            norm_adj_up, norm_adj_down, 
                            pCRC_adj_up, pCRC_adj_down)
# Print list upregulated miRNA
signature_mirnas$up_mirna
```

Upregulated in tissue.type\_metastasis.lung\_vs\_tumor.colorect

| miRNA | LFC | lfcSE | FDR | RPM metastasis.lung | RPM tumor.colorect | miRBase\_ID | Family | Seed | Chr | Cell-Type Specific | Norm Background | pCRC Background |
| --- | --- | --- | --- | --- | --- | --- | --- | --- | --- | --- | --- | --- |
| Hsa-Mir-155\_5p | 0.76 | 0.18 | 7.24e-05 | 733 | 387 | hsa-mir-155 | MIR-155 | UAAUGCU | chr21 | c(“Lymphocyte”, “Macrophage”) |  |  |
| Hsa-Mir-210\_3p | 1.15 | 0.19 | 1.55e-08 | 693 | 291 | hsa-mir-210 | MIR-210 | UGUGCGU | chr11 |  |  |  |
| Hsa-Mir-142\_5p | 0.84 | 0.18 | 2.01e-05 | 7117 | 3453 | hsa-mir-142 | MIR-142 | AUAAAGU | chr17 |  |  |  |
| Hsa-Mir-19-P2a\_3p/P2b\_3p | 0.80 | 0.15 | 9.38e-07 | 2345 | 1297 | hsa-mir-19b-1 | MIR-19 | GUGCAAA | chr13 |  |  |  |
| Hsa-Mir-8-P1b\_3p | 0.75 | 0.21 | 1.12e-03 | 15111 | 7518 | hsa-mir-141 | MIR-8 | AACACUG | chr12 |  |  |  |
| Hsa-Mir-191\_5p | 0.74 | 0.16 | 9.81e-06 | 26989 | 13630 | hsa-mir-191 | MIR-191 | AACGGAA | chr3 |  |  |  |
| Hsa-Mir-374-P1\_5p | 0.72 | 0.12 | 9.07e-09 | 217 | 115 | hsa-mir-374a | MIR-374 | UAUAAUA | chrX |  |  |  |
| Hsa-Mir-19-P1\_3p | 0.65 | 0.16 | 3.42e-04 | 597 | 379 | hsa-mir-19a | MIR-19 | GUGCAAA | chr13 |  |  |  |
| Hsa-Mir-145\_5p | 0.67 | 0.22 | 6.74e-03 | 1629 | 840 | hsa-mir-145 | MIR-145 | UCCAGUU | chr5 | Mesenchymal |  | yes |
| Hsa-Mir-150\_5p | 1.14 | 0.24 | 2.51e-06 | 738 | 280 | hsa-mir-150 | MIR-150 | CUCCCAA | chr19 | Lymphocyte |  | yes |
| Hsa-Mir-10-P3c\_5p | 0.85 | 0.21 | 9.78e-05 | 574 | 239 | hsa-mir-125b-2 | MIR-10 | CCCUGAG | chr21 |  |  | yes |
| Hsa-Mir-29-P1b\_3p | 0.71 | 0.14 | 3.39e-06 | 597 | 331 | hsa-mir-29c | MIR-29 | AGCACCA | chr1 |  |  | yes |
| Hsa-Mir-15-P2a\_5p/P2b\_5p | 0.71 | 0.12 | 1.51e-08 | 8258 | 4522 | hsa-mir-16-1 | MIR-15 | AGCAGCA | chr13 |  |  | yes |
| Hsa-Mir-26-P3\_5p | 0.65 | 0.14 | 6.79e-06 | 6205 | 3467 | hsa-mir-26a-1 | MIR-26 | UCAAGUA | chr3 |  |  | yes |
| Hsa-Mir-331\_3p | 0.62 | 0.18 | 1.05e-03 | 105 | 60 | hsa-mir-331 | MIR-331 | CCCCUGG | chr12 |  |  | yes |
| Hsa-Mir-335\_5p | 0.88 | 0.12 | 1.45e-12 | 295 | 150 | hsa-mir-335 | MIR-335 | CAAGAGC | chr7 | Retinal Epithelial Cell | yes | yes |
| Hsa-Mir-451\_5p | 0.87 | 0.26 | 1.88e-03 | 3152 | 1565 | hsa-mir-451a | MIR-451 | AACCGUU | chr17 | Red Blood Cell | yes | yes |
| Hsa-Mir-24-P1\_3p/P2\_3p | 0.74 | 0.14 | 7.89e-07 | 1289 | 626 | hsa-mir-24-2 | MIR-24 | GGCUCAG | chr19 | Macrophage | yes | yes |
| Hsa-Mir-126\_5p | 1.67 | 0.15 | 6.56e-29 | 13328 | 3455 | hsa-mir-126 | MIR-126 | AUUAUUA | chr9 | c(“Endothelial Cell”, “Platelet”) | yes | yes |
| Hsa-Mir-146-P2\_5p | 0.94 | 0.23 | 1.47e-04 | 15403 | 6279 | hsa-mir-146b | MIR-146 | GAGAACU | chr10 | c(“Dendritic Cell”, “Lymphocyte”) | yes | yes |
| Hsa-Mir-342\_3p | 2.25 | 0.18 | 6.24e-36 | 872 | 145 | hsa-mir-342 | MIR-342 | CUCACAC | chr14 | c(“Dendritic Cell”, “Lymphocyte”, “Macrophage”) | yes | yes |
| Hsa-Mir-34-P2b\_5p | 4.24 | 0.24 | 2.76e-67 | 989 | 33 | hsa-mir-34c | MIR-34 | GGCAGUG | chr11 |  | yes | yes |
| Hsa-Mir-34-P2a\_5p | 3.97 | 0.27 | 8.59e-45 | 128 | 5 | hsa-mir-34b | MIR-34 | GGCAGUG | chr11 |  | yes | yes |
| Hsa-Mir-30-P1a\_5p | 1.74 | 0.16 | 1.56e-26 | 13224 | 3165 | hsa-mir-30a | MIR-30 | GUAAACA | chr6 |  | yes | yes |
| Hsa-Mir-15-P2c\_5p | 1.64 | 0.17 | 6.99e-22 | 975 | 257 | hsa-mir-195 | MIR-15 | AGCAGCA | chr17 |  | yes | yes |
| Hsa-Mir-218-P1\_5p/P2\_5p | 1.57 | 0.19 | 3.94e-16 | 135 | 37 | hsa-mir-218-1 | MIR-218 | UGUGCUU | chr4 |  | yes | yes |
| Hsa-Mir-374-P2\_5p | 1.14 | 0.14 | 1.78e-15 | 226 | 95 | hsa-mir-374b | MIR-374 | UAUAAUA | chrX |  | yes | yes |
| Hsa-Mir-338-P1\_3p | 1.10 | 0.22 | 1.51e-06 | 236 | 98 | hsa-mir-338 | MIR-338 | CCAGCAU | chr17 |  | yes | yes |
| Hsa-Mir-26-P1\_5p/P2\_5p | 1.07 | 0.10 | 7.47e-28 | 84008 | 35584 | hsa-mir-26b | MIR-26 | UCAAGUA | chr2 |  | yes | yes |
| Hsa-Mir-10-P2c\_5p | 0.63 | 0.23 | 8.68e-03 | 289 | 142 | hsa-mir-99a | MIR-10 | ACCCGUA | chr21 |  | yes | yes |
| Hsa-Mir-130-P1a\_3p | 0.62 | 0.14 | 4.12e-05 | 729 | 421 | hsa-mir-130a | MIR-130 | AGUGCAA | chr11 |  | yes | yes |

```
# Number of upregulated miRNA
signature_mirnas$number_upregulated
```

```
## [1] 31
```

```
# Print list downregulated miRNA
signature_mirnas$down_mirna
```

Downregulated in tissue.type\_metastasis.lung\_vs\_tumor.colorect

| miRNA | LFC | lfcSE | FDR | RPM metastasis.lung | RPM tumor.colorect | miRBase\_ID | Family | Seed | Chr | Cell-Type Specific | Norm Background | pCRC Background |
| --- | --- | --- | --- | --- | --- | --- | --- | --- | --- | --- | --- | --- |
| Hsa-Mir-423\_5p | -1.17 | 0.17 | 1.35e-10 | 202 | 433 | hsa-mir-423 | MIR-423 | GAGGGGC | chr17 |  |  |  |
| Hsa-Let-7-P1b\_5p | -0.86 | 0.16 | 2.55e-07 | 650 | 1057 | hsa-let-7e | LET-7 | GAGGUAG | chr19 |  |  |  |
| Hsa-Mir-197\_3p | -0.74 | 0.13 | 1.76e-07 | 113 | 181 | hsa-mir-197 | MIR-197 | UCACCAC | chr1 |  |  |  |
| Hsa-Mir-92-P1c\_3p | -0.72 | 0.18 | 2.22e-04 | 981 | 1398 | hsa-mir-92b | MIR-92 | AUUGCAC | chr1 |  |  |  |
| Hsa-Mir-362-P2\_3p/P4\_3p | -0.65 | 0.14 | 2.36e-05 | 302 | 441 | hsa-mir-500a | MIR-362 | UGCACCU | chrX |  |  |  |
| Hsa-Mir-221-P2\_3p | -0.60 | 0.14 | 8.50e-05 | 1101 | 1633 | hsa-mir-222 | MIR-221 | GCUACAU | chrX |  |  |  |
| Hsa-Mir-7-P1\_5p/P2\_5p/P3\_5p | -0.80 | 0.25 | 1.88e-03 | 106 | 199 | hsa-mir-7-1 | MIR-7 | GGAAGAC | chr9 | c(“Islet Cell”, “Neural”) |  | yes |
| Hsa-Mir-154-P36\_3p | -1.40 | 0.14 | 3.20e-21 | 80 | 195 | hsa-mir-409 | MIR-154 | AAUGUUG | chr14 |  |  | yes |
| Hsa-Mir-154-P12\_3p | -1.38 | 0.15 | 7.64e-18 | 46 | 115 | hsa-mir-410 | MIR-154 | AUAUAAC | chr14 |  |  | yes |
| Hsa-Mir-31\_5p | -1.01 | 0.34 | 1.99e-03 | 242 | 627 | hsa-mir-31 | MIR-31 | GGCAAGA | chr9 |  |  | yes |
| Hsa-Mir-96-P2\_5p | -0.78 | 0.17 | 1.58e-05 | 6399 | 10417 | hsa-mir-182 | MIR-96 | UUGGCAA | chr7 |  |  | yes |
| Hsa-Mir-199-P1\_3p/P2\_3p/P3\_3p | -0.65 | 0.14 | 1.06e-05 | 3904 | 5772 | hsa-mir-199b | MIR-199 | CAGUAGU | chr9 |  |  | yes |
| Hsa-Mir-15-P1d\_5p | -0.63 | 0.21 | 5.51e-03 | 158 | 235 | hsa-mir-424 | MIR-15 | AGCAGCA | chrX |  |  | yes |
| Hsa-Mir-92-P1a\_3p/P1b\_3p | -0.63 | 0.15 | 1.76e-04 | 26467 | 39320 | hsa-mir-92a-1 | MIR-92 | AUUGCAC | chr13 |  |  | yes |
| Hsa-Mir-143\_3p | -1.34 | 0.19 | 1.00e-11 | 67942 | 148431 | hsa-mir-143 | MIR-143 | GAGAUGA | chr5 | Mesenchymal | yes |  |
| Hsa-Mir-133-P1\_3p/P2\_3p/P3\_3p | -1.73 | 0.26 | 4.97e-11 | 37 | 116 | hsa-mir-133a-2 | MIR-133 | UUGGUCC | chr20 | c(“Skeletal Myocyte”, “Stem Cell”) | yes |  |
| Hsa-Mir-362-P3\_3p | -0.89 | 0.25 | 1.47e-03 | 61 | 108 | hsa-mir-501 | MIR-362 | AUGCACC | chrX |  | yes |  |
| Hsa-Mir-8-P2a\_3p | -0.60 | 0.16 | 6.49e-04 | 6834 | 10330 | hsa-mir-200b | MIR-8 | AAUACUG | chr1 | Epithelial Cell | yes | yes |
| Hsa-Mir-127\_3p | -1.99 | 0.22 | 3.22e-17 | 617 | 2263 | hsa-mir-127 | MIR-127 | CGGAUCC | chr14 |  | yes | yes |
| Hsa-Mir-154-P13\_5p | -1.34 | 0.15 | 3.38e-17 | 142 | 328 | hsa-mir-411 | MIR-154 | AGUAGAC | chr14 |  | yes | yes |
| Hsa-Mir-154-P9\_3p | -1.34 | 0.14 | 1.80e-18 | 107 | 253 | hsa-mir-381 | MIR-154 | AUACAAG | chr14 |  | yes | yes |
| Hsa-Mir-190-P1\_5p | -1.26 | 0.18 | 2.77e-11 | 58 | 140 | hsa-mir-190a | MIR-190 | GAUAUGU | chr15 |  | yes | yes |
| Hsa-Mir-154-P23\_3p | -1.22 | 0.14 | 1.46e-16 | 69 | 148 | hsa-mir-654 | MIR-154 | AUGUCUG | chr14 |  | yes | yes |
| Hsa-Mir-136\_3p | -1.09 | 0.15 | 5.02e-12 | 86 | 173 | hsa-mir-136 | MIR-136 | AUCAUCG | chr14 |  | yes | yes |
| Hsa-Mir-378\_3p | -0.69 | 0.14 | 2.25e-06 | 3998 | 6426 | hsa-mir-378a | MIR-378 | CUGGACU | chr5 |  | yes | yes |
| Hsa-Mir-10-P1b\_5p | -0.66 | 0.25 | 1.93e-02 | 30285 | 39286 | hsa-mir-10b | MIR-10 | ACCCUGU | chr2 |  | yes | yes |

```
# Number of downregulated miRNA
signature_mirnas$number_downregulated
```

```
## [1] 26
```

```
res_tibble <- res$res
res_tibble$miRNA <- rownames(res_tibble)
res_tibble <- as_tibble(res_tibble)

metslfc <- res_dict$tissue.type_metastasis.lung_vs_tumor.colorect$log2FoldChange
normlfc <- res_dict$tissue.type_normal.lung_vs_normal.colorect$log2FoldChange

res_tibble$LFC_adj_background <- mapply(SubtractLFC, metslfc, normlfc)

metsP <- res_dict$tissue.type_metastasis.lung_vs_tumor.colorect$padj
normP <- res_dict$tissue.type_normal.lung_vs_normal.colorect$padj

res_tibble$padj_subt_normal <- mapply(SubtractAdjP, metslfc, normlfc,metsP, normP)

res_tibble %>% select(miRNA, log2FoldChange, lfcSE, LFC_adj_background, padj_subt_normal, baseMean, stat, pvalue, padj) %>% write_csv(path = 
                      '/Users/eirikhoy/Dropbox/projects/comet_analysis/data/Deseq_result_mlu_vs_pcrc.csv')
```

#### pCRC vs PM

```
#pCRC versus PC metastasis, union of mLi and mLu normal control

column='tissue.type'
tissue_type_A <- 'metastasis.pc'
tissue_type_B <- 'tumor.colorect'
norm_adj_up       = union(dict_sig_mirna$tissue.type_normal.liver_vs_normal.colorect_up, dict_sig_mirna$tissue.type_normal.lung_vs_normal.colorect_up)
norm_adj_down     = union(dict_sig_mirna$tissue.type_normal.liver_vs_normal.colorect_down, dict_sig_mirna$tissue.type_normal.lung_vs_normal.colorect_down)
pCRC_adj_up       = union(dict_sig_mirna$tissue.type_normal.liver_vs_tumor.colorect_up, dict_sig_mirna$tissue.type_normal.lung_vs_tumor.colorect_up)
pCRC_adj_down     = union(dict_sig_mirna$tissue.type_normal.liver_vs_tumor.colorect_down, dict_sig_mirna$tissue.type_normal.lung_vs_tumor.colorect_down)

coef <- paste(column, tissue_type_A, 'vs', tissue_type_B, sep='_')
res <- DeseqResult(dds, column, coef, tissue_type_A, tissue_type_B,
                   lfc.Threshold, rpm.Threshold,
                   norm_adj_up,
                   norm_adj_down,
                   pCRC_adj_up,
                   pCRC_adj_down)

dict_sig_mirna[paste(coef, "up",   sep='_')] <- list(res$up_mirna)
dict_sig_mirna[paste(coef, "down", sep='_')] <- list(res$down_mirna)
res_res <- res$res
res_dict[coef] <- res_res
plotMA(res$res, alpha=0.05)
```

```
# Plot volcano plot
VolcanoPlot(res$res, coef, res$sig,
            res$up_mirna, res$down_mirna,
            norm_adj_up, norm_adj_down,
            pCRC_adj_up, pCRC_adj_down)
```

```
ExpressionPlot(res$res, res$rpm, coef, res$sig,
               tissue_type_A, tissue_type_B,
               res$up_mirna, res$down_mirna,
               norm_adj_up, norm_adj_down,
               pCRC_adj_up, pCRC_adj_down)
```

```
signature_mirnas <- SigList(res, dds, tissue_type_A, tissue_type_B, coef,
                            norm_adj_up, norm_adj_down, 
                            pCRC_adj_up, pCRC_adj_down)

# Print list upregulated miRNA
print("as no normal adjacent PC tissue was available, control was union of lung and liver normal adjacent")
```

```
## [1] "as no normal adjacent PC tissue was available, control was union of lung and liver normal adjacent"
```

```
signature_mirnas$up_mirna
```

Upregulated in tissue.type\_metastasis.pc\_vs\_tumor.colorect

| miRNA | LFC | lfcSE | FDR | RPM metastasis.pc | RPM tumor.colorect | miRBase\_ID | Family | Seed | Chr | Cell-Type Specific | Norm Background | pCRC Background |
| --- | --- | --- | --- | --- | --- | --- | --- | --- | --- | --- | --- | --- |
| Hsa-Mir-155\_5p | 0.70 | 0.17 | 1.59e-04 | 654 | 387 | hsa-mir-155 | MIR-155 | UAAUGCU | chr21 | c(“Lymphocyte”, “Macrophage”) |  |  |
| Hsa-Mir-506-P4a1\_3p/P4a2\_3p/P4b\_3p | 2.72 | 0.29 | 5.75e-11 | 191 | 14 | hsa-mir-509-1 | MIR-506 | GAUUGGU | chrX |  |  |  |
| Hsa-Mir-506-P3\_3p | 2.69 | 0.29 | 6.35e-09 | 112 | 6 | hsa-mir-508 | MIR-506 | GAUUGUA | chrX |  |  |  |
| Hsa-Mir-154-P9\_3p | 0.85 | 0.14 | 1.43e-08 | 455 | 253 | hsa-mir-381 | MIR-154 | AUACAAG | chr14 |  |  |  |
| Hsa-Mir-154-P36\_3p | 0.74 | 0.13 | 7.13e-07 | 330 | 195 | hsa-mir-409 | MIR-154 | AAUGUUG | chr14 |  |  |  |
| Hsa-Mir-210\_3p | 0.71 | 0.18 | 3.37e-04 | 491 | 291 | hsa-mir-210 | MIR-210 | UGUGCGU | chr11 |  |  |  |
| Hsa-Mir-1307\_5p | 0.62 | 0.17 | 8.29e-04 | 797 | 492 | hsa-mir-1307 | MIR-1307 | CGACCGG | chr10 |  |  |  |
| Hsa-Mir-127\_3p | 0.61 | 0.20 | 1.42e-02 | 2888 | 2263 | hsa-mir-127 | MIR-127 | CGGAUCC | chr14 |  |  |  |
| Hsa-Mir-191\_5p | 0.61 | 0.15 | 2.41e-04 | 20429 | 13630 | hsa-mir-191 | MIR-191 | AACGGAA | chr3 |  |  |  |
| Hsa-Mir-150\_5p | 0.70 | 0.21 | 2.45e-03 | 528 | 280 | hsa-mir-150 | MIR-150 | CUCCCAA | chr19 | Lymphocyte |  | yes |
| Hsa-Mir-154-P13\_5p | 0.94 | 0.14 | 2.08e-09 | 628 | 328 | hsa-mir-411 | MIR-154 | AGUAGAC | chr14 |  |  | yes |
| Hsa-Mir-15-P2a\_5p/P2b\_5p | 0.80 | 0.11 | 7.44e-11 | 8204 | 4522 | hsa-mir-16-1 | MIR-15 | AGCAGCA | chr13 |  |  | yes |
| Hsa-Mir-136\_3p | 0.69 | 0.14 | 1.18e-05 | 274 | 173 | hsa-mir-136 | MIR-136 | AUCAUCG | chr14 |  |  | yes |
| Hsa-Mir-335\_5p | 0.59 | 0.11 | 2.33e-06 | 232 | 150 | hsa-mir-335 | MIR-335 | CAAGAGC | chr7 | Retinal Epithelial Cell | yes | yes |
| Hsa-Mir-122\_5p | 1.45 | 0.29 | 9.08e-08 | 244 | 9 | hsa-mir-122 | MIR-122 | GGAGUGU | chr18 | Hepatocyte | yes | yes |
| Hsa-Mir-486\_5p | 0.76 | 0.22 | 2.47e-03 | 4442 | 2013 | hsa-mir-486-1 | MIR-486 | CCUGUAC | chr8 | c(“Platelet”, “Red Blood Cell”) | yes | yes |
| Hsa-Mir-126\_5p | 0.63 | 0.14 | 2.84e-05 | 5738 | 3455 | hsa-mir-126 | MIR-126 | AUUAUUA | chr9 | c(“Endothelial Cell”, “Platelet”) | yes | yes |
| Hsa-Mir-342\_3p | 0.76 | 0.16 | 1.39e-05 | 275 | 145 | hsa-mir-342 | MIR-342 | CUCACAC | chr14 | c(“Dendritic Cell”, “Lymphocyte”, “Macrophage”) | yes | yes |
| Hsa-Mir-10-P2c\_5p | 1.10 | 0.20 | 1.07e-06 | 349 | 142 | hsa-mir-99a | MIR-10 | ACCCGUA | chr21 |  | yes | yes |
| Hsa-Let-7-P1c\_5p | 0.95 | 0.18 | 1.40e-06 | 1282 | 606 | hsa-let-7c | LET-7 | GAGGUAG | chr21 |  | yes | yes |
| Hsa-Mir-10-P3c\_5p | 0.80 | 0.19 | 1.02e-04 | 497 | 239 | hsa-mir-125b-2 | MIR-10 | CCCUGAG | chr21 |  | yes | yes |
| Hsa-Mir-154-P23\_3p | 0.80 | 0.13 | 5.21e-08 | 256 | 148 | hsa-mir-654 | MIR-154 | AUGUCUG | chr14 |  | yes | yes |
| Hsa-Mir-130-P1a\_3p | 0.70 | 0.13 | 1.40e-06 | 728 | 421 | hsa-mir-130a | MIR-130 | AGUGCAA | chr11 |  | yes | yes |
| Hsa-Mir-15-P2c\_5p | 0.60 | 0.15 | 3.85e-04 | 413 | 257 | hsa-mir-195 | MIR-15 | AGCAGCA | chr17 |  | yes | yes |
| Hsa-Mir-26-P1\_5p/P2\_5p | 0.59 | 0.09 | 3.22e-09 | 52635 | 35584 | hsa-mir-26b | MIR-26 | UCAAGUA | chr2 |  | yes | yes |

```
# Number of upregulated miRNA
signature_mirnas$number_upregulated
```

```
## [1] 25
```

```
# Print list downregulated miRNA
print("as no normal adjacent PC tissue was available, control was union of lung and liver normal adjacent")
```

```
## [1] "as no normal adjacent PC tissue was available, control was union of lung and liver normal adjacent"
```

```
signature_mirnas$down_mirna
```

Downregulated in tissue.type\_metastasis.pc\_vs\_tumor.colorect

| miRNA | LFC | lfcSE | FDR | RPM metastasis.pc | RPM tumor.colorect | miRBase\_ID | Family | Seed | Chr | Cell-Type Specific | Norm Background | pCRC Background |
| --- | --- | --- | --- | --- | --- | --- | --- | --- | --- | --- | --- | --- |
| Hsa-Mir-223\_3p | -0.94 | 0.21 | 4.62e-05 | 280 | 619 | hsa-mir-223 | MIR-223 | GUCAGUU | chrX | c(“Dendritic Cell”, “Macrophage”) |  |  |
| Hsa-Mir-143\_3p | -0.86 | 0.17 | 6.37e-06 | 77753 | 148431 | hsa-mir-143 | MIR-143 | GAGAUGA | chr5 | Mesenchymal | yes | yes |
| Hsa-Mir-133-P1\_3p/P2\_3p/P3\_3p | -1.82 | 0.23 | 3.20e-14 | 25 | 116 | hsa-mir-133a-2 | MIR-133 | UUGGUCC | chr20 | c(“Skeletal Myocyte”, “Stem Cell”) | yes | yes |

```
# Number of downregulated miRNA
signature_mirnas$number_downregulated
```

```
## [1] 3
```

```
res_tibble <- res$res
res_tibble$miRNA <- rownames(res_tibble)
res_tibble <- as_tibble(res_tibble)

metslfc <- res_dict$tissue.type_metastasis.pc_vs_tumor.colorect$log2FoldChange
normlfc <- ( (res_dict$tissue.type_normal.lung_vs_normal.colorect$log2FoldChange + res_dict$tissue.type_normal.liver_vs_normal.colorect$log2FoldChange) / 2)

res_tibble$LFC_adj_background <- mapply(SubtractLFC, metslfc, normlfc)

metsP <- res_dict$tissue.type_metastasis.pc_vs_tumor.colorect$padj
normP <- ( (res_dict$tissue.type_normal.lung_vs_normal.colorect$padj - res_dict$tissue.type_normal.liver_vs_normal.colorect$padj) / 2 )

res_tibble$padj_subt_normal <- mapply(SubtractAdjP, metslfc, normlfc,metsP, normP)


res_tibble %>% select(miRNA, log2FoldChange, lfcSE, LFC_adj_background, padj_subt_normal, baseMean, stat, pvalue, padj) %>% write_csv(path = '/Users/eirikhoy/Dropbox/projects/comet_analysis/data/Deseq_result_pc_vs_pcrc.csv')
```

##### All mCRC vs pCRC

```
DeseqObject <- function(DESIGN, countdata, coldata, consensus="None", sample_type="None", Ref) {
  "
  Function to create DESeq2 object
  "
  
  dds <- DESeqDataSetFromMatrix(countData = countdata,
                                colData = coldata,
                                design = as.formula(paste("~", DESIGN)))
    # Kick out non-consensus samples
  if (!(consensus == "None")) {
    dds <- dds[, dds$paper %in% consensus]
  }

  # Kick out samples that are not bulk tissue
  if (!(sample_type == "None")) {
    dds <- dds[, dds$sample_type == sample_type]
  }

  dds$type <- relevel(dds$type, ref=ref)
  dds$type <- droplevels(dds$type)
  
  dds <- DESeq(dds,
               parallel=TRUE,
               BPPARAM=MulticoreParam(3)
               )

  return(dds)
}
```

```
SigList <- function(res, dds, tissue_type_A, tissue_type_B, coef,
                            norm_adj_up, norm_adj_down, 
                            pCRC_adj_up, pCRC_adj_down){
  "
  Function to create annotated lists of signature miRNA
  Return will print upregulated or downregulated miRNA, 
  by printing <signature_list>$up_mirna
  or          <signature_list>$down_mirna
  "
  group_A_rpm <- rowMeans(res$rpm[res$sig, dds$type == tissue_type_A])
  group_A_rpm_std <- rowSds(res$rpm[res$sig, dds$type == tissue_type_A])
  group_B_rpm <- rowMeans(res$rpm[res$sig, dds$type == tissue_type_B])
  group_B_rpm_std <- rowSds(res$rpm[res$sig, dds$type == tissue_type_B])
  lfc.deseq2  <- res$res[res$sig, ]$log2FoldChange
  lfcSE.deseq2<- res$res[res$sig, ]$lfcSE
  neg.log.10.adj.p <- -log10(res$res[res$sig, ]$padj)
  signature_mirna <- res$sig
  sig_list <- dplyr::tibble(signature_mirna, lfc.deseq2, lfcSE.deseq2,
                            group_A_rpm, #group_A_rpm_std,
                            group_B_rpm, #group_B_rpm_std,
                            neg.log.10.adj.p)
  
  # create list of upregulated mirna
  up_mirna <- sig_list %>%
    filter(lfc.deseq2 > lfc.Threshold) %>% 
    
    # Annotate which miRNA are cell markers
    mutate(
      cell_marker = ifelse(signature_mirna %in% names(cell_spec_dict_inv), cell_spec_dict_inv[signature_mirna], '')) %>%
    mutate(
      cell_marker = cell_spec(cell_marker, color = ifelse(cell_marker != '', 'white', 'black'),
                              background = ifelse(cell_marker != '', 'blue', 'white'),
                              bold = ifelse(cell_marker != '', F, F))) %>%
  # Annotate which miRNA are present in blood cells
    mutate(
      blood_cell = ifelse(signature_mirna %in% blood.cell.mirna, 'yes', '')) %>%
    mutate(
      blood_cell = cell_spec(blood_cell, color = ifelse(blood_cell == 'yes', 'white', 'black'),
                              background = ifelse(blood_cell == 'yes', 'red', 'white'),
                              bold = ifelse(blood_cell == 'yes', T, F)))

    # Annotate which miRNA are in normal_adjacent
    if (norm_adj_up != "None") {
      up_mirna <- up_mirna %>%
        mutate(
          norm_adj = ifelse(signature_mirna %in% norm_adj_up, 'yes', '')) %>%
        mutate(
          norm_adj = cell_spec(norm_adj, color = ifelse(norm_adj == 'yes', 'white', 'black'),
                               background = ifelse(norm_adj == 'yes', 'black', 'white'),
                               bold = ifelse(norm_adj == 'yes', T, F))
        )
    }
    else up_mirna$norm_adj <- "na"
    
  # Annotate which miRNA are in pCRC_adjacent
    if (pCRC_adj_up != "None") {
      up_mirna <- up_mirna %>%
        mutate(
          pCRC_adj = ifelse(signature_mirna %in% pCRC_adj_up, 'yes', '')) %>%
        mutate(
          pCRC_adj = cell_spec(pCRC_adj, color = ifelse(pCRC_adj == 'yes', 'white', 'black'),
                               background = ifelse(pCRC_adj == 'yes', 'black', 'white'),
                               bold = ifelse(pCRC_adj == 'yes', T, F))
        )
    }
    else up_mirna$pCRC_adj <- "na"

    # number of upregulated miRNA
    number_upregulated <- dim(up_mirna)[1]

  # Create kable list with annotations
    up_mirna <- up_mirna %>%
      arrange(-lfc.deseq2) %>%
      arrange(desc(cell_marker)) %>%
      arrange(pCRC_adj) %>%
      arrange(norm_adj) %>%
      kable(col.names = c("miRNA", "LFC", "lfcSE",
                          paste('RPM', tissue_type_A), #paste('std', tissue_type_A), 
                          paste('RPM', tissue_type_B), #paste('std', tissue_type_B), 
                          "-log10(adj p-value)", "cell_marker", "blood_cell", 'norm_adj', 'pCRC_adj'),
            escape = F, booktabs = F, caption = paste("Upregulated in ", coef),
            digits = c(0, 2, 2, 0, 0, 3, 2, 3, 0, 0, 0, 0)) %>%
      kable_styling(bootstrap_options = c("striped", "hover", "condensed"), full_width = T, 
                  fixed_thead = list(enabled = T)) %>%
      column_spec(2, bold = T)
    
  # create list of downregulated miRNA
  down_mirna <- sig_list %>%
    filter(lfc.deseq2 < -lfc.Threshold) %>% 
    
    # Annotate which miRNA are cell markers
    mutate(
      cell_marker = ifelse(signature_mirna %in% names(cell_spec_dict_inv), cell_spec_dict_inv[signature_mirna], '')) %>%
    mutate(
      cell_marker = cell_spec(cell_marker, color = ifelse(cell_marker != '', 'white', 'black'),
                              background = ifelse(cell_marker != '', 'blue', 'white'),
                              bold = ifelse(cell_marker != '', F, F))) %>%
  # Annotate which miRNA are present in blood cells
    mutate(
      blood_cell = ifelse(signature_mirna %in% blood.cell.mirna, 'yes', '')) %>%
    mutate(
      blood_cell = cell_spec(blood_cell, color = ifelse(blood_cell == 'yes', 'white', 'black'),
                              background = ifelse(blood_cell == 'yes', 'red', 'white'),
                              bold = ifelse(blood_cell == 'yes', T, F)))

    # Annotate which miRNA are in normal_adjacent
    if (norm_adj_down != "None") {
      down_mirna <- down_mirna %>%
        mutate(
          norm_adj = ifelse(signature_mirna %in% norm_adj_down, 'yes', '')) %>%
        mutate(
          norm_adj = cell_spec(norm_adj, color = ifelse(norm_adj == 'yes', 'white', 'black'),
                               background = ifelse(norm_adj == 'yes', 'black', 'white'),
                               bold = ifelse(norm_adj == 'yes', T, F))
        )
    }
    else down_mirna$norm_adj <- "na"

    # Annotate which miRNA are in pCRC_adjacent
    if (pCRC_adj_down != "None") {
      down_mirna <- down_mirna %>%
        mutate(
          pCRC_adj = ifelse(signature_mirna %in% pCRC_adj_down, 'yes', '')) %>%
        mutate(
          pCRC_adj = cell_spec(pCRC_adj, color = ifelse(pCRC_adj == 'yes', 'white', 'black'),
                               background = ifelse(pCRC_adj == 'yes', 'black', 'white'),
                               bold = ifelse(pCRC_adj == 'yes', T, F))
        )
    }
    else down_mirna$pCRC_adj <- "na"
  
    # number of upregulated miRNA
    number_downregulated <- dim(down_mirna)[1]

  # Create kable list with annotations    
    down_mirna <- down_mirna %>%
      arrange(lfc.deseq2) %>%
      arrange(desc(cell_marker)) %>%
      arrange(pCRC_adj) %>%
      arrange(norm_adj) %>%
      kable(col.names = c("miRNA", "LFC", "lfcSE",
                          paste('RPM', tissue_type_A), #paste('std', tissue_type_A), 
                          paste('RPM', tissue_type_B), #paste('std', tissue_type_B), 
                          "-log10(adj p-value)", "cell_marker", "blood_cell", 'norm_adj', 'pCRC_adj'),
            escape = F, booktabs = F, caption = paste("Downregulated in ", coef),
            digits = c(0, 2, 2, 0, 0, 3, 2, 3, 0, 0, 0, 0)) %>%
      kable_styling(bootstrap_options = c("striped", "hover", "condensed"), full_width = T, 
                  fixed_thead = list(enabled = T)) %>%
      column_spec(2, bold = T)
    

  # Function return is to print kable, either upregulated or downregulated miRNA
  return_list = list("up_mirna" = up_mirna, "down_mirna" = down_mirna, 
                     "number_upregulated" = number_upregulated, 
                     "number_downregulated" = number_downregulated)
  return(return_list)
}
```

```
design <- as.formula(~ type)
ref <- 'tumor'
dds <- DeseqObject(design, countdata, sampleinfo, "None", "None", ref)
#
```

```
column='type'
tissue_type_A <- 'metastasis'
tissue_type_B <- 'tumor'
norm_adj_up       = union(dict_sig_mirna$tissue.type_normal.liver_vs_normal.colorect_up, dict_sig_mirna$tissue.type_normal.lung_vs_normal.colorect_up)
norm_adj_down     = union(dict_sig_mirna$tissue.type_normal.liver_vs_normal.colorect_down, dict_sig_mirna$tissue.type_normal.lung_vs_normal.colorect_down)
pCRC_adj_up       = union(dict_sig_mirna$tissue.type_normal.liver_vs_tumor.colorect_up, dict_sig_mirna$tissue.type_normal.lung_vs_tumor.colorect_up)
pCRC_adj_down     = union(dict_sig_mirna$tissue.type_normal.liver_vs_tumor.colorect_down, dict_sig_mirna$tissue.type_normal.lung_vs_tumor.colorect_down)

coef <- paste(column, tissue_type_A, 'vs', tissue_type_B, sep='_')
res <- DeseqResult(dds, column, coef, tissue_type_A, tissue_type_B,
                   lfc.Threshold, rpm.Threshold,
                   norm_adj_up,
                   norm_adj_down,
                   pCRC_adj_up,
                   pCRC_adj_down)
```

```
## using 'normal' for LFC shrinkage, the Normal prior from Love et al (2014).
## 
## Note that type='apeglm' and type='ashr' have shown to have less bias than type='normal'.
## See ?lfcShrink for more details on shrinkage type, and the DESeq2 vignette.
## Reference: https://doi.org/10.1093/bioinformatics/bty895
```

```
## Warning in if (!(norm_adj_up == "None")) {: the condition has length > 1 and
## only the first element will be used
```

```
## Warning in if (!(norm_adj_down == "None")) {: the condition has length > 1 and
## only the first element will be used
```

```
## Warning in if (!(pCRC_adj_up == "None")) {: the condition has length > 1 and
## only the first element will be used
```

```
## Warning in if (!(pCRC_adj_down == "None")) {: the condition has length > 1 and
## only the first element will be used
```

```
dict_sig_mirna[paste(coef, "up",   sep='_')] <- list(res$up_mirna)
dict_sig_mirna[paste(coef, "down", sep='_')] <- list(res$down_mirna)
res_res <- res$res
res_dict[coef] <- res_res
```

```
## Warning in `[<-`(`*tmp*`, coef, value = new("DESeqResults", priorInfo = list(:
## implicit list embedding of S4 objects is deprecated
```

```
plotMA(res$res, alpha=0.05)
```

```
# Plot volcano plot
VolcanoPlot(res$res, coef, res$sig,
            res$up_mirna, res$down_mirna,
            norm_adj_up, norm_adj_down,
            pCRC_adj_up, pCRC_adj_down)
```

```
## Warning in if (norm_adj_up != "None" & length(up_mirna > 0)) {: the condition
## has length > 1 and only the first element will be used
```

```
## Warning in if (norm_adj_down != "None" & length(down_mirna > 0)) {: the
## condition has length > 1 and only the first element will be used
```

```
## Warning in if (pCRC_adj_up != "None" & length(up_mirna > 0)) {: the condition
## has length > 1 and only the first element will be used
```

```
## Warning in if (pCRC_adj_down != "None" & length(down_mirna > 0)) {: the
## condition has length > 1 and only the first element will be used
```

```
#ExpressionPlot(res$res, res$rpm, coef, res$sig,
#               tissue_type_A, tissue_type_B,
#               res$up_mirna, res$down_mirna,
#               norm_adj_up, norm_adj_down,
#               pCRC_adj_up, pCRC_adj_down)

signature_mirnas <- SigList(res, dds, tissue_type_A, tissue_type_B, coef,
                            norm_adj_up, norm_adj_down, 
                            pCRC_adj_up, pCRC_adj_down)
```

```
## Warning in if (norm_adj_up != "None") {: the condition has length > 1 and only
## the first element will be used
```

```
## Warning in if (pCRC_adj_up != "None") {: the condition has length > 1 and only
## the first element will be used
```

```
## Warning in if (norm_adj_down != "None") {: the condition has length > 1 and only
## the first element will be used
```

```
## Warning in if (pCRC_adj_down != "None") {: the condition has length > 1 and only
## the first element will be used
```

```
# Print list upregulated miRNA
print("control was union of lung and liver normal adjacent")
```

```
## [1] "control was union of lung and liver normal adjacent"
```

```
signature_mirnas$up_mirna
```

Upregulated in type\_metastasis\_vs\_tumor

| miRNA | LFC | lfcSE | RPM metastasis | RPM tumor | -log10(adj p-value) | cell\_marker | blood\_cell | norm\_adj | pCRC\_adj |
| --- | --- | --- | --- | --- | --- | --- | --- | --- | --- |
| Hsa-Mir-210\_3p | 1.10 | 0.13 | 625 | 291 | 14.551 |  |  |  |  |
| Hsa-Mir-1307\_5p | 0.72 | 0.12 | 837 | 492 | 7.519 |  |  |  |  |
| Hsa-Mir-191\_5p | 0.72 | 0.11 | 23467 | 13630 | 9.458 |  |  |  |  |
| Hsa-Mir-8-P1b\_3p | 0.66 | 0.17 | 12715 | 7518 | 3.068 |  |  |  |  |
| Hsa-Mir-10-P1a\_5p | 0.59 | 0.15 | 140769 | 97123 | 3.469 |  |  |  |  |
| Hsa-Mir-150\_5p | 1.05 | 0.16 | 664 | 280 | 9.458 | Lymphocyte |  |  | yes |
| Hsa-Mir-15-P2a\_5p/P2b\_5p | 0.74 | 0.08 | 7988 | 4522 | 18.164 |  |  |  | yes |
| Hsa-Mir-331\_3p | 0.70 | 0.12 | 104 | 60 | 7.519 |  |  |  | yes |
| Hsa-Mir-335\_5p | 0.76 | 0.09 | 262 | 150 | 15.670 | Retinal Epithelial Cell |  | yes | yes |
| Hsa-Mir-122\_5p | 1.56 | 0.31 | 1459 | 9 | 24.422 | Hepatocyte |  | yes | yes |
| Hsa-Mir-126\_5p | 0.92 | 0.12 | 7434 | 3455 | 12.397 | c(“Endothelial Cell”, “Platelet”) |  | yes | yes |
| Hsa-Mir-342\_3p | 1.46 | 0.13 | 472 | 145 | 27.005 | c(“Dendritic Cell”, “Lymphocyte”, “Macrophage”) |  | yes | yes |
| Hsa-Mir-34-P2b\_5p | 2.61 | 0.21 | 332 | 33 | 19.564 |  |  | yes | yes |
| Hsa-Mir-30-P1a\_5p | 0.93 | 0.13 | 7100 | 3165 | 10.616 |  |  | yes | yes |
| Hsa-Mir-15-P2c\_5p | 0.87 | 0.12 | 537 | 257 | 10.661 |  |  | yes | yes |
| Hsa-Mir-10-P3c\_5p | 0.82 | 0.15 | 514 | 239 | 6.775 |  |  | yes | yes |
| Hsa-Mir-26-P1\_5p/P2\_5p | 0.71 | 0.07 | 60843 | 35584 | 23.244 |  |  | yes | yes |
| Hsa-Mir-338-P1\_3p | 0.70 | 0.16 | 169 | 98 | 4.263 |  |  | yes | yes |
| Hsa-Mir-10-P2c\_5p | 0.70 | 0.17 | 264 | 142 | 3.839 |  |  | yes | yes |

```
# Number of upregulated miRNA
signature_mirnas$number_upregulated
```

```
## [1] 19
```

```
# Print list downregulated miRNA
print("control was union of lung and liver normal adjacent")
```

```
## [1] "control was union of lung and liver normal adjacent"
```

```
signature_mirnas$down_mirna
```

Downregulated in type\_metastasis\_vs\_tumor

| miRNA | LFC | lfcSE | RPM metastasis | RPM tumor | -log10(adj p-value) | cell\_marker | blood\_cell | norm\_adj | pCRC\_adj |
| --- | --- | --- | --- | --- | --- | --- | --- | --- | --- |
| Hsa-Mir-7-P1\_5p/P2\_5p/P3\_5p | -0.60 | 0.18 | 112 | 199 | 2.657 | c(“Islet Cell”, “Neural”) |  |  | yes |
| Hsa-Mir-31\_5p | -0.68 | 0.23 | 339 | 627 | 2.235 |  |  |  | yes |
| Hsa-Mir-143\_3p | -1.08 | 0.14 | 72990 | 148431 | 13.050 | Mesenchymal |  | yes | yes |
| Hsa-Mir-133-P1\_3p/P2\_3p/P3\_3p | -1.80 | 0.19 | 33 | 116 | 18.097 | c(“Skeletal Myocyte”, “Stem Cell”) |  | yes | yes |

```
# Number of downregulated miRNA
signature_mirnas$number_downregulated
```

```
## [1] 4
```

```
res_dict
```

```
## $tissue.type_normal.liver_vs_normal.colorect
## log2 fold change (MAP): tissue.type normal.liver vs normal.colorect 
## Wald test p-value: tissue.type normal.liver vs normal.colorect 
## DataFrame with 389 rows and 6 columns
##                                          baseMean     log2FoldChange
##                                         <numeric>          <numeric>
## Hsa-Let-7-P1a_5p/P2a1_5p/P2a2_5p 89174.7542403281 -0.114702438165718
## Hsa-Let-7-P1b_5p                 3359.93816553835 -0.320114053507846
## Hsa-Let-7-P1c_5p                 3992.69006418194   3.18371450404926
## Hsa-Let-7-P2a3_5p                52000.6252125236  0.135005279299899
## Hsa-Let-7-P2b1_5p                9751.60360980849  0.321769922951614
## ...                                           ...                ...
## Hsa-Mir-95-P2_3p                 390.216333625945 -0.259423173236495
## Hsa-Mir-95-P3_5p                 3.95328293251241  0.578777309350614
## Hsa-Mir-96-P1_5p                 202.656949570096  -2.86706387684201
## Hsa-Mir-96-P2_5p                 27209.4573430933  -2.64828835135873
## Hsa-Mir-96-P3_5p                 3344.18853507094  -3.56720648489265
##                                              lfcSE               stat
##                                          <numeric>          <numeric>
## Hsa-Let-7-P1a_5p/P2a1_5p/P2a2_5p 0.137100418774597 -0.876702398466493
## Hsa-Let-7-P1b_5p                 0.223907107455248   -1.4689025908534
## Hsa-Let-7-P1c_5p                 0.279081137786442   11.1516794079232
## Hsa-Let-7-P2a3_5p                0.120493686572101   1.11711255877344
## Hsa-Let-7-P2b1_5p                0.122630483027464   2.60055162322238
## ...                                            ...                ...
## Hsa-Mir-95-P2_3p                  0.19641408635223  -1.08552839444029
## Hsa-Mir-95-P3_5p                 0.384387609595463   1.62826495421394
## Hsa-Mir-96-P1_5p                 0.333638316705081  -7.80833454343775
## Hsa-Mir-96-P2_5p                 0.243241112732279  -10.2148298811257
## Hsa-Mir-96-P3_5p                 0.324276388539775  -10.0394868651728
##                                                pvalue                 padj
##                                             <numeric>            <numeric>
## Hsa-Let-7-P1a_5p/P2a1_5p/P2a2_5p    0.380648303687109    0.543582632940632
## Hsa-Let-7-P1b_5p                    0.141859211826177    0.240787346389169
## Hsa-Let-7-P1c_5p                 7.02674679240312e-29 1.59961824038824e-27
## Hsa-Let-7-P2a3_5p                   0.263946201391012    0.404220147427516
## Hsa-Let-7-P2b1_5p                  0.0093074014267008   0.0220979408106332
## ...                                               ...                  ...
## Hsa-Mir-95-P2_3p                    0.277687694717835    0.421431913160009
## Hsa-Mir-95-P3_5p                    0.103468717163438     0.18284197964498
## Hsa-Mir-96-P1_5p                 5.79486025768406e-15 5.33954980886603e-14
## Hsa-Mir-96-P2_5p                  1.7017739490561e-24 2.99357508311232e-23
## Hsa-Mir-96-P3_5p                 1.02204432533755e-23 1.71970066915492e-22
## 
## $tissue.type_normal.lung_vs_normal.colorect
## log2 fold change (MAP): tissue.type normal.lung vs normal.colorect 
## Wald test p-value: tissue.type normal.lung vs normal.colorect 
## DataFrame with 389 rows and 6 columns
##                                          baseMean      log2FoldChange
##                                         <numeric>           <numeric>
## Hsa-Let-7-P1a_5p/P2a1_5p/P2a2_5p 89174.7542403281   0.159488915677012
## Hsa-Let-7-P1b_5p                 3359.93816553835   0.197555867999725
## Hsa-Let-7-P1c_5p                 3992.69006418194    1.96610542233694
## Hsa-Let-7-P2a3_5p                52000.6252125236   0.289175178805092
## Hsa-Let-7-P2b1_5p                9751.60360980849    1.10414120876771
## ...                                           ...                 ...
## Hsa-Mir-95-P2_3p                 390.216333625945   0.165907582077109
## Hsa-Mir-95-P3_5p                 3.95328293251241 -0.0273303033157357
## Hsa-Mir-96-P1_5p                 202.656949570096  -0.415869888949985
## Hsa-Mir-96-P2_5p                 27209.4573430933  -0.107488420248239
## Hsa-Mir-96-P3_5p                 3344.18853507094  -0.456955953013122
##                                              lfcSE                stat
##                                          <numeric>           <numeric>
## Hsa-Let-7-P1a_5p/P2a1_5p/P2a2_5p 0.171252727891764   0.886346895351987
## Hsa-Let-7-P1b_5p                 0.280263432235887   0.631339921419183
## Hsa-Let-7-P1c_5p                 0.350196583213482    5.61986491346816
## Hsa-Let-7-P2a3_5p                 0.15044933221432    1.91622249772382
## Hsa-Let-7-P2b1_5p                0.153065517284977    7.17242652538056
## ...                                            ...                 ...
## Hsa-Mir-95-P2_3p                 0.243046643416424   0.843104626057063
## Hsa-Mir-95-P3_5p                 0.458035265321474   0.143432621264712
## Hsa-Mir-96-P1_5p                 0.388545838443054  -0.690134766070179
## Hsa-Mir-96-P2_5p                 0.304517199045329 -0.0704700708063357
## Hsa-Mir-96-P3_5p                 0.404724536391786  -0.772763885607241
##                                                pvalue                 padj
##                                             <numeric>            <numeric>
## Hsa-Let-7-P1a_5p/P2a1_5p/P2a2_5p    0.375430626009251    0.530261504618906
## Hsa-Let-7-P1b_5p                     0.52781828944235    0.684422033196025
## Hsa-Let-7-P1c_5p                 1.91106824923059e-08 1.57358172862179e-07
## Hsa-Let-7-P2a3_5p                  0.0553367808976523    0.104465044914105
## Hsa-Let-7-P2b1_5p                7.36799615073784e-13 1.05607944827242e-11
## ...                                               ...                  ...
## Hsa-Mir-95-P2_3p                    0.399169931746439    0.555679005704575
## Hsa-Mir-95-P3_5p                    0.885948521274591    0.960397976843884
## Hsa-Mir-96-P1_5p                    0.490109441851604    0.651795030916051
## Hsa-Mir-96-P2_5p                    0.943819521347161     0.98187676011116
## Hsa-Mir-96-P3_5p                    0.439662130177162    0.603366114817595
## 
## $tissue.type_tumor.colorect_vs_normal.colorect
## log2 fold change (MAP): tissue.type tumor.colorect vs normal.colorect 
## Wald test p-value: tissue.type tumor.colorect vs normal.colorect 
## DataFrame with 389 rows and 6 columns
##                                          baseMean      log2FoldChange
##                                         <numeric>           <numeric>
## Hsa-Let-7-P1a_5p/P2a1_5p/P2a2_5p 89174.7542403281  -0.248058505837001
## Hsa-Let-7-P1b_5p                 3359.93816553835  -0.129712872859404
## Hsa-Let-7-P1c_5p                 3992.69006418194 -0.0894387113372064
## Hsa-Let-7-P2a3_5p                52000.6252125236   0.161274363696562
## Hsa-Let-7-P2b1_5p                9751.60360980849  -0.129885530327206
## ...                                           ...                 ...
## Hsa-Mir-95-P2_3p                 390.216333625945    1.16639047991208
## Hsa-Mir-95-P3_5p                 3.95328293251241   0.128041251542255
## Hsa-Mir-96-P1_5p                 202.656949570096    1.99605999628903
## Hsa-Mir-96-P2_5p                 27209.4573430933     1.8371528804308
## Hsa-Mir-96-P3_5p                 3344.18853507094     1.5305097819346
##                                               lfcSE               stat
##                                           <numeric>          <numeric>
## Hsa-Let-7-P1a_5p/P2a1_5p/P2a2_5p   0.09974589771869  -2.50880006172293
## Hsa-Let-7-P1b_5p                  0.160828091638605 -0.872616006654668
## Hsa-Let-7-P1c_5p                  0.198255228124062  -0.14744126618236
## Hsa-Let-7-P2a3_5p                0.0878077644317027   1.81863066456019
## Hsa-Let-7-P2b1_5p                0.0893412951279261  -1.43703312496472
## ...                                             ...                ...
## Hsa-Mir-95-P2_3p                  0.140554605668826   8.28848119000958
## Hsa-Mir-95-P3_5p                  0.264177931179274  0.737283525185868
## Hsa-Mir-96-P1_5p                  0.217577914511228   8.93516022919441
## Hsa-Mir-96-P2_5p                  0.173955103251303   10.4458859814953
## Hsa-Mir-96-P3_5p                  0.226074853962932   6.65702796429079
##                                                pvalue                 padj
##                                             <numeric>            <numeric>
## Hsa-Let-7-P1a_5p/P2a1_5p/P2a2_5p   0.0121142030924821   0.0250889779893481
## Hsa-Let-7-P1b_5p                     0.38287241295549    0.480492200364671
## Hsa-Let-7-P1c_5p                    0.882783735719796    0.926800653290313
## Hsa-Let-7-P2a3_5p                  0.0689677962587844    0.122143900850838
## Hsa-Let-7-P2b1_5p                   0.150708581526998    0.225175837320617
## ...                                               ...                  ...
## Hsa-Mir-95-P2_3p                  1.1470369823237e-16 1.40234521387317e-15
## Hsa-Mir-95-P3_5p                    0.460949948804283    0.561736432787213
## Hsa-Mir-96-P1_5p                 4.06588568948011e-19 8.56094820173867e-18
## Hsa-Mir-96-P2_5p                 1.53019828709604e-25 6.44383500899331e-24
## Hsa-Mir-96-P3_5p                 2.79420040427442e-11 1.96111472818519e-10
## 
## $tissue.type_normal.liver_vs_tumor.colorect
## log2 fold change (MAP): tissue.type normal.liver vs tumor.colorect 
## Wald test p-value: tissue.type normal.liver vs tumor.colorect 
## DataFrame with 389 rows and 6 columns
##                                          baseMean      log2FoldChange
##                                         <numeric>           <numeric>
## Hsa-Let-7-P1a_5p/P2a1_5p/P2a2_5p 89174.7542403281   0.134516855928531
## Hsa-Let-7-P1b_5p                 3359.93816553835   -0.18431192252938
## Hsa-Let-7-P1c_5p                 3992.69006418194    3.24635633273589
## Hsa-Let-7-P2a3_5p                52000.6252125236 -0.0245108495144203
## Hsa-Let-7-P2b1_5p                9751.60360980849   0.451258084717704
## ...                                           ...                 ...
## Hsa-Mir-95-P2_3p                 390.216333625945   -1.41840794320829
## Hsa-Mir-95-P3_5p                 3.95328293251241   0.414413322947411
## Hsa-Mir-96-P1_5p                 202.656949570096   -4.81237877419974
## Hsa-Mir-96-P2_5p                 27209.4573430933   -4.46595236546167
## Hsa-Mir-96-P3_5p                 3344.18853507094   -5.07557947770261
##                                               lfcSE               stat
##                                           <numeric>          <numeric>
## Hsa-Let-7-P1a_5p/P2a1_5p/P2a2_5p   0.11131421654252   1.19461982343763
## Hsa-Let-7-P1b_5p                  0.184409102642062  -1.03024741017946
## Hsa-Let-7-P1c_5p                  0.232675048404373    13.987286383856
## Hsa-Let-7-P2a3_5p                0.0976442188709745 -0.267757085486139
## Hsa-Let-7-P2b1_5p                0.0994097267910815   4.53746602977892
## ...                                             ...                ...
## Hsa-Mir-95-P2_3p                  0.161409886777642    -8.797157721989
## Hsa-Mir-95-P3_5p                  0.332664401974054   1.34224159185026
## Hsa-Mir-96-P1_5p                  0.288868128705996  -16.6454924674176
## Hsa-Mir-96-P2_5p                  0.201189454623787  -22.1766896561084
## Hsa-Mir-96-P3_5p                  0.274372198803725  -18.4172382155403
##                                                 pvalue                  padj
##                                              <numeric>             <numeric>
## Hsa-Let-7-P1a_5p/P2a1_5p/P2a2_5p     0.232235600538294     0.301594555061476
## Hsa-Let-7-P1b_5p                     0.302893879225991     0.381823880327226
## Hsa-Let-7-P1c_5p                  1.86389083593418e-44  3.13619892828925e-43
## Hsa-Let-7-P2a3_5p                    0.788886305673403     0.867326705385247
## Hsa-Let-7-P2b1_5p                 5.69341978961745e-06   1.4887523368797e-05
## ...                                                ...                   ...
## Hsa-Mir-95-P2_3p                  1.40325015866435e-18  7.75796873433006e-18
## Hsa-Mir-95-P3_5p                     0.179517674775297     0.240392180408443
## Hsa-Mir-96-P1_5p                  3.26270971024334e-62  7.89167911165108e-61
## Hsa-Mir-96-P2_5p                 5.76691295122432e-109 4.46359062424763e-107
## Hsa-Mir-96-P3_5p                  9.55562651928938e-76  2.84463650997307e-74
## 
## $tissue.type_normal.lung_vs_tumor.colorect
## log2 fold change (MAP): tissue.type normal.lung vs tumor.colorect 
## Wald test p-value: tissue.type normal.lung vs tumor.colorect 
## DataFrame with 389 rows and 6 columns
##                                          baseMean     log2FoldChange
##                                         <numeric>          <numeric>
## Hsa-Let-7-P1a_5p/P2a1_5p/P2a2_5p 89174.7542403281  0.408125846033093
## Hsa-Let-7-P1b_5p                 3359.93816553835  0.332599900506941
## Hsa-Let-7-P1c_5p                 3992.69006418194     2.020544361604
## Hsa-Let-7-P2a3_5p                52000.6252125236  0.129817388788356
## Hsa-Let-7-P2b1_5p                9751.60360980849   1.23298675588116
## ...                                           ...                ...
## Hsa-Mir-95-P2_3p                 390.216333625945 -0.987860472334413
## Hsa-Mir-95-P3_5p                 3.95328293251241 -0.191008452152824
## Hsa-Mir-96-P1_5p                 202.656949570096   -2.3341261742075
## Hsa-Mir-96-P2_5p                 27209.4573430933  -1.90878481829015
## Hsa-Mir-96-P3_5p                 3344.18853507094  -1.93676271625054
##                                              lfcSE               stat
##                                          <numeric>          <numeric>
## Hsa-Let-7-P1a_5p/P2a1_5p/P2a2_5p 0.151440987364688   2.68337077689517
## Hsa-Let-7-P1b_5p                 0.249992618790543   1.30066611269162
## Hsa-Let-7-P1c_5p                 0.314717268191971   6.48763117983787
## Hsa-Let-7-P2a3_5p                0.132884439212234  0.963563940574538
## Hsa-Let-7-P2b1_5p                0.135208009802693   9.11432613077411
## ...                                            ...                ...
## Hsa-Mir-95-P2_3p                 0.215830901283507  -4.59605232940141
## Hsa-Mir-95-P3_5p                 0.415244927482672 -0.364000003055949
## Hsa-Mir-96-P1_5p                 0.350565916495824  -6.75367079090678
## Hsa-Mir-96-P2_5p                 0.272289955724401  -7.05756424076837
## Hsa-Mir-96-P3_5p                 0.366207615281681  -5.32493458906083
##                                                pvalue                 padj
##                                             <numeric>            <numeric>
## Hsa-Let-7-P1a_5p/P2a1_5p/P2a2_5p  0.00728841357441722   0.0138946603610811
## Hsa-Let-7-P1b_5p                    0.193372766419832    0.281397733924404
## Hsa-Let-7-P1c_5p                 8.71964116059556e-11 4.62260428650751e-10
## Hsa-Let-7-P2a3_5p                   0.335264592721268    0.444340401997023
## Hsa-Let-7-P2b1_5p                7.91608806736923e-20 9.88234220023191e-19
## ...                                               ...                  ...
## Hsa-Mir-95-P2_3p                  4.3057060677156e-06 1.35472215301296e-05
## Hsa-Mir-95-P3_5p                    0.715858007098838    0.805340257986192
## Hsa-Mir-96-P1_5p                 1.44150633672609e-11 8.45246897443934e-11
## Hsa-Mir-96-P2_5p                 1.69446438026879e-12 1.11145375451529e-11
## Hsa-Mir-96-P3_5p                 1.00989374164518e-07   3.947766444613e-07
## 
## $tissue.type_metastasis.liver_vs_tumor.colorect
## log2 fold change (MAP): tissue.type metastasis.liver vs tumor.colorect 
## Wald test p-value: tissue.type metastasis.liver vs tumor.colorect 
## DataFrame with 389 rows and 6 columns
##                                          baseMean      log2FoldChange
##                                         <numeric>           <numeric>
## Hsa-Let-7-P1a_5p/P2a1_5p/P2a2_5p 89174.7542403281 -0.0291372578148132
## Hsa-Let-7-P1b_5p                 3359.93816553835 -0.0384102612949004
## Hsa-Let-7-P1c_5p                 3992.69006418194   0.165919575225585
## Hsa-Let-7-P2a3_5p                52000.6252125236 -0.0667790393731562
## Hsa-Let-7-P2b1_5p                9751.60360980849  -0.022055090655077
## ...                                           ...                 ...
## Hsa-Mir-95-P2_3p                 390.216333625945   0.197986570574988
## Hsa-Mir-95-P3_5p                 3.95328293251241    0.87517582162084
## Hsa-Mir-96-P1_5p                 202.656949570096  -0.153456519278348
## Hsa-Mir-96-P2_5p                 27209.4573430933  0.0668521603991884
## Hsa-Mir-96-P3_5p                 3344.18853507094   0.480378406060166
##                                               lfcSE               stat
##                                           <numeric>          <numeric>
## Hsa-Let-7-P1a_5p/P2a1_5p/P2a2_5p  0.087811822186451  -0.34524409879388
## Hsa-Let-7-P1b_5p                  0.143352612343951 -0.309464165623311
## Hsa-Let-7-P1c_5p                  0.178645206267475   1.08002602868534
## Hsa-Let-7-P2a3_5p                0.0771691545429361 -0.884124420143461
## Hsa-Let-7-P2b1_5p                0.0785334182092049 -0.274463101822189
## ...                                             ...                ...
## Hsa-Mir-95-P2_3p                  0.123880822993862   1.53530515265106
## Hsa-Mir-95-P3_5p                  0.226325334424462   3.88027612923505
## Hsa-Mir-96-P1_5p                  0.196070125088779 -0.994964941991442
## Hsa-Mir-96-P2_5p                  0.155589993744074  0.303386567615509
## Hsa-Mir-96-P3_5p                  0.205702483629426   2.17135483299889
##                                                pvalue                 padj
##                                             <numeric>            <numeric>
## Hsa-Let-7-P1a_5p/P2a1_5p/P2a2_5p    0.729910868456285    0.886990854282231
## Hsa-Let-7-P1b_5p                    0.756968466884636    0.898589554206931
## Hsa-Let-7-P1c_5p                    0.280130589584733    0.477579463300844
## Hsa-Let-7-P2a3_5p                   0.376629052036082    0.589523977412743
## Hsa-Let-7-P2b1_5p                   0.783728755707429     0.90746054828939
## ...                                               ...                  ...
## Hsa-Mir-95-P2_3p                    0.124708889065693    0.283896118049548
## Hsa-Mir-95-P3_5p                 0.000104337942960983 0.000776515075498082
## Hsa-Mir-96-P1_5p                     0.31975331550125    0.522757414533685
## Hsa-Mir-96-P2_5p                    0.761595281084944    0.898589554206931
## Hsa-Mir-96-P3_5p                   0.0299043603592752   0.0972519954541135
## 
## $tissue.type_metastasis.lung_vs_tumor.colorect
## log2 fold change (MAP): tissue.type metastasis.lung vs tumor.colorect 
## Wald test p-value: tissue.type metastasis.lung vs tumor.colorect 
## DataFrame with 389 rows and 6 columns
##                                          baseMean     log2FoldChange
##                                         <numeric>          <numeric>
## Hsa-Let-7-P1a_5p/P2a1_5p/P2a2_5p 89174.7542403281 -0.371287975266059
## Hsa-Let-7-P1b_5p                 3359.93816553835  -0.86478570992646
## Hsa-Let-7-P1c_5p                 3992.69006418194  0.135121670461325
## Hsa-Let-7-P2a3_5p                52000.6252125236 -0.362077135000897
## Hsa-Let-7-P2b1_5p                9751.60360980849 0.0403928595342307
## ...                                           ...                ...
## Hsa-Mir-95-P2_3p                 390.216333625945  0.244457301347207
## Hsa-Mir-95-P3_5p                 3.95328293251241  0.770986123093289
## Hsa-Mir-96-P1_5p                 202.656949570096 -0.746540861417062
## Hsa-Mir-96-P2_5p                 27209.4573430933  -0.78147625945163
## Hsa-Mir-96-P3_5p                 3344.18853507094 -0.326709592212615
##                                               lfcSE              stat
##                                           <numeric>         <numeric>
## Hsa-Let-7-P1a_5p/P2a1_5p/P2a2_5p 0.0959152266130106 -3.87096089411713
## Hsa-Let-7-P1b_5p                  0.156536850438983 -5.51447766923507
## Hsa-Let-7-P1c_5p                  0.194969507259272 0.835611129624883
## Hsa-Let-7-P2a3_5p                0.0842935080279992 -4.30344211748187
## Hsa-Let-7-P2b1_5p                0.0857694713451329 0.474626440937984
## ...                                             ...               ...
## Hsa-Mir-95-P2_3p                  0.135123404458061  1.75056733087186
## Hsa-Mir-95-P3_5p                   0.24193629758336  3.22815141458786
## Hsa-Mir-96-P1_5p                  0.213973544808363 -3.63856405996494
## Hsa-Mir-96-P2_5p                  0.169868922201866 -4.66162915975394
## Hsa-Mir-96-P3_5p                  0.224439059665663 -1.53101854305705
##                                                pvalue                 padj
##                                             <numeric>            <numeric>
## Hsa-Let-7-P1a_5p/P2a1_5p/P2a2_5p   0.0001084071837261  0.00040731631166991
## Hsa-Let-7-P1b_5p                 3.49817317913999e-08  2.5543264534475e-07
## Hsa-Let-7-P1c_5p                    0.403373705499327    0.532783699755084
## Hsa-Let-7-P2a3_5p                1.68164797190917e-05 7.65644429563353e-05
## Hsa-Let-7-P2b1_5p                   0.635053256632943    0.757086098142951
## ...                                               ...                  ...
## Hsa-Mir-95-P2_3p                   0.0800204667405637     0.14746628870761
## Hsa-Mir-95-P3_5p                  0.00124593006501447  0.00359832041164626
## Hsa-Mir-96-P1_5p                 0.000274162436246517  0.00098241539655002
## Hsa-Mir-96-P2_5p                 3.13716090303122e-06 1.57672892139361e-05
## Hsa-Mir-96-P3_5p                    0.125764809854213    0.211612962667741
## 
## $tissue.type_metastasis.pc_vs_tumor.colorect
## log2 fold change (MAP): tissue.type metastasis.pc vs tumor.colorect 
## Wald test p-value: tissue.type metastasis.pc vs tumor.colorect 
## DataFrame with 389 rows and 6 columns
##                                          baseMean      log2FoldChange
##                                         <numeric>           <numeric>
## Hsa-Let-7-P1a_5p/P2a1_5p/P2a2_5p 89174.7542403281  -0.194926960102631
## Hsa-Let-7-P1b_5p                 3359.93816553835  -0.170006997664929
## Hsa-Let-7-P1c_5p                 3992.69006418194   0.950788049188073
## Hsa-Let-7-P2a3_5p                52000.6252125236  -0.235739655653004
## Hsa-Let-7-P2b1_5p                9751.60360980849 -0.0275610759199226
## ...                                           ...                 ...
## Hsa-Mir-95-P2_3p                 390.216333625945  -0.391736823339089
## Hsa-Mir-95-P3_5p                 3.95328293251241   0.193480385452117
## Hsa-Mir-96-P1_5p                 202.656949570096  -0.537251798926674
## Hsa-Mir-96-P2_5p                 27209.4573430933  -0.144468039225267
## Hsa-Mir-96-P3_5p                 3344.18853507094  -0.149480720489416
##                                               lfcSE               stat
##                                           <numeric>          <numeric>
## Hsa-Let-7-P1a_5p/P2a1_5p/P2a2_5p 0.0919497346955033  -2.12106919715872
## Hsa-Let-7-P1b_5p                  0.146429745186075  -1.18446872666545
## Hsa-Let-7-P1c_5p                  0.178397588170818   5.34338828103208
## Hsa-Let-7-P2a3_5p                  0.08107727951498  -2.91425069775732
## Hsa-Let-7-P2b1_5p                0.0824764385946126  -0.32740564020296
## ...                                             ...                ...
## Hsa-Mir-95-P2_3p                  0.128019210385648  -3.07056591636714
## Hsa-Mir-95-P3_5p                  0.222349699850271  0.967689700538171
## Hsa-Mir-96-P1_5p                  0.193381569174697   -2.9161157752392
## Hsa-Mir-96-P2_5p                  0.157787812752101  -1.01238717153534
## Hsa-Mir-96-P3_5p                  0.201177138993338 -0.825123443737066
##                                                pvalue                 padj
##                                             <numeric>            <numeric>
## Hsa-Let-7-P1a_5p/P2a1_5p/P2a2_5p   0.0339159796632756   0.0901196031052751
## Hsa-Let-7-P1b_5p                    0.236227568744681    0.370368857833221
## Hsa-Let-7-P1c_5p                 9.12250626572605e-08 1.40060447984391e-06
## Hsa-Let-7-P2a3_5p                 0.00356543456685069   0.0141511415829511
## Hsa-Let-7-P2b1_5p                   0.743361101075885    0.848252544785979
## ...                                               ...                  ...
## Hsa-Mir-95-P2_3p                  0.00213653519975517    0.009575796316975
## Hsa-Mir-95-P3_5p                    0.333199363429196     0.48229635484693
## Hsa-Mir-96-P1_5p                  0.00354418957989829   0.0141511415829511
## Hsa-Mir-96-P2_5p                    0.311352969649893     0.46861091937278
## Hsa-Mir-96-P3_5p                    0.409301511166841    0.561003252050181
## 
## $type_metastasis_vs_tumor
## log2 fold change (MAP): type metastasis vs tumor 
## Wald test p-value: type metastasis vs tumor 
## DataFrame with 389 rows and 6 columns
##                                          baseMean       log2FoldChange
##                                         <numeric>            <numeric>
## Hsa-Let-7-P1a_5p/P2a1_5p/P2a2_5p 89174.7542403281   -0.180701783114052
## Hsa-Let-7-P1b_5p                 3359.93816553835   -0.294623990307531
## Hsa-Let-7-P1c_5p                 3992.69006418194     0.48926666782258
## Hsa-Let-7-P2a3_5p                52000.6252125236   -0.206937259890988
## Hsa-Let-7-P2b1_5p                9751.60360980849 -0.00482780603867569
## ...                                           ...                  ...
## Hsa-Mir-95-P2_3p                 390.216333625945   0.0426916660211096
## Hsa-Mir-95-P3_5p                 3.95328293251241    0.678957110901101
## Hsa-Mir-96-P1_5p                 202.656949570096   -0.464482054965902
## Hsa-Mir-96-P2_5p                 27209.4573430933   -0.221824399033929
## Hsa-Mir-96-P3_5p                 3344.18853507094   0.0672687256455847
##                                               lfcSE               stat
##                                           <numeric>          <numeric>
## Hsa-Let-7-P1a_5p/P2a1_5p/P2a2_5p 0.0639958634974693  -2.81887235438572
## Hsa-Let-7-P1b_5p                  0.106490720796663  -2.76160115635425
## Hsa-Let-7-P1c_5p                  0.149925843445819   3.33812788609782
## Hsa-Let-7-P2a3_5p                0.0563710904824311  -3.67052297626806
## Hsa-Let-7-P2b1_5p                0.0617282834747614 -0.070874324259544
## ...                                             ...                ...
## Hsa-Mir-95-P2_3p                 0.0918217873522948  0.437722915855047
## Hsa-Mir-95-P3_5p                  0.170411797990612   3.95560185906774
## Hsa-Mir-96-P1_5p                  0.152114644217898  -3.14528253438741
## Hsa-Mir-96-P2_5p                   0.12892939045268  -1.79236001772164
## Hsa-Mir-96-P3_5p                  0.164329004650402  0.314884459867911
##                                                pvalue                 padj
##                                             <numeric>            <numeric>
## Hsa-Let-7-P1a_5p/P2a1_5p/P2a2_5p  0.00481926786262815   0.0172690431744175
## Hsa-Let-7-P1b_5p                  0.00575186955495521   0.0200538154753844
## Hsa-Let-7-P1c_5p                  0.00084344919760993  0.00370925953948912
## Hsa-Let-7-P2a3_5p                0.000242054709304128  0.00121656068182724
## Hsa-Let-7-P2b1_5p                   0.943497778247006     0.97368970715091
## ...                                               ...                  ...
## Hsa-Mir-95-P2_3p                    0.661587155561431    0.823261187145574
## Hsa-Mir-95-P3_5p                 7.63422117073344e-05 0.000440961730309529
## Hsa-Mir-96-P1_5p                  0.00165926501866658  0.00690468346477385
## Hsa-Mir-96-P2_5p                   0.0730753150258787     0.17139482978797
## Hsa-Mir-96-P3_5p                    0.752849381089381     0.86454810231926
```
