## Supplementary material for "A microRNA Signature of Metastatic Colorectal Cancer": Supplementary_file_4_figures.html

Supplementary\_file\_4\_figures.utf8


```
library('tidyverse')
library("DESeq2")
library("RColorBrewer")
library("ggpubr")
library("FactoMineR")
library("factoextra")
library(ggsci)
library(umap)
library(dendsort)
library(pheatmap)
library(viridis)
library(ggforce)
library(scales)
```

```
#
dds_dict <- c("ref_pCRC" = dds <- DESeqDataSetFromMatrix(countData = countdata, colData = sampleinfo, design = ~ type.tissue),
              "ref_nCR"  = dds <- DESeqDataSetFromMatrix(countData = countdata, colData = sampleinfo, design = ~ type.tissue))

dds_dict[["ref_pCRC"]]$type.tissue = factor(dds_dict[["ref_pCRC"]]$type.tissue)
dds_dict[["ref_pCRC"]]$type.tissue = relevel(dds_dict[["ref_pCRC"]]$type.tissue, "pCRC")
dds_dict[["ref_pCRC"]] <- DESeq(dds_dict[["ref_pCRC"]] , modelMatrixType = "standard")

dds_dict[["ref_nCR"]]$type.tissue = factor(dds_dict[["ref_nCR"]]$type.tissue)
dds_dict[["ref_nCR"]]$type.tissue = relevel(dds_dict[["ref_nCR"]]$type.tissue, "nCR")
dds_dict[["ref_nCR"]] <- DESeq(dds_dict[["ref_nCR"]] , modelMatrixType = "standard")

#dds <- dds[ , dds$paper %in% consensus]
##dds <- dds[ , dds$type.tissue %in% tissue_types_keep]
#dds
#dds$type.tissue <- factor(dds$type.tissue)
#dds$type.tissue <- relevel( dds$type.tissue, "pCRC")
#dds$type        <- factor( dds$type )
#dds$type        <- relevel( dds$type, 'tumor')
#dds$malignant   <- factor(dds$malignant)
#dds             <- DESeq(dds, modelMatrixType="standard")
#rlog_dds <- rlog(dds, blind=TRUE)
```

```
##
```

```
dds_vst <- varianceStabilizingTransformation(dds)
umap.data <- t(as.matrix(assay(dds_vst)))
set.seed(1)
vst_umap = umap(umap.data)
vst_umap <- as.data.frame(vst_umap$layout)
colnames(vst_umap) <- c("UMAP1", "UMAP2")
p1 <- vst_umap %>% 
  mutate(Tissue = dds_vst$type.tissue) %>%
  mutate(Type = dds_vst$type) %>%
  ggplot(aes(UMAP1, UMAP2, fill = Tissue, shape = Type)) + 
  scale_fill_manual(
    "",
    values=c(
      'nCR'   = '#3C5488E5',
      "pCRC"  = "#E64B35E5",
      "mLi"   = "#F39B7FE5",
      "nLi"   = "#4DBBD5E5",
      "mLu"   = "#00A087E5",
      "nLu"   = "#8491B4E5",
      "PM"= "#91D1C2E5")) +
#  scale_shape_manual(values=c(24,21,22)) +
  scale_shape_manual(values=c(21,22,24)) +
  geom_point(size=4) +
  ggtitle("Tissue") +
  theme_void() +
  theme(legend.position = "none",
        plot.title = element_text(hjust = 0.5))

#ggexport(p1, filename='/Users/eirikhoy/Dropbox/Fromm_miRNA_manuscript/Rmarkdown_output/Figure_1_alternative_raw_all_miRNA.pdf', width=12, height = 5.5)
p1
```

```
rpm_values <- as.matrix(t(t(counts(dds)) / colSums(counts(dds))) * 1000000)
cell_spec_rpm <- rpm_values[rownames(rpm_values) %in% names(cell_spec_dict_inv), ]

high_exp_cell_spec <- rownames(cell_spec_rpm[rowMeans(cell_spec_rpm[, dds$type.tissue == 'pCRC']) > 100  |
                                             rowMeans(cell_spec_rpm[, dds$type.tissue == 'mLi'])  > 100  |
                                             rowMeans(cell_spec_rpm[, dds$type.tissue == 'nLi'])  > 100  |
                                             rowMeans(cell_spec_rpm[, dds$type.tissue == 'mLu'])  > 100  |
                                             rowMeans(cell_spec_rpm[, dds$type.tissue == 'nLu'])  > 100  |
                                             rowMeans(cell_spec_rpm[, dds$type.tissue == 'PM'])   > 100, ])

only_cell_marker_dds <- dds[rownames(dds) %in% high_exp_cell_spec, ]
only_cell_marker_vst <- varianceStabilizingTransformation(only_cell_marker_dds)
rownames(only_cell_marker_vst) <- rownames(only_cell_marker_vst) %>% gsub("Hsa-", "", .) %>% gsub("_5p", "", .) %>% gsub("_3p", "", .) %>% gsub("/.*", '', .)
#only_cell_marker_vst <- only_cell_marker_vst[, only_cell_marker_vst$file != 'SRR5914616']
vst.pca <- PCA(t(assay(only_cell_marker_vst)), graph = FALSE)

fviz_screeplot(vst.pca, addlabels=TRUE, ylim = c(0, 50))
```

```
var_ <- get_pca_var(vst.pca)

# Contributions of variables to PC1
fviz_contrib(vst.pca, choice = 'var', axes = 1)
```

```
fviz_contrib(vst.pca, choice = 'var', axes = 2)
```

```
axes <- c(1,2)
palette <- 'jco'

p1 <- fviz_pca_ind(vst.pca,
             axes = axes,
             geom.ind = "point", # show points only (nbut not "text")
             pointshape = 21,
             pointsize = 2.5,
             fill.ind = only_cell_marker_vst$type.tissue, # color by groups
             col.ind = "black",
             #palette = palette,
             addEllipses = TRUE, ellipse.type = "confidence", ellipse.level=0.95,# 95 % confidence elipses
             legend.title = "",
             alpha.ind = 0.15
             ) + ggtitle("PCA Cell Specific miRNA by Tissue") +
                scale_fill_manual(
                              "",
                              values=c(
                                'nCR'   = '#3C5488FF',
                                "pCRC"  = "#E64B35FF",
                                "mLi"   = "#F39B7FFF",
                                "nLi"   = "#4DBBD5FF",
                                "mLu"   = "#00A087FF",
                                "nLu"   = "#8491B4FF",
                                "PM"= "#91D1C2FF")) +
                scale_color_manual(
                                "",
                                values=c(
                                'nCR' = '#00000050',
                                "pCRC"="#00000050",
                                "CLM"="#00000050",
                                "nLi"="#00000050",
                                "mLu"="#00000050",
                                "nLu"="#00000050",
                                "PM"="#00000050")) +
  theme_transparent() + theme(legend.position = 'none')

set.seed(123)
res.km <- kmeans(vst.pca$var$coord[ , c('Dim.1', 'Dim.2')], centers = 4, nstart = 25)
grp <- as.factor(res.km$cluster)


p2 <- fviz_pca_biplot(vst.pca,
             geom.ind = "point", # show points only (nbut not "text")
             pointshape = 21,
             pointsize = 2.5,
             fill.ind = only_cell_marker_vst$type.tissue, # color by groups
             col.ind = "black",
             #palette = palette,
             #addEllipses = TRUE, ellipse.type = "confidence",# Concentration ellipses
             legend.title = "",
             alpha.ind = 0.,
             select.var = list(contrib = 15),
             repel = TRUE,
             col.var = 'black',
             alpha.var = .8,
             fill.var = 'white',
             label = 'none'
             
) + ggtitle("Correlation Circle of Cell Specific miRNA") + 
                scale_fill_manual(
                              "",
                              values=c(
                                'nCR'   = '#3C5488FF',
                                "pCRC"  = "#E64B35FF",
                                "mLi"   = "#F39B7FFF",
                                "nLi"   = "#4DBBD5FF",
                                "mLu"   = "#00A087FF",
                                "nLu"   = "#8491B4FF",
                                "PM"    = "#91D1C2FF")) +
                scale_color_manual(
                                "",
                                values=c(
                                'nCR' = '#00000050',
                                "pCRC"="00000050",
                                "mLi"="00000050",
                                "nLi"="00000050",
                                "mLu"="00000050",
                                "nLu"="00000050",
                                "PM"="00000050")) +
  theme_transparent()


#p3 <- ggarrange(p1, p2, ncol = 2, common.legend = T)
p3 <- ggarrange(p1, p2, ncol = 2, legend = 'none', common.legend = F)
#ggexport(p3, filename='/Users/eirikhoy/Dropbox/Fromm_miRNA_manuscript/Rmarkdown_output/Figure_4a_raw_excl_low_exp_hide_names.pdf', width=12, height = 6)
p3
```

```
design <- tibble(
  contrast = rep('type.tissue', 9),
  group_1 = c('nCR', 'pCRC', 'nLi', 'nLu', 'nLi',  'nLu' , 'mLi' , 'mLu' , 'PM'),
  group_2 = c('pCRC', 'nCR' , 'nCR', 'nCR', 'pCRC', 'pCRC', 'pCRC', 'pCRC', 'pCRC'),
  experiment = c('nCR.vs.pCRC', 'pCRC.vs.nCR', 'nLi.vs.nCR', 'nLu.vs.nCR', 'nLi.vs.pCRC', 'nLu.vs.pCRC','mLi.vs.pCRC', 'mLu.vs.pCRC', 'PM.vs.pCRC'),
  ref        = c('ref_pCRC',    'ref_nCR',     'ref_nCR',    'ref_nCR',    'ref_pCRC',    'ref_pCRC',   'ref_pCRC',    'ref_pCRC',    'ref_pCRC'),
  exp_type = c('Normal Colon', 'Normal Colon', 'Normal vs Normal', 'Normal vs Normal', 'Normal', 'Normal', 'Metastasis', 'Metastasis', 'Metastasis'),
  norm_cont = c('NA', 'NA', 'NA', 'NA', 'NA', 'NA', 'nLi.vs.nCR', 'nLu.vs.nCR', 'nLi.vs.nCR'),
  pcrc_cont = c('NA', 'NA', 'NA', 'NA', 'NA', 'NA', 'nLi.vs.pCRC', 'nLu.vs.pCRC', 'nLi.vs.pCRC')
)
```

```
RPM_mean <- function(dds, select_tissue){
  
  rpm <- t(t(counts(dds)) / colSums(counts(dds))) * 1000000
  
  return(tibble(
    miRNA = rownames(dds), 
    RPM = rowMeans(rpm[ , dds$type.tissue == select_tissue] )))
}
```

```
VST_mean <- function(dds, select_tissue){
  vst <- varianceStabilizingTransformation(dds)
  
  return(tibble(
    miRNA = rownames(dds),
    VST = rowMeans(assay(vst[ , dds$type.tissue == select_tissue] ))
  ))
}
```

```
Res2Tibble <- function(dds, design) {
  first = TRUE
  for (n in 1:nrow(design)){
    
    print(design[n, 'experiment'])
    
    if (first == TRUE){
    coef = paste(design[n, ]$contrast, design[n, ]$group_1, 'vs', design[n, ]$group_2, sep='_')
    #res <- results( dds , contrast = c(design[n, ]$contrast, design[n, ]$group_1, design[n, ]$group_2))
    #res <- lfcShrink(dds, contrast = c(design[n, ]$contrast, design[n, ]$group_1, design[n, ]$group_2), res=res)
    res <- results( dds_dict[[design[n, ]$ref]], name = coef, alpha = p.Threshold)
    res <- lfcShrink(dds_dict[[design[n, ]$ref]], coef=coef, res=res)
    res <- as.data.frame(res)
    res$miRNA <- rownames(res)
    res <- as.tibble(res)
    res$contrast <- design[n, ]$contrast
    res$group_1 <- design[n, ]$group_1
    res$group_2 <- design[n, ]$group_2
    res$experiment <- design[n, ]$experiment
    res$exp_type <- design[n, ]$exp_type
    res$norm_cont <- design[n, ]$norm_cont
    res$pcrc_cont <- design[n, ]$pcrc_cont
    res$group_1_RPM <- RPM_mean(dds, design[n, ]$group_1)$RPM
    res$group_2_RPM <- RPM_mean(dds, design[n, ]$group_2)$RPM
    res$group_1_VST <- VST_mean(dds, design[n, ]$group_1)$VST
    res$group_2_VST <- VST_mean(dds, design[n, ]$group_2)$VST
    first = FALSE
    next
    }
  if (first == FALSE){
    coef = paste(design[n, ]$contrast, design[n, ]$group_1, 'vs', design[n, ]$group_2, sep='_')
    #res_ <- results( dds , contrast = c(design[n, ]$contrast, design[n, ]$group_1, design[n, ]$group_2))
    #res_ <- lfcShrink(dds, contrast = c(design[n, ]$contrast, design[n, ]$group_1, design[n, ]$group_2), res=res_)
    res_ <- results( dds_dict[[design[n, ]$ref]], name = coef, alpha = p.Threshold)
    res_ <- lfcShrink(dds_dict[[design[n, ]$ref]], coef=coef, res=res_)
    res_ <- as.data.frame(res_)
    res_$miRNA <- rownames(res_)
    res_ <- as.tibble(res_)
    res_$contrast <- design[n, ]$contrast
    res_$group_1 <- design[n, ]$group_1
    res_$group_2 <- design[n, ]$group_2
    res_$experiment <- design[n, ]$experiment
    res_$exp_type <- design[n, ]$exp_type
    res_$norm_cont <- design[n, ]$norm_cont
    res_$pcrc_cont <- design[n, ]$pcrc_cont
    res_$group_1_RPM <- RPM_mean(dds, design[n, ]$group_1)$RPM
    res_$group_2_RPM <- RPM_mean(dds, design[n, ]$group_2)$RPM
    res_$group_1_VST <- VST_mean(dds, design[n, ]$group_1)$VST
    res_$group_2_VST <- VST_mean(dds, design[n, ]$group_2)$VST
    res <- bind_rows(res, res_)
  }
  }
  RPMs <- tibble(
    miRNA = rownames(dds),
    rpm_pCRC = RPM_mean(dds, 'pCRC')$RPM,
    rpm_nCR = RPM_mean(dds, 'nCR')$RPM,
    rpm_nLi = RPM_mean(dds, 'nLi')$RPM,
    rpm_nLu = RPM_mean(dds, 'nLu')$RPM,
    rpm_CLM = RPM_mean(dds, 'mLi')$RPM,
    rpm_mLu = RPM_mean(dds, 'mLu')$RPM,
    rpm_PM = RPM_mean(dds, 'PM')$RPM
    )
  VSTs <- tibble(
    miRNA = rownames(dds),
    vst_pCRC = VST_mean(dds, 'pCRC')$VST,
    vst_nCR = VST_mean(dds, 'nCR')$VST,
    vst_nLi = VST_mean(dds, 'nLi')$VST,
    vst_nLu = VST_mean(dds, 'nLu')$VST,
    vst_CLM = VST_mean(dds, 'mLi')$VST,
    vst_mLu = VST_mean(dds, 'mLu')$VST,
    vst_PM = VST_mean(dds, 'PM')$VST
  )
  
  
  res <- inner_join(res, RPMs)
  res <- inner_join(res, VSTs)
  return(res)
}
```

```
# Make results tibble

res <- Res2Tibble(dds, design)%>%
  mutate(exp_thr = ifelse(group_1_RPM > rpm.Threshold | group_2_RPM > rpm.Threshold, 'yes', '')) %>%
  #mutate(signature = ifelse(padj < p.Threshold & abs(log2FoldChange) > lfc.Threshold & exp_thr == 'yes', 'yes' , ''))
  mutate(signature = case_when(
    padj < p.Threshold & log2FoldChange >  lfc.Threshold & exp_thr == 'yes' ~ 'up',
    padj < p.Threshold & log2FoldChange < -lfc.Threshold & exp_thr == 'yes' ~ 'down',
    TRUE ~ "")) %>%
  mutate(norm_adj = '') %>%
  mutate(pcrc_adj = '')
```

```
## # A tibble: 1 x 1
##   experiment 
##   <chr>      
## 1 nCR.vs.pCRC
## # A tibble: 1 x 1
##   experiment 
##   <chr>      
## 1 pCRC.vs.nCR
## # A tibble: 1 x 1
##   experiment
##   <chr>     
## 1 nLi.vs.nCR
## # A tibble: 1 x 1
##   experiment
##   <chr>     
## 1 nLu.vs.nCR
## # A tibble: 1 x 1
##   experiment 
##   <chr>      
## 1 nLi.vs.pCRC
## # A tibble: 1 x 1
##   experiment 
##   <chr>      
## 1 nLu.vs.pCRC
## # A tibble: 1 x 1
##   experiment 
##   <chr>      
## 1 mLi.vs.pCRC
## # A tibble: 1 x 1
##   experiment 
##   <chr>      
## 1 mLu.vs.pCRC
## # A tibble: 1 x 1
##   experiment
##   <chr>     
## 1 PM.vs.pCRC
```

```
for (n in 1:nrow(res)){

  if ((res[n, ]$norm_cont == 'NA') & (res[n, ]$pcrc_cont == 'NA')){ next }
  if ((res[n, ]$signature == 'up') & (res %>% filter(miRNA == res[n, ]$miRNA & experiment == res[n, ]$norm_cont))$signature == 'up'){ res[n, ]$norm_adj = 'yes' }
  if ((res[n, ]$signature == 'down') & (res %>% filter(miRNA == res[n, ]$miRNA & experiment == res[n, ]$norm_cont))$signature == 'down' ){ res[n, ]$norm_adj = 'yes' }
  if ((res[n, ]$signature == 'up') & (res %>% filter(miRNA == res[n, ]$miRNA & experiment == res[n, ]$pcrc_cont))$signature == 'up'){ res[n, ]$norm_adj = 'yes' }
  if ((res[n, ]$signature == 'down') & (res %>% filter(miRNA == res[n, ]$miRNA & experiment == res[n, ]$pcrc_cont))$signature == 'down' ){ res[n, ]$norm_adj = 'yes' }
  }

res <- res %>%
  mutate(Annotation = case_when(
    signature %in% c('up', 'down') & norm_adj != 'yes' ~ 'Signature miRNA',
    miRNA %in% names(cell_spec_dict_inv) & signature %in% c('up', 'down') & norm_adj != 'yes' ~ 'Cell Type Specific',
    #miRNA %in% blood.cell.mirna & signature %in% c('up', 'down') ~ 'High Blood Cell Expression',
    #miRNA %in% cell.marker.mccall ~ 'Cell Marker',
    norm_adj == 'yes' ~ 'Normal Control',
    TRUE ~ "Non-Signature"))
res$experiment <- factor(res$experiment, levels = c('nCR.vs.pCRC', 'pCRC.vs.nCR', 'mLi.vs.pCRC', 'mLu.vs.pCRC', 'PM.vs.pCRC', 
                                                    'nLi.vs.nCR', 'nLi.vs.pCRC', 'nLu.vs.nCR', 'nLu.vs.pCRC'))
```

```
min_zscore <- -1 # reduce range of zscores to inprove readability of heatmap
max_zscore <- 2  # reduce range of zscores to inprove readability of heatmap
heatmap_limits = c(min_zscore, max_zscore)


# make individual plots, no ggarrange
# Heatmap per tissue type plot
columns <- c("miRNA","vst_nCR","vst_pCRC","vst_nLi","vst_CLM","vst_nLu","vst_mLu","vst_PM")
#columns2 <- c('PM', 'mLu', 'CLM', 'pCRC', 'nLu', 'nLi','nCR')
columns2 <- c('nCR', 'nLi','nLu', 'pCRC', 'CLM', 'mLu', 'PM')
 to_zscore<-function(x){
   (x - mean(x))/(sd(x))
 }


tmp_res <- res %>%
  filter(exp_type == 'Metastasis', experiment=='mLi.vs.pCRC') %>%
  
  select(columns)  %>%
  dplyr::rename(pCRC = vst_pCRC , nCR = vst_nCR , nLi = vst_nLi , nLu = vst_nLu , 
         CLM = vst_CLM , mLu = vst_mLu, PM = vst_PM) %>%
  pivot_longer(cols=colnames(.)[2:length(colnames(.))], names_to = "Tissue",
               values_to="VST") %>% 
  
  group_by(miRNA) %>%
  mutate(z_score = to_zscore(VST)) %>%
  ungroup
```

```
## Note: Using an external vector in selections is ambiguous.
## ℹ Use `all_of(columns)` instead of `columns` to silence this message.
## ℹ See <https://tidyselect.r-lib.org/reference/faq-external-vector.html>.
## This message is displayed once per session.
```

```
tmp_res$Tissue <- factor(tmp_res$Tissue, levels = columns2)
  
tmp_res$z_score[tmp_res$z_score < min_zscore] = min_zscore
tmp_res$z_score[tmp_res$z_score > max_zscore] = max_zscore

p1 <- tmp_res %>% 
  filter(miRNA %in% high_exp_cell_spec) %>%
  mutate(miRNA = gsub("/.*", "", miRNA)) %>% 
  mutate(miRNA = gsub("Hsa-", "", miRNA)) %>%
  mutate(miRNA = factor(miRNA, levels=c(
    "Mir-133-P1_3p", "Mir-122_5p", "Mir-143_3p", "Mir-145_5p", "Mir-144_5p", "Mir-451_5p", "Mir-486_5p", "Mir-146-P2_5p", 
    "Mir-342_3p" , "Mir-155_5p", "Mir-150_5p","Mir-223_3p", "Mir-24-P1_3p", "Mir-128-P1_3p",  "Mir-192-P1_5p", "Mir-375_3p", "Mir-8-P2a_3p", "Mir-8-P2b_3p",
    "Mir-126_5p", "Mir-204-P1_5p", "Mir-335_5p", "Mir-7-P1_5p", "Mir-17-P1a_5p", "Mir-15-P1a_5p","Mir-15-P1b_5p"))) %>%
  ggplot(aes(Tissue, miRNA)) +
  geom_tile(aes( fill=z_score), color='black') +
  #geom_text(aes(label = round(VST, 0)), size=2.5, color='white') #+ 
  #coord_flip()# +
  #scale_fill_gradient2(low = 'blue', high='red')#+
  #scale_fill_distiller(palette = "RdGn") #+
  scale_fill_viridis(discrete=FALSE, limits = heatmap_limits) +
  #theme_test() #+
  #ggtitle(cell_type) +
  theme(axis.text.x = element_text(angle = 90, hjust = 1),
        axis.text.y = element_text(face='bold'),
        axis.title.x = element_blank(),
        axis.title.y = element_blank(),
        legend.position = 'bottom') +
  scale_x_discrete(expand = c(0,0)) +
  scale_y_discrete(expand = c(0,0)) +
  coord_fixed(ratio=1) +
  coord_flip()
```

```
## Coordinate system already present. Adding new coordinate system, which will replace the existing one.
```

```
#p1 %>% ggexport(filename=paste('/Users/eirikhoy/Dropbox/Fromm_miRNA_manuscript/Rmarkdown_output/Figure_4b_raw/combined_heatmap.pdf', sep=''), width=10, height = 4.4)
p1
```

```
# make individual plots, no ggarrange
# Volcano Plots

x_lims = c(-2.3, 2)

signature_miRNA = c("Mir-210_3p", "Mir-191_5p", "Mir-8-P1b_3p", "Mir-1307_5p", "Mir-155_5p")

p1 <- res %>% 
  mutate(miRNA = gsub("Hsa-", "", miRNA)) %>%
  #mutate(miRNA = gsub("_5p", "", miRNA)) %>%
  #mutate(miRNA = gsub("_3p", "", miRNA)) %>%
  mutate(experiment = gsub("mLi.vs.pCRC", "mLi vs pCRC", experiment)) %>%
  mutate(experiment = gsub("mLu.vs.pCRC", "mLu vs pCRC", experiment)) %>%
  mutate(experiment = gsub("PM.vs.pCRC", "PM vs pCRC", experiment)) %>%
  filter(exp_type == 'Metastasis') %>%
  mutate(Annotation = case_when(
  miRNA %in% signature_miRNA & Annotation == "Signature miRNA"           ~ 'Signature miRNA',
 # miRNA %in% names(cell_spec_dict_inv) & Annotation == "Signature miRNA" ~ 'Cell Type Specific',
  norm_adj == 'yes' | pcrc_adj == 'yes' ~ 'Normal Background',
  Annotation == "Non-Signature"         ~ "Not Significant",
  TRUE                                  ~ "Differentially Expressed")) %>%
  mutate(alpha_ = case_when(
    Annotation == 'Not Significant' ~ .2,
    TRUE ~ 1.0)) %>%
  ggplot(aes(x = log2FoldChange, y = -log(padj, 10), fill=Annotation, alpha=alpha_, size=Annotation))+#, size=dot_size)) +
  geom_point(shape=21, color='black') + 
  xlim(x_lims) +
  ylim(c(0, 30)) +
  
  scale_size_manual(values=c(
    "Signature miRNA"    = 3,
    "Normal Background"     = 1,
    "Not Significant"      = 1,
   # "Cell Type Specific" = 2,
    "Differentially Expressed" = 2
    )
    ) +
  scale_fill_manual(values=c(
#    "Cell Type Specific" = "#3C5488FF",
    "Signature miRNA"    = "#E64B35FF",
    "Normal Background"     = "#4DBBD5FF",
    "Not Significant"      = "grey",
    "Differentially Expressed"     = "black")
    ) +

  facet_wrap(~experiment, ncol = 3) +
  geom_vline(xintercept = lfc.Threshold, color='black', linetype='dashed', alpha=0.4) +
  geom_vline(xintercept = -lfc.Threshold, color='black', linetype='dashed', alpha=0.4) +
  geom_hline(yintercept = -log(p.Threshold, 10), color='black', linetype='dashed', alpha=0.4) +
  theme_minimal() + 
  scale_alpha(guide = 'none') +
  theme(legend.position = 'none',
        axis.title.x=element_blank(),
        axis.title.y=element_blank())
        #axis.text.x=element_blank(),)
  
miRNA_of_interest <- res %>%
  filter(experiment == 'mLi.vs.pCRC' | experiment == 'mLu.vs.pCRC' | experiment == 'PM.vs.pCRC') %>% 
  filter(Annotation == "Signature miRNA" | Annotation == "Cell Type Specific") %>% 
  filter(norm_adj != 'yes') %>%
  filter(pcrc_adj != 'yes') %>%
  arrange(desc(log2FoldChange))
miRNA_of_interest <- miRNA_of_interest$miRNA

signature_miRNA = c("Hsa-Mir-210_3p", "Hsa-Mir-191_5p", "Hsa-Mir-8-P1b_3p", "Hsa-Mir-1307_5p", "Hsa-Mir-155_5p")

p2 <- res %>%
  filter(miRNA %in% miRNA_of_interest, exp_type == 'Metastasis') %>%
  mutate(Annotation = case_when(
  miRNA %in% signature_miRNA & Annotation == "Signature miRNA" ~ 'Signature miRNA',
  #miRNA %in% names(cell_spec_dict_inv) & Annotation == "Signature miRNA" ~ 'Cell Type Specific',
  norm_adj == 'yes' | pcrc_adj == 'yes' ~ 'Background Expression',
  Annotation == "Non-Signature"         ~ "Not Significant",
  TRUE                                  ~ "Differentially Expressed")) %>%
  mutate(miRNA_ = gsub("Hsa-", "", miRNA)) %>%
  #mutate(miRNA_ = gsub("_5p", "", miRNA_)) %>%
  #mutate(miRNA_ = gsub("_3p", "", miRNA_)) %>%
  mutate(experiment = gsub("mLi.vs.pCRC", "mLi vs pCRC", experiment)) %>%
  mutate(experiment = gsub("mLu.vs.pCRC", "mLu vs pCRC", experiment)) %>%
  mutate(experiment = gsub("PM.vs.pCRC", "PM vs pCRC", experiment)) %>%
  ggplot(aes(x=reorder(miRNA_, log2FoldChange), y=log2FoldChange, fill=Annotation)) +
  geom_bar(position=position_dodge(), stat='identity', color='black', size=.2) + 

  scale_fill_manual(values=c(
    #"Cell Type Specific" = "#3C5488FF",
    "Signature miRNA"    = "#E64B35FF",
    "Background Expression"     = "#4DBBD5FF",
    "Not Significant"      = "grey",
    "Differentially Expressed"     = "black")
    ) +
  coord_flip() +
  theme_minimal() + 
  facet_wrap(~experiment, ncol = 3) +
  theme(legend.position = 'bottom',
    strip.background = element_blank(),
    strip.placement = "outside",
    axis.title.y=element_blank()) +
  geom_hline(yintercept = 0, color='black') +
  xlab("miRNA") +
  geom_hline(yintercept =  lfc.Threshold, color='black', linetype='dashed', alpha=0.4) +
  geom_hline(yintercept = -lfc.Threshold, color='black', linetype='dashed', alpha=0.4) +
#  theme(axis.title.x=element_blank()) +
  #theme(panel.background = element_rect(color='black'),
  #      axis.text.y = element_text(hjust=0),
  #      axis.title.x = element_text(face='bold'),
  #      axis.title.y = element_text(face='bold'),
  #      legend.position = 'none',
  #      legend.text = element_text()
  #      ) +

  #theme(axis.text.x = element_text(angle=90, face='bold'),
  #      legend.position = 'bottom',
  #      legend.direction = 'horizontal',
  #      panel.margin=unit(.05, "lines"),
  #      panel.border = element_rect(color = "black", fill = NA, size = 1), f
  #      strip.text.x = element_text(size=10, colour = 'white', face='bold'),
  #      strip.background = element_rect(fill='black')
  #      )# + 
  geom_errorbar(aes(ymin=log2FoldChange-lfcSE*qnorm(.025), ymax=log2FoldChange+lfcSE*qnorm(.025)), size=.3)+
  scale_y_continuous(limits = c(x_lims), oob=rescale_none)


p3 <- cowplot::plot_grid(p1, p2, ncol=1, align = 'v', rel_heights = c(1, 2))
```

```
## Warning: Removed 38 rows containing missing values (geom_point).
```

```
p3
```

```
ggexport(p3, filename='/Users/eirikhoy/Dropbox/Fromm_miRNA_manuscript/Rmarkdown_output/Figure_2_raw.pdf', width=10, height=10)
```

```
## file saved to /Users/eirikhoy/Dropbox/Fromm_miRNA_manuscript/Rmarkdown_output/Figure_2_raw.pdf
```

```
# make individual plots, no ggarrange
# FoldChange per tissue type plot
library(scales)
library(ggbeeswarm)
#miRNA_of_interest <- c("Mir-210_3p", "Mir-8-P1b_3p",  "Mir-1307_5p", "Mir-191-v1/v2_5p", 
#                       "Mir-10-P1a-v1/P1a", "Mir-338-P1_3p", "Mir-374-P1_3p", "Mir-425_5p",
#                       "Mir-577_5p")
miRNA_of_interest <- c("Mir-210_3p","Mir-8-P1b_3p","Mir-191_5p", "Mir-1307_5p", "Mir-155_5p")
miRNA_of_interest <- factor(miRNA_of_interest, levels=c("Mir-210_3p", "Mir-191_5p", "Mir-8-P1b_3p","Mir-1307_5p", "Mir-155_5p"))

#level_order <- c("pCRC.vs.nCR", "nCR.vs.pCRC", "CLM.vs.pCRC", "mLu.vs.pCRC", "PM.vs.pCRC", "nLi.vs.nCR", 
#                                               "nLi.vs.pCRC", "nLu.vs.nCR","nLu.vs.pCRC")

exp_type_of_interest <- c("Metastasis", "Normal", "Normal Colon")
p1 <- res %>%
  mutate(miRNA = gsub("Hsa-", "", miRNA)) %>%
  mutate(experiment = gsub("nCR.vs.pCRC", "nCR vs pCRC", experiment)) %>%
  mutate(experiment = gsub("pCRC.vs.nCR", "pCRC vs nCR", experiment)) %>%
  mutate(experiment = gsub("mLi.vs.pCRC", "mLi vs pCRC", experiment)) %>%
  mutate(experiment = gsub("mLu.vs.pCRC", "mLu vs pCRC", experiment)) %>%
  mutate(experiment = gsub("PM.vs.pCRC",  "PM vs pCRC",  experiment)) %>%
  mutate(experiment = gsub("nLi.vs.pCRC", "nLi vs pCRC", experiment)) %>%
  mutate(experiment = gsub("nLu.vs.pCRC", "nLu vs pCRC", experiment)) %>%
  filter(miRNA %in% miRNA_of_interest, exp_type %in% exp_type_of_interest) %>%
  mutate(miRNA = factor(miRNA, levels=c("Mir-210_3p", "Mir-191_5p", "Mir-8-P1b_3p","Mir-1307_5p", "Mir-155_5p"))) %>%
  #filter(experiment != 'pCRC vs nCR') %>%
  #filter(experiment != 'nCR vs pCRC') %>%
  ggplot(aes(x=factor(experiment, 
                      levels=c("nCR vs pCRC", "pCRC vs nCR", "mLi vs pCRC", "mLu vs pCRC", 
                               "PM vs pCRC", "nLi vs pCRC", "nLu vs pCRC")), 
    y=log2FoldChange, fill=experiment)) +
  geom_bar(position="dodge", stat='identity', color='black') + 
  scale_fill_manual(
    "",
    values=c(
      'pCRC vs nCR'='black',
      "nCR vs pCRC"="grey",
      "mLi vs pCRC"="#E64B34FF",
      "nLi vs pCRC"="#4DBBD5FF",
      "nLu vs pCRC"="#4DBBD57F",
      "mLu vs pCRC"="#E64B34CC",
      "PM vs pCRC" ="#E64B347F")) +

  #theme(axis.text.x = element_text(face = 'bold', size=20, angle=45), panel.spacing = unit(2, "lines")) +
  facet_wrap(~miRNA, ncol = 9) +
  theme_minimal() + 
  geom_hline(yintercept = 0) +
  geom_hline(yintercept =  lfc.Threshold, color='black', linetype='dashed', alpha=0.4) +
  geom_hline(yintercept = -lfc.Threshold, color='black', linetype='dashed', alpha=0.4) +
  geom_hline(yintercept = 0, color='black') +
  theme(axis.text.x = element_text(angle=90, hjust=1),
        axis.text.y = element_text(),
        #axis.title.y = element_blank(),
        axis.title.x = element_blank(),
        #axis.title.y = element_text("LFC"),
        legend.position = 'none',
        legend.direction = 'horizontal',
        panel.margin=unit(1, "lines")
        #panel.border = element_rect(color = "black", fill = NA, size = 1), 
        #strip.text.x = element_text(colour = 'white', face='bold'),
        #strip.background = element_rect(fill='black'),
        #axis.title = element_text(face= 'bold')
        ) + 
  labs(y = "LFC") +
  geom_errorbar(aes(ymin=log2FoldChange-lfcSE*qnorm(.025), ymax=log2FoldChange+lfcSE*qnorm(.025))) +
  scale_y_continuous(limits = c(-2, 2), oob=rescale_none)


RPM_df_maker <- function(dds, select_tissue){
  
  rpm <- t(t(counts(dds)) / colSums(counts(dds))) * 1000000
  listmiRNA <- rownames(rpm)
  rpm_tibble <- as.tibble(rpm[ , dds$type.tissue == select_tissue])
  rpm_tibble['miRNA'] <- listmiRNA
  rpm_tibble_wide <- rpm_tibble %>%
    pivot_longer(-miRNA, names_to = 'sample', values_to = 'RPM') %>%
    mutate(tissue = select_tissue)
  return(rpm_tibble_wide)
}

rpm_wide <- bind_rows(RPM_df_maker(dds, 'pCRC'), 
                      RPM_df_maker(dds, 'nCR'),
                      RPM_df_maker(dds, 'mLi'),
                      RPM_df_maker(dds, 'nLi'),
                      RPM_df_maker(dds, 'nLu'),
                      RPM_df_maker(dds, 'mLu'),
                      RPM_df_maker(dds, 'PM'))


rpm_wide['tissue'] <- factor(rpm_wide$tissue, levels=c('nCR', 'pCRC', 'mLi', 'mLu', 'PM', 'nLi', 'nLu'))

#y_lims <- 10^c(min(floor(min(log10(df$y))), 0), ceiling(max(log10(df$y))))


p2 <- rpm_wide %>%
  mutate(miRNA = gsub("Hsa-", "", miRNA)) %>%
  filter(miRNA %in% miRNA_of_interest) %>%
  mutate(miRNA = factor(miRNA, levels=c("Mir-210_3p", "Mir-191_5p", "Mir-8-P1b_3p","Mir-1307_5p", "Mir-155_5p"))) %>%
  #mutate(tissue = gsub("PM_CRC", "PM", tissue)) %>% 
  #mutate(log10_RPM = log(RPM, 10)) %>%
  ggplot(aes(x=tissue, y=RPM, fill=tissue)) +
  
  theme_minimal() + 

  facet_wrap(~miRNA, ncol = 9) +
  geom_beeswarm(shape=21) +
  stat_summary(fun=mean, geom="point", fill = 'white', color='black', shape=21, size=3) +
  geom_hline(yintercept = rpm.Threshold, color='black', linetype='dashed', alpha=0.4) +
  scale_fill_manual(
    "",
    values=c(
      'nCR'   = "grey",
      "pCRC"  = "black",
      "mLi"   = "#E64B34FF",
      "mLu"   = "#E64B34CC",
      "PM"    = "#E64B347F",
      "nLi"   = "#4DBBD5FF",
      "nLu"   = "#4DBBD57F")) +
  theme(axis.text.x = element_text(angle=90, hjust=1),
        axis.text.y = element_text(),
        axis.title.x = element_blank(),
        legend.position = 'none',
        legend.direction = 'horizontal',
        panel.spacing=unit(1, "lines")
        #panel.border = element_rect(color = "black", fill = NA, size = 1), 
        #strip.text.x = element_text(colour = 'white', face='bold'),
        #strip.background = element_rect(fill='black'),
        #axis.title = element_text(face= 'bold')
        ) +
  scale_y_log10(breaks = trans_breaks("log10", function(x) 10^x),
                       labels = comma)
p3 <- cowplot::plot_grid(p1, p2, ncol=1, align = 'v', rel_heights = c(1, 1))
#ggexport(p3, filename='/Users/eirikhoy/Dropbox/Fromm_miRNA_manuscript/Rmarkdown_output/Figure_3_raw.pdf', width=8, height = 7)

ggexport(p1, p2, ncol=1, filename='/Users/eirikhoy/Dropbox/Fromm_miRNA_manuscript/Rmarkdown_output/Figure_3_raw.pdf', width=10, height = 10)

#ggarrange(p1, p2, ncol=1, align = 'v')
p3
```

```
miRNA_of_interest <- c("Mir-210_3p","Mir-8-P1b_3p","Mir-191_5p", "Mir-1307_5p", "Mir-155_5p")
miRNA_of_interest <- factor(miRNA_of_interest, levels=c("Mir-10-P1a_5p", "Mir-191_5p", "Mir-8-P1b_3p","Mir-1307_5p", "Mir-210_3p"))

RPM_df_maker <- function(dds, select_tissue){
  
  rpm <- t(t(counts(dds)) / colSums(counts(dds))) * 1000000
  listmiRNA <- rownames(rpm)
  rpm_tibble <- as.tibble(rpm[ , dds$type.tissue == select_tissue])
  rpm_tibble['miRNA'] <- listmiRNA
  rpm_tibble_wide <- rpm_tibble %>%
    pivot_longer(-miRNA, names_to = 'sample', values_to = 'RPM') %>%
    mutate(tissue = select_tissue)
  return(rpm_tibble_wide)
}

rpm_wide <- bind_rows(RPM_df_maker(dds, 'pCRC'), 
                      RPM_df_maker(dds, 'nCR'),
                      RPM_df_maker(dds, 'mLi'),
                      RPM_df_maker(dds, 'nLi'),
                      RPM_df_maker(dds, 'nLu'),
                      RPM_df_maker(dds, 'mLu'),
                      RPM_df_maker(dds, 'PM'))

rpm_wide['tissue'] <- factor(rpm_wide$tissue, levels=c('nCR', 'pCRC', 'mLi', 'mLu', 'PM', 'nLi', 'nLu'))

#y_lims <- 10^c(min(floor(min(log10(df$y))), 0), ceiling(max(log10(df$y))))


p1 <- rpm_wide %>%
  mutate(miRNA = gsub("Hsa-", "", miRNA)) %>%
  filter(miRNA %in% miRNA_of_interest) %>%
  #mutate(tissue = gsub("PM_CRC", "PM", tissue)) %>% 
  #mutate(log10_RPM = log(RPM, 10)) %>%
  ggplot(aes(x=tissue, y=RPM, fill=tissue)) +
  
  #theme_minimal() + 

  facet_wrap(~miRNA, ncol = 1, scales = 'free') +
  geom_beeswarm(shape=24, cex=1.4) +
  stat_summary(fun=mean, geom="point", fill = 'white', color='black', shape=21, size=3) +
  geom_hline(yintercept = rpm.Threshold, color='black', linetype='dashed', alpha=0.4) +
  scale_fill_manual(
    "",
    values=c(
      'nCR'   = "grey",
      "pCRC"  = "black",
      "mLi"   = "#E64B34FF",
      "mLu"   = "#E64B34CC",
      "PM"    = "#E64B347F",
      "nLi"   = "#4DBBD5FF",
      "nLu"   = "#4DBBD57F")) +
  theme(axis.text.x = element_text(angle=90, hjust=1),
        axis.text.y = element_text(),
        legend.position = 'none',
        legend.direction = 'horizontal',
        panel.spacing=unit(.05, "lines")
        #panel.border = element_rect(color = "black", fill = NA, size = 1), 
        #strip.text.x = element_text(colour = 'white', face='bold'),
        #strip.background = element_rect(fill='black'),
        #axis.title = element_text(face= 'bold')
        ) #+
  #scale_y_log10(breaks = trans_breaks("log10", function(x) 10^x),
  #                     labels = comma)
ggexport(p1, filename='/Users/eirikhoy/Dropbox/Fromm_miRNA_manuscript/Rmarkdown_output/figs_new_neerincx/Sup_Figure_1_raw.pdf', width=10, height = 20)
```

```
## file saved to /Users/eirikhoy/Dropbox/Fromm_miRNA_manuscript/Rmarkdown_output/figs_new_neerincx/Sup_Figure_1_raw.pdf
```

```
p1
```

```
"
Define threshold for signature miRNA, including effect size, 
significance therhold, and expression size
"
```

```
#
dds_dict <- c("ref_pCRC" = dds <- DESeqDataSetFromMatrix(countData = countdata, colData = sampleinfo, design = ~ type.tissue),
              "ref_nCR"  = dds <- DESeqDataSetFromMatrix(countData = countdata, colData = sampleinfo, design = ~ type.tissue))

dds_dict[["ref_pCRC"]]$type.tissue = factor(dds_dict[["ref_pCRC"]]$type.tissue)
dds_dict[["ref_pCRC"]]$type.tissue = relevel(dds_dict[["ref_pCRC"]]$type.tissue, "pCRC")
dds_dict[["ref_pCRC"]] <- DESeq(dds_dict[["ref_pCRC"]] , modelMatrixType = "standard")

dds_dict[["ref_nCR"]]$type.tissue = factor(dds_dict[["ref_nCR"]]$type.tissue)
dds_dict[["ref_nCR"]]$type.tissue = relevel(dds_dict[["ref_nCR"]]$type.tissue, "nCR")
dds_dict[["ref_nCR"]] <- DESeq(dds_dict[["ref_nCR"]] , modelMatrixType = "standard")

#dds <- dds[ , dds$paper %in% consensus]
##dds <- dds[ , dds$type.tissue %in% tissue_types_keep]
#dds
#dds$type.tissue <- factor(dds$type.tissue)
#dds$type.tissue <- relevel( dds$type.tissue, "pCRC")
#dds$type        <- factor( dds$type )
#dds$type        <- relevel( dds$type, 'tumor')
#dds$malignant   <- factor(dds$malignant)
#dds             <- DESeq(dds, modelMatrixType="standard")
#rlog_dds <- rlog(dds, blind=TRUE)
```

```
rpm_values <- as.matrix(t(t(counts(dds)) / colSums(counts(dds))) * 1000000)
cell_spec_rpm <- rpm_values[rownames(rpm_values) %in% names(cell_spec_dict_inv), ]

high_exp_cell_spec <- rownames(cell_spec_rpm[rowMeans(cell_spec_rpm[, dds$type.tissue == 'pCRC']) > 100  |
                                             rowMeans(cell_spec_rpm[, dds$type.tissue == 'mLi'])  > 100  |
                                             rowMeans(cell_spec_rpm[, dds$type.tissue == 'nLi'])  > 100  |
                                             rowMeans(cell_spec_rpm[, dds$type.tissue == 'mLu'])  > 100  |
                                             rowMeans(cell_spec_rpm[, dds$type.tissue == 'nLu'])  > 100  |
                                             rowMeans(cell_spec_rpm[, dds$type.tissue == 'PM'])   > 100, ])

only_cell_marker_dds <- dds[rownames(dds) %in% high_exp_cell_spec, ]
only_cell_marker_vst <- varianceStabilizingTransformation(only_cell_marker_dds)
```

```
## -- note: fitType='parametric', but the dispersion trend was not well captured by the
##    function: y = a/x + b, and a local regression fit was automatically substituted.
##    specify fitType='local' or 'mean' to avoid this message next time.
```

```
# test of Mir-143_3p for pCRC and nLu versus the rest
tissue_df <- as.tibble(colData(only_cell_marker_vst)[c('filename', 'type.tissue')])


long_df <- as.data.frame(assay(only_cell_marker_vst))
long_df['miRNA'] <- rownames(long_df)
long_df <- as.tibble(long_df)
long_df <- pivot_longer(long_df, cols=colnames(long_df)[1: length(colnames(long_df)) - 1], names_to = 'filename', values_to = 'VST')
long_df <- left_join(x = long_df, y = tissue_df, by='filename')
long_df
```

```
## # A tibble: 6,700 x 4
##    miRNA          filename                                      VST type.tissue
##    <chr>          <chr>                                       <dbl> <fct>      
##  1 Hsa-Mir-122_5p COMET_0003M1_1_RPI25_ACTGAT_S31_L006_R1_001 10.8  mLi        
##  2 Hsa-Mir-122_5p COMET_0011M1_1_RPI26_ATGAGC_S33_L006_R1_001 12.2  mLi        
##  3 Hsa-Mir-122_5p COMET_0014M1_1_RPI27_ATTCCT_S35_L006_R1_001  9.78 mLi        
##  4 Hsa-Mir-122_5p COMET_0016M1_1_RPI28_CAAAAG_S37_L006_R1_001 16.2  mLi        
##  5 Hsa-Mir-122_5p COMET_0026M1_1_RPI29_CAACTA_S39_L006_R1_001  6.02 mLi        
##  6 Hsa-Mir-122_5p COMET_0027M3_1_RPI30_CACCGG_S41_L006_R1_001 14.7  mLi        
##  7 Hsa-Mir-122_5p COMET_0028M1_1_RPI31_CACGAT_S43_L006_R1_001  5.24 mLi        
##  8 Hsa-Mir-122_5p COMET_0035M1_1_RPI32_CACTCA_S45_L006_R1_001  9.59 mLi        
##  9 Hsa-Mir-122_5p COMET_0059M1_1_RPI34_CATGGC_S47_L006_R1_001  9.32 mLi        
## 10 Hsa-Mir-122_5p COMET_0128M1_S30_R1_001                      3.09 mLi        
## # … with 6,690 more rows
```

```
test_miRNA <- 'Hsa-Mir-143_3p'
groups_to_test = c( 'mLi' ='Group_2',
                    'mLu' ='Group_2',
                    'PM'  ='Group_2',
                    'pCRC'='Group_2',
                    'nLi' ='Group_2',
                    'nLu' ='Group_1', 
                    'nCR' ='Group_1' )


long_df_sub <- long_df %>% filter(miRNA == test_miRNA)
long_df_sub$group <- str_replace_all(long_df_sub$type.tissue, groups_to_test)

long_df_sub$group <- factor(long_df_sub$group)
ind.t.test <- t.test(VST ~ group, data=long_df_sub, alternative='two.sided')
ind.t.test
```

```
## 
##  Welch Two Sample t-test
## 
## data:  VST by group
## t = 2.0829, df = 39.435, p-value = 0.04379
## alternative hypothesis: true difference in means is not equal to 0
## 95 percent confidence interval:
##  0.0238408 1.6070434
## sample estimates:
## mean in group Group_1 mean in group Group_2 
##              19.09370              18.27826
```

```
test_miRNA <- 'Hsa-Mir-145_5p'
groups_to_test = c( 'mLi' ='Group_2',
                    'mLu' ='Group_2',
                    'PM'  ='Group_2',
                    'pCRC'='Group_2',
                    'nLi' ='Group_2',
                    'nLu' ='Group_1', 
                    'nCR' ='Group_1' )


long_df_sub <- long_df %>% filter(miRNA == test_miRNA)
long_df_sub$group <- str_replace_all(long_df_sub$type.tissue, groups_to_test)

long_df_sub$group <- factor(long_df_sub$group)
ind.t.test <- t.test(VST ~ group, data=long_df_sub, alternative='two.sided')
ind.t.test
```

```
## 
##  Welch Two Sample t-test
## 
## data:  VST by group
## t = 3.9098, df = 40.768, p-value = 0.0003415
## alternative hypothesis: true difference in means is not equal to 0
## 95 percent confidence interval:
##  0.9188013 2.8827437
## sample estimates:
## mean in group Group_1 mean in group Group_2 
##             10.168565              8.267792
```

```
test_miRNA <- 'Hsa-Mir-8-P2a_3p'
groups_to_test = c( 'mLi' ='Group_2',
                    'mLu' ='Group_2',
                    'PM'  ='Group_2',
                    'pCRC'='Group_2',
                    'nLi' ='Group_1',
                    'nLu' ='Group_1', 
                    'nCR' ='Group_2' )


long_df_sub <- long_df %>% filter(miRNA == test_miRNA)
long_df_sub$group <- str_replace_all(long_df_sub$type.tissue, groups_to_test)
long_df_sub$group <- factor(long_df_sub$group)
ind.t.test <- t.test(VST ~ group, data=long_df_sub, alternative='two.sided')
ind.t.test
```

```
## 
##  Welch Two Sample t-test
## 
## data:  VST by group
## t = -25.186, df = 37.636, p-value < 2.2e-16
## alternative hypothesis: true difference in means is not equal to 0
## 95 percent confidence interval:
##  -8.045591 -6.848087
## sample estimates:
## mean in group Group_1 mean in group Group_2 
##               6.83544              14.28228
```

```
test_miRNA <- 'Hsa-Mir-8-P2b_3p'
groups_to_test = c( 'mLi' ='Group_2',
                    'mLu' ='Group_2',
                    'PM'  ='Group_2',
                    'pCRC'='Group_2',
                    'nLi' ='Group_1',
                    'nLu' ='Group_1', 
                    'nCR' ='Group_2' )

long_df_sub <- long_df %>% filter(miRNA == test_miRNA)
long_df_sub$group <- str_replace_all(long_df_sub$type.tissue, groups_to_test)

long_df_sub$group <- factor(long_df_sub$group)
ind.t.test <- t.test(VST ~ group, data=long_df_sub, alternative='two.sided')
ind.t.test
```

```
## 
##  Welch Two Sample t-test
## 
## data:  VST by group
## t = -15.547, df = 31.672, p-value = 2.236e-16
## alternative hypothesis: true difference in means is not equal to 0
## 95 percent confidence interval:
##  -9.086970 -6.980901
## sample estimates:
## mean in group Group_1 mean in group Group_2 
##              3.694996             11.728932
```
