## Supplementary material for "A microRNA Signature of Metastatic Colorectal Cancer": Supplementary_file_5_comet_qpcr_analysis.html

COMET qPCR\_Validation\_Report


### COMET qPCR\_Validation\_Report

###### 03 March, 2021

Analysis of qPCR analysis of Mir-210\_3p. Mir-191\_5p and Mir-103-P1\_3p was originally selected as reference miRNAs, as they are reported to be ubiquitously expressed in all cell types [McCall, Matthew N et al. “Toward the human cellular microRNAome.” Genome research vol. 27,10 (2017): 1769-1781. doi:10.1101/gr.222067.117]. However the NGS differential expression analasis revealed that Mir-191\_5p is upregulated in the metastatic sites, therefore Mir-191\_5p was removed from the reference.

```
# Load Libraries
library(tidyverse); 
library(pastecs); 
library("ggpubr")
#library(WRS2)
library(boot); 
#library(ggm); 
library(ggplot2); 
library(Hmisc);
#library(polycor)
#library(egg)
#library(plyr)
```

```
# Load Data
qpcr_data <- read_tsv('/Users/eirikhoy/Dropbox/comet_analysis/data/qPCR_data/merged_Cq_values.tsv')
qpcr_data <- qpcr_data %>%
  filter(Cq_value < 35) # Drop samples with raw Cq value < 35
```

```
# Normalize qPCR plates on calibrator samples
plate1 <- qpcr_data %>%
  filter((Sample %in% c("COMET-0122M2-2", "COMET-0134M1-1", "BMM-0043", "BMM-0063")) &
         (PCR_plate == "05.11.19-1"))
plate1_cal <- median(plate1$Cq_value)

plate2 <- qpcr_data %>%
  filter((Sample %in% c("COMET-0122M2-2", "COMET-0134M1-1", "BMM-0043", "BMM-0063")) &
         (PCR_plate == "06.11.19-2"))
plate2_cal <- median(plate2$Cq_value)

lower_PCR_plate <-c(unique(qpcr_data$PCR_plate))[which.min(c(plate1_cal, plate2_cal))]

calibrator <- max(plate1_cal, plate2_cal) - min(plate1_cal, plate2_cal)

qpcr_data_calibrated <- qpcr_data %>%
  mutate(median_diff = calibrator) %>%
  mutate(coefficient = ifelse(PCR_plate == lower_PCR_plate, 1, 0)) %>%
  mutate(Cq_value_c = Cq_value + median_diff*coefficient)
```

```
y_max = max(drop_na(qpcr_data)$Cq_value)
y_min = min(drop_na(qpcr_data)$Cq_value)

p1 <- ggplot(qpcr_data, aes(x = PCR_plate, y = Cq_value, fill = miRNA)) + 
  geom_dotplot(aes(),
               binaxis = 'y',
               binwidth = 0.2,
               stackdir = "center") + 
  scale_y_reverse(lim = c(y_max + 1, y_min - 1)) +
  theme(axis.text.x = element_text(angle = 45, hjust = 1)) +
  ggtitle("Raw Cq values") + theme(legend.position = "none")

p2 <- ggplot(qpcr_data_calibrated, aes(x = PCR_plate, y = Cq_value_c, fill = miRNA)) + 
  geom_dotplot(aes(),
               binaxis = 'y',
               binwidth = 0.2,
               stackdir = "center") + 
  scale_y_reverse(lim = c(y_max + 1, y_min - 1)) +
  theme(axis.text.x = element_text(angle = 45, hjust = 1)) +
  ggtitle("Calibrated Cq values")

ggarrange(p1, p2, nrow=1)
```

```
# Update Cq values to between plate calibrated Cq values
qpcr_data$Cq_value <- qpcr_data_calibrated$Cq_value_c
```

Mir-191\_5p and Mir-103-P1\_3p was originally selected as reference miRNAs, as they are reported to be ubiquitously expressed in all cell types [McCall, Matthew N et al. “Toward the human cellular microRNAome.” Genome research vol. 27,10 (2017): 1769-1781. doi:10.1101/gr.222067.117].

However the NGS differential expression analasis revealed that Mir-191\_5p is upregulated in the metastatic sites, therefore Mir-191\_5p was removed from the reference.

#### pCRC vs mLi: Mir-210\_3p

```
# Discard Mir-191_5p from df

qpcr_data <- qpcr_data %>% filter(miRNA != 'Hsa-Mir-191_5p')

# Define plates and miRNA
plates       <- c('05.11.19-1', '06.11.19-2')
target_miRNA <- 'Hsa-Mir-210_3p'
reference    <- 'Hsa-Mir-103-P1_3p'

sub_qpcr <- qpcr_data %>%
  filter(!(Sample == "BMM-0043"       & PCR_plate == "06.11.19-2")) %>% # Discard duplicates
  filter(!(Sample == "COMET-0122M2-2" & PCR_plate == "06.11.19-2")) %>% # Discard duplicates
  filter(!(Sample == "BMM-0063"       & PCR_plate == "05.11.19-1")) %>% # Discard duplicates
  filter(!(Sample == "COMET-0134M1-1" & PCR_plate == "05.11.19-1")) %>% # Discard duplicates
  subset(PCR_plate %in% plates) %>%
  spread(key='miRNA', value='Cq_value')
sub_qpcr <- drop_na(sub_qpcr) # Drop na values

sub_qpcr['dCq'] <- sub_qpcr[target_miRNA] - sub_qpcr[reference]#(sub_qpcr[HK_miRNA_1] + sub_qpcr[HK_miRNA_2])/2
sub_qpcr <- gather(sub_qpcr, key="miRNA", value = "Cq_value", c(target_miRNA, reference))

sub_qpcr
```

```
## # A tibble: 44 x 6
##    Sample   PCR_plate  tissue_type      dCq miRNA          Cq_value
##    <chr>    <chr>      <chr>          <dbl> <chr>             <dbl>
##  1 BMM-0043 05.11.19-1 tumor_colorect  3.09 Hsa-Mir-210_3p     30.1
##  2 BMM-0044 05.11.19-1 tumor_colorect  7.46 Hsa-Mir-210_3p     33.6
##  3 BMM-0050 05.11.19-1 tumor_colorect  3.30 Hsa-Mir-210_3p     30.6
##  4 BMM-0053 05.11.19-1 tumor_colorect  3.81 Hsa-Mir-210_3p     31.4
##  5 BMM-0054 05.11.19-1 tumor_colorect  4.80 Hsa-Mir-210_3p     31.7
##  6 BMM-0056 05.11.19-1 tumor_colorect  5.70 Hsa-Mir-210_3p     32.3
##  7 BMM-0063 06.11.19-2 tumor_colorect  5.35 Hsa-Mir-210_3p     32.9
##  8 BMM-0129 06.11.19-2 tumor_colorect  5.65 Hsa-Mir-210_3p     33.6
##  9 BMM-0138 06.11.19-2 tumor_colorect  6.70 Hsa-Mir-210_3p     34.4
## 10 BMM-0145 06.11.19-2 tumor_colorect  6.61 Hsa-Mir-210_3p     32.6
## # … with 34 more rows
```

##### Descriptive Statistics dCq

```
stats_df <- sub_qpcr %>%
  subset(miRNA==target_miRNA) %>%
  select(tissue_type, dCq, PCR_plate)

# Descriptive Statistics
by(stats_df$dCq, stats_df$tissue_type, stat.desc, basic=TRUE,norm=TRUE)
```

```
## stats_df$tissue_type: metastasis_liver
##      nbr.val     nbr.null       nbr.na          min          max        range 
##   11.0000000    0.0000000    0.0000000    2.3350000    6.2900000    3.9550000 
##          sum       median         mean      SE.mean CI.mean.0.95          var 
##   43.8600000    4.0950000    3.9872727    0.4152575    0.9252514    1.8968268 
##      std.dev     coef.var     skewness     skew.2SE     kurtosis     kurt.2SE 
##    1.3772534    0.3454124    0.3180447    0.2406922   -1.4883739   -0.5816615 
##   normtest.W   normtest.p 
##    0.9249435    0.3619855 
## ------------------------------------------------------------ 
## stats_df$tissue_type: tumor_colorect
##      nbr.val     nbr.null       nbr.na          min          max        range 
##   11.0000000    0.0000000    0.0000000    3.0900000    7.4650000    4.3750000 
##          sum       median         mean      SE.mean CI.mean.0.95          var 
##   58.7700000    5.6500000    5.3427273    0.4366954    0.9730180    2.0977318 
##      std.dev     coef.var     skewness     skew.2SE     kurtosis     kurt.2SE 
##    1.4483549    0.2710891   -0.2602791   -0.1969760   -1.4303024   -0.5589670 
##   normtest.W   normtest.p 
##    0.9434336    0.5616640
```

##### Independent T-test between ddCq groups

```
# Independent t-test
ind.t.test <- t.test(dCq ~ tissue_type, data=stats_df, alternative = "two.sided")
ind.t.test
```

```
## 
##  Welch Two Sample t-test
## 
## data:  dCq by tissue_type
## t = -2.2493, df = 19.95, p-value = 0.03595
## alternative hypothesis: true difference in means is not equal to 0
## 95 percent confidence interval:
##  -2.61268584 -0.09822325
## sample estimates:
## mean in group metastasis_liver   mean in group tumor_colorect 
##                       3.987273                       5.342727
```

##### Boxplot ddCq

```
# Boxplot of adjusted Cq values Target miRNA, normalized to HK genes
y_min <- round(min(sub_qpcr$dCq), digits = 0)
y_max <- round(max(sub_qpcr$dCq), digits = 0)

boxplot_df <- subset(sub_qpcr, miRNA==target_miRNA)
p <- ggplot(boxplot_df, aes(tissue_type, dCq, fill=tissue_type))
p + geom_boxplot(alpha=1.0) + geom_jitter(aes(shape=PCR_plate),
                              size=1.5, position=position_jitter(0.2)) +
  ggtitle(target_miRNA) + scale_y_continuous(breaks=seq(y_min, y_max,1)) +
  scale_y_reverse()
```

```
# Plot unadjusted Cq values of both target and HK genes for metastasis and tumor

# PCR_plate
```

##### Boxplots of Raw Cq values

```
y_min <- round(min(sub_qpcr$Cq_value), digits = 0)
y_max <- round(max(sub_qpcr$Cq_value), digits = 0)

# Boxplots of Raw Cq values of both target miRNA and HK genes
p_target <- ggplot(subset(sub_qpcr, miRNA==target_miRNA), aes(tissue_type, Cq_value, fill=tissue_type)) + 
  geom_boxplot() + geom_jitter(shape=16, position=position_jitter(0.2)) +
  scale_y_reverse(lim = c(y_max, y_min)) + theme(legend.position = "none") +
  ggtitle(target_miRNA)
p_hk_1 <- ggplot(subset(sub_qpcr, miRNA==reference), aes(tissue_type, Cq_value, fill=tissue_type)) +
  geom_boxplot() + geom_jitter(shape=16, position=position_jitter(0.2)) +
  scale_y_reverse(lim = c(y_max, y_min)) + theme(legend.position = "none") +
  ggtitle(reference)
ggarrange(p_target, p_hk_1, ncol=2, nrow=1)
```

```
qpcr_spike_in <- read_tsv('/Users/eirikhoy/Dropbox/comet_analysis/data/qPCR_spike-in/merged_Cq_values.tsv')
```

##### RNA Spike-in UniSp2, UniSp4, UniSp5 results

```
# Boxplots of Raw Cq values of both target miRNA and HK genes
y_max = max(drop_na(qpcr_spike_in)$Cq_value)
y_min = min(drop_na(qpcr_spike_in)$Cq_value)

UniSp2_mean <- mean(drop_na(subset(qpcr_spike_in, miRNA == "UniSp2"))$Cq_value)
UniSp4_mean <- mean(drop_na(subset(qpcr_spike_in, miRNA == "UniSp4"))$Cq_value)

p <- ggplot(qpcr_spike_in, aes(x = Sample, y = Cq_value, fill = miRNA)) + 
  geom_dotplot(aes(),
               binaxis = 'y',
               binwidth = 0.4,
               stackdir = "center") + 
  scale_y_reverse(lim = c(y_max + 1, y_min - 1)) +
  theme(axis.text.x = element_text(angle = 45, hjust = 1)) +
  geom_hline(yintercept=UniSp2_mean) +
  geom_hline(yintercept=UniSp4_mean)

p
```
